# Cheetahs Combine Head Pitch Stabilisation with Prospective Yaw Orientation during High-speed Pursuit

**DOI:** 10.64898/2026.09.03.748592

**Authors:** Shengyang Zhuang, Kamryn Norton, Jiahe Yu, Dimitrios Kanoulas, Amir Patel

**Author notes:** Corresponding authors Corresponding author: Shengyang Zhuang, Amir Patel.

## Abstract

Predators pursuing agile prey must coordinate locomotion with rapidly changing visual information, yet how they control head orientation during high-speed terrestrial pursuit remains poorly understood. We reconstructed three-dimensional head, torso, and lure target trajectories from synchronised multi-camera recordings of freely running cheetahs (*Acinonyx jubatus*) using model-based markerless reconstruction, and propagated reconstruction uncertainty through Monte Carlo analysis. We reconstructed 1,708 frames from 18 pursuits. Across high-speed pursuit frames, head-on-torso counter-rotation largely cancelled locomotor torso oscillations, maintaining precise head-pitch alignment with the moving target. Head yaw varied with pursuit geometry: frontal pursuits favoured current-target alignment, whereas lateral and close pass-by pursuits showed ahead-of-target orientations. During lateral pursuit, head and torso maintained distinct headings while the cheetahs continued to close range. Several look-ahead rules accounted for lateral head-yaw orientation, including constant angular lead, velocity extrapolation and spatial lead. Together, these findings reveal complementary head orientation strategies during pursuit: robust sagittal stabilisation and prospective head-yaw orientation keep the head target-referenced while the torso generates a closing trajectory towards intersection.

## 1 Introduction

Accurate sensing during locomotion requires animals to distinguish motion in the external world from motion generated by their own bodies. Translations and rotations of the supporting body displace the sensory apparatus and generate self-induced retinal motion, potentially degrading information about behaviourally relevant objects [1, 2]. The cheetah (*Acinonyx jubatus*), the fastest terrestrial animal in the world, presents an especially demanding instance of this problem. The long strides, extensive spinal flexion–extension and rapid manoeuvres that enable high-speed pursuit also produce large and rapidly changing movements of the torso [3–5]. At the same time, successful interception requires the cheetah to maintain an informative visual reference to prey whose position and direction can change continuously.

Animals can reduce locomotion-induced visual disturbance through coordinated movements of the eyes, head and neck [6]. Head control varies with locomotor context [7]: monkeys and horses compensate for trunk motion [8–10], running lizards attenuate trunk yaw oscillations [11, 12], and pigeons combine rotational stabilisation with head saccades [13–15]. Recent work shows that dwarf reed snakes stabilise the head through coordinated body movements during rapid rolling [16]. Mechanistic studies identify passive attenuation of wingbeat-induced motion by a swan-inspired neck model [17] and complementary head–body feedback that broadens gaze-stabilisation bandwidth in flies [18]. In hawkmoths, measured compensatory head movements expand the speed range encoded by a model motion-vision system [19]. Related studies span echolocating bats [20, 21], other flying insects [22, 23] and harbour seals [24]. However, how a terrestrial predator stabilises its head while simultaneously tracking a moving target during high-speed pursuit remains largely unexplored. Addressing this question in cheetahs is particularly challenging because their sparse populations and large home ranges make studies in the wild difficult. Their ongoing decline also makes quantitative knowledge of their hunting behaviour particularly valuable [5, 25].

Although cheetah limb dynamics, spinal motion, gait mechanics and whole-body kinematics have received considerable attention [4, 5, 26–31], little is known about how the head-borne sensory platform is controlled during high-speed pursuit. Modern cheetahs possess an unusually enlarged and specialised vestibular system relative to other felids, which has been proposed to enhance sensitivity to head motion and support postural and visual stability during rapid locomotion [32]. However, this anatomical hypothesis has not been tested through three-dimensional (3D) measurements of head and torso motion during high-speed pursuit. It therefore remains unknown how much cheetahs stabilise head orientation against torso motion, how such stabilisation is produced, or whether these demands are expressed differently across rotational axes. Resolving these questions requires precise simultaneous reconstruction of the head, torso and target under field conditions, where marker attachment is impractical, and occlusion, distance and variable viewpoints make accurate measurement challenging.

Here, we used synchronised multiview high-speed video recordings of freely running cheetahs and model-based markerless reconstruction to quantify the 3D orientation of their head and torso as they pursued a motor-driven lure on a track, with reconstruction uncertainty propagated through Monte Carlo analysis. We examined pitch and yaw separately to determine how the head combines locomotor disturbance rejection with target-directed orientation. In pitch, we asked whether the head remained aligned with target elevation despite torso oscillations and whether head-on-torso counter-rotation offset concurrent torso motion. In yaw, we tested how orientation relative to the target varied with pursuit geometry and whether a fore-cast of target motion explained head direction. We show that head pitch remained comparatively stable and closely aligned with target elevation through near-unit counter-rotation of torso-pitch motion. Head yaw shifted from approximate target alignment in frontal pursuit to positive angular leads in lateral and close pass-by pursuit. Together, we show how cheetahs overcome the sensory challenge by controlling the head as a stabilised and prospectively oriented sensory platform.

## 2 Material and methods

### (a) Dataset and analytical cohorts

We conducted a secondary analysis of the original multi-camera recordings from the AcinoSet dataset [26], rather than of its published baseline 3D trajectories. We re-synchronised the camera streams post hoc for the present study. The recordings were collected during lure-pursuit enrichment exercises at the Ann van Dyk Cheetah Centre and Cheetah Outreach in South Africa in 2017 and 2019 (figure 1a). The present study included 6 cheetahs. Pursuits were filmed using calibrated GoPro cameras (90 or 120 frames s^−1^), with four to six usable camera views per sequence. No new animal handling, experiments or video collection was undertaken for this study. Pursuit recordings were selected when the head, torso and lure were simultaneously visible from sufficient calibrated views to support reliable reconstruction. The final analysis included 1,708 reconstructed frames from 18 pursuits. Frame-level observations were analysed within each sequence, with aggregation at the sequence or animal level used where appropriate. Details on analysis cohorts and frame counts are listed in table S1. To account for speed fluctuations within each stride, we averaged speed over complete limb cycles [5]. Across the full available sequences, stride-averaged horizontal trunk-centre speeds were 5.3–15.8 m s^−1^ (median 10.6 m s^−1^, 41 strides), calculated as path length divided by the duration of each complete forelimb cycle (supplementary note S6) using the AcinoSet full-trajectory estimation results [26].

**Figure 1:**
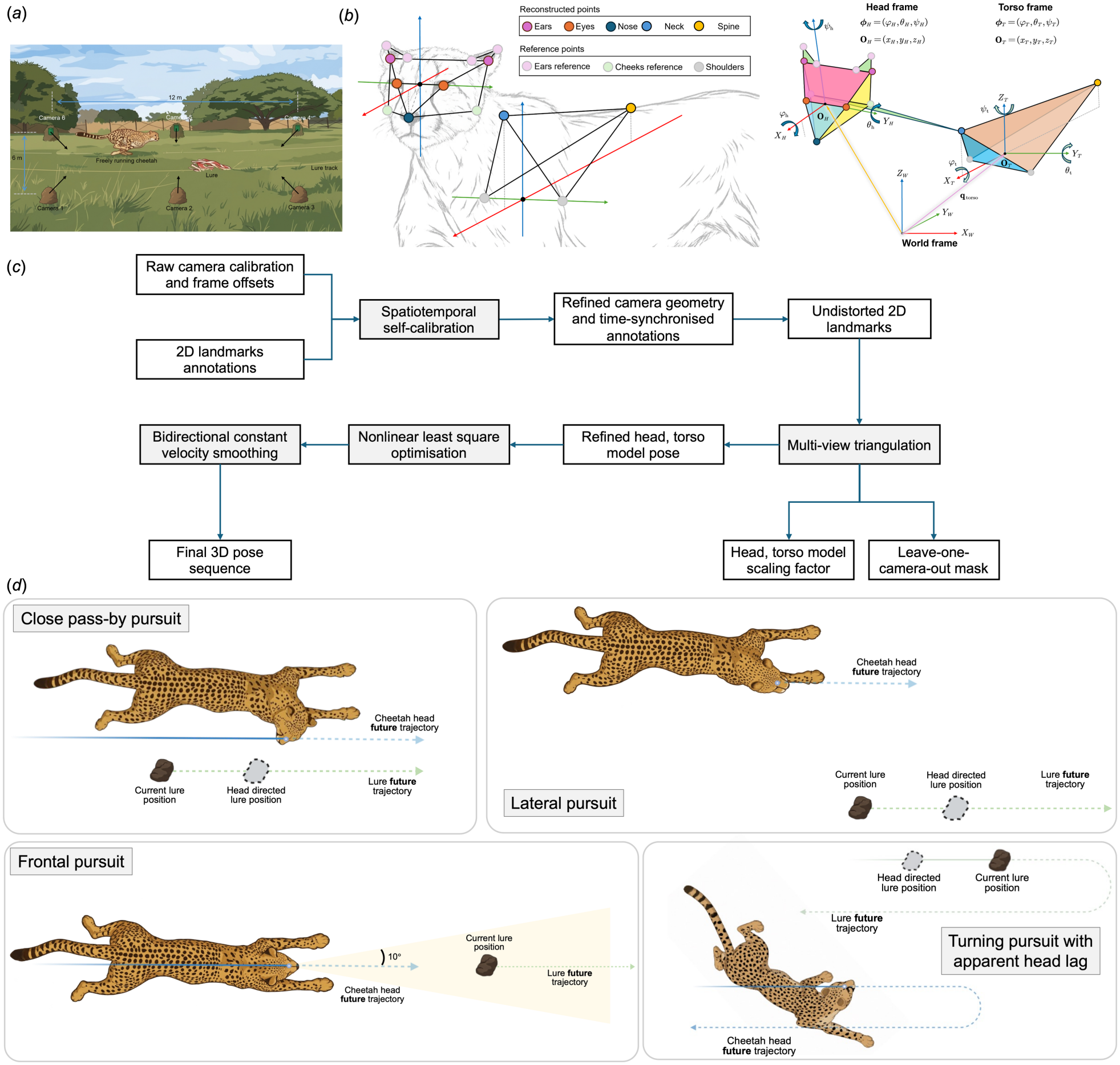
Multicamera recording setup, reconstruction pipeline and pursuit geometries. **(a)** AcinoSet pursuit field setup, showing the lure track, cheetah running area, and placement of calibrated GoPro cameras. **(b)** Reconstructed anatomical landmarks and model template reference points were used to establish animal-specific head and torso geometry. **(c)** Overview of the CRBR reconstruction pipeline. **(d)** Schematic close pass-by, lateral, frontal and turning pursuit geometries. Blue and green paths indicate head and lure trajectories, respectively; dashed extensions indicate future paths. Grey lure symbols illustrate positions towards which the head could be directed. The frontal example shows the 10^°^ alignment tolerance. The turning example illustrates how orientation towards a possible post-turn target position could appear lagged relative to instantaneous target travel. It is a conceptual example of path-dependent orientation.

### (b) 3D reconstruction of head, torso and lure target kinematics

Head and torso pose were represented by two anatomically scaled rigid-body models. The head model was defined by the left and right eyes, nose and left and right ears, with its origin at the inter-eye midpoint, whereas the torso model was defined by the neck base and spine landmarks, with its origin at the midpoint between shoulder reference points (figure 1b). Model dimensions were sourced from the AcinoSet study [26], except for the ear dimensions, which were estimated by multiview triangulation in the present study (table S2). For each analysed frame, two-dimensional (2D) image coordinates for these seven anatomical landmarks and the lure target were obtained using semi-automated landmark annotation in Blender (version 5.0.1), with example 2D poses shown in supplementary movie S2.

We developed the *Consensus-weighted Robust nonlinear least-squares optimisation with Bidirectional constant-velocity smoothing for Reconstruction (CRBR)* pipeline (figure 1c, supplementary movie S1). AcinoSet used independently recording GoPro cameras with LED-based synchronisation rather than hardware-synchronised image acquisition [26]. During high-speed pursuit, even small timing mismatches cause different views to capture different animal poses, compromising multiview reconstruction. The first stage of CRBR therefore used spatiotemporal self-calibration to refine camera timing and extrinsics from the landmark annotations (algorithm S1; table S3). Image coordinates were then corrected for lens distortion using the refined camera calibration. For each frame, visible landmarks were triangulated from the calibrated camera views. An unreliable camera view was downweighted or excluded when its held-out reprojection residual was inconsistent with the remaining cameras (figure S1, algorithm S2). Lure target position was reconstructed independently by direct multiview triangulation of the labelled lure point. Details are provided in supplementary notes S1 and S2. To account for differences in cheetah size, the head and torso templates were scaled separately for each animal using robust across-frame median estimates from multiview triangulation. The scaled models were then fitted independently in each frame to the retained 2D image observations using robust nonlinear least squares to minimise landmark-uncertainty-normalised reprojection residuals across retained cameras (figure S2, algorithm S3). Further details of the rigid-body model, scale estimation and reprojection-residual handling are provided in supplementary note S3. The framewise estimates were subsequently refined using an offline, bidirectional constant-velocity smoother on both rotation and translation (algorithm S4, supplementary note S4). Supplementary movie S3 illustrates a reconstructed pursuit.

### (c) Reconstruction diagnostics and uncertainty propagation

3D reconstruction quality was assessed from image-plane reprojection residuals (figure S8, S9); diagnostics before self-calibration are shown separately in figure S7. Across the retained camera views, mean reprojection root-mean-square error (RMSE) was 5.13 pixels across the final 1,708 analysed frames (table S4). Because ground-truth 3D pose was unavailable, this measure characterises agreement with the multiview observations rather than absolute 3D accuracy. We therefore propagated Monte Carlo uncertainty in landmark annotation, lure position and anatomical geometry through 500 joint reconstructions per pursuit, repeating the CRBR pose-fitting and smoothing stages with camera calibration and synchronisation offsets held fixed. We perturbed head, torso and lure annotations with landmark-specific image-plane Gaussian noise (radial RMS 2.5–3.5 pixels) and varied scaled template dimensions according to their estimated anatomical uncertainty (table S5, S6). Perturbation noise magnitudes were chosen to approximate reported human annotation variability [33] and increased for less visually distinct landmarks. Each Monte Carlo draw successfully generated a complete joint head, torso and lure trajectory. Continuous reconstruction uncertainty was summarised by the 2.5th and 97.5th percentiles. These intervals and probabilities quantify sensitivity to the specified annotation and reconstruction uncertainties. They are not population-level confidence intervals and do not treat frames as independent biological replicates. Full details are provided in supplementary note S5.

### (d) Kinematic definitions and statistical analyses

To characterise head stabilisation, we analysed pitch and yaw separately. Kinematic variables were computed directly from the reconstructed head, torso and lure target trajectories, with head and torso forward directions defined by their local anatomical **X**_*H*_ and **X**_*T*_ axes, respectively. World-frame head and torso pitch were denoted by *θ*_*H*,pitch_ and *θ*_*T*,pitch_, and the elevations of the corresponding head-to-target and torso-to-target lines of sight by *λ*_*H*,pitch_ and *λ*_*T*,pitch_. Head–target pitch error *α*(*t*), torso–target pitch error *β*(*t*) and head-on-torso pitch *q* (*t*) were therefore defined as (figure 2a):

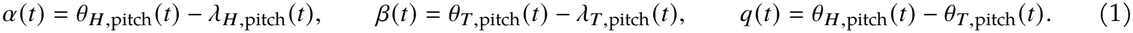

**Figure 2:**
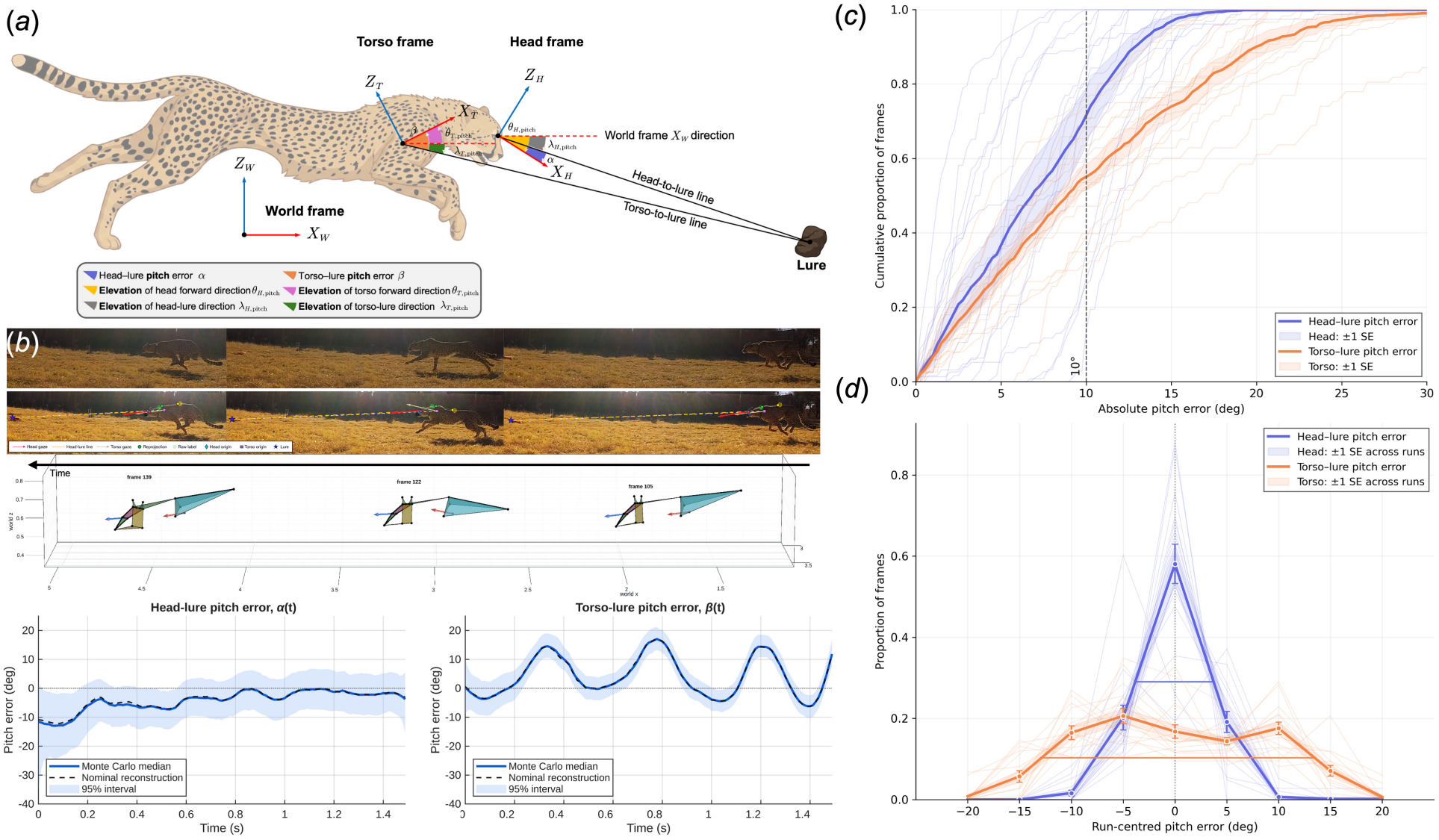
Head pitch is tightly aligned with the pursuit target. **(a)** Definition of pitch coordinates and errors. Head–target pitch error *α* and torso–target pitch error *β* are the elevation differences between each segment’s forward direction and its line of sight to the target. **(b)** Representative reconstructed pursuit sequence (phantomTopRun1_20170903, frames 105, 122 and 139, camera 5 view). Top, synchronised video frames; middle, reconstructed head and torso directions in world coordinates; bottom, the corresponding head–target and torso–target pitch-error trajectories. Solid blue lines show the median across 500 Monte Carlo reconstructions, dashed black lines show the nominal reconstruction, and shaded regions show the 95% reconstruction intervals. **(c)** Cumulative distributions of absolute pitch error across 1,590 pursuit frames. Thick lines show the pointwise means of the per-sequence distributions with each pursuit weighted equally (run-equal mean), shading shows SE across runs, and faint lines show individual runs. **(d)** Distributions of signed pitch error after centring each run on its own mean error. This removes run-specific offsets and visualises within-run precision. Distributions use 5^°^ bins. Thick lines show run-equal mean distributions, faint lines show individual runs, and shading and error bars show ±SE across runs. Horizontal bars indicate FWHM.

Values of *α* and *β* near zero indicate alignment of the relevant anatomical forward axis with the target in the pitch plane, whereas *q* describes the elevation of the head relative to the torso. Similarly, for yaw, head–target yaw error *γ* (*t*), torso–target yaw error *δ*(*t*), and head-on-torso yaw *η*(*t*) were defined as (figure 4a)

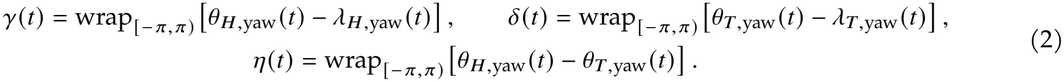

where wrap_[−*π,π*)_ (·) denotes angular wrapping to [−*π, π*).

#### Pitch alignment and torso-to-head transmission

Pitch analyses included 16 pursuits (11 frontal and 5 lateral pursuits; 1,590 frames from 5 cheetahs). The two close pass-by pursuits were excluded because the cheetah’s head was directed away from the nearby lure during the pass-by. The lure therefore did not provide a consistent reference for assessing target-directed pitch stabilisation in these sequences. Both pursuits were retained in the yaw analyses.

Within each pursuit, pitch-alignment accuracy was quantified from empirical cumulative distribution functions (ECDF) of absolute head–target and torso–target errors, |*α*| and |*β*|, and from the proportions of valid frames within 10° and 15° of the target direction. These distributions and proportions were averaged equally across pursuits, irrespective of frame count (run-equal). We also calculated the within-pursuit standard deviations (SDs) of world-frame head and torso pitch, summarised as run-equal means ± SE across pursuits. To assess within-pursuit precision independently of pointing offsets, signed *α* and *β* trajectories were centred on their within-pursuit means, converted to pursuit-level distributions and averaged equally across pursuits. Full width at half maximum (FWHM) was calculated to quantify the widths of the *α* and *β* distributions. For the animal-equal sensitivity analysis, pursuit-level distributions were averaged within animals before the animal means were weighted equally (animal-equal).

To quantify torso-to-head transmission and determine whether head-on-torso counter-rotation accounted for the attenuation of torso-pitch motion, we calculated changes over 50-ms intervals. Within each pursuit, we fitted the model

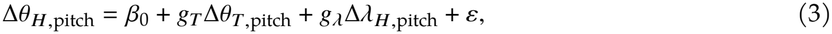

where *g*_*T*_ is the partial torso-to-head transmission gain. Because head motion could reflect both transmission of torso motion and reorientation towards the target, we included concurrent target-elevation change Δ*λ*_*H*,pitch_ as a covariate. A gain of *g*_*T*_ = 0 indicates complete rejection of torso-pitch motion, whereas *g*_*T*_ = 1 indicates complete transmission to world-frame head pitch. Because Δ*θ*_*H*,pitch_ = Δ*θ*_*T*,pitch_ + Δ*q*, the equivalent counter-rotation gain was *g*_*D*_ = 1 − *g*_*T*_, such that *g*_*D*_ = 1 indicates complete opposing counter-rotation. Co-efficients were estimated separately within each pursuit, averaged across pursuits within each animal, and then summarised as the mean and standard error (SE) across animals.

#### Yaw pursuit geometry and tracking

We analysed all 18 pursuits (1,708 frames) in three contexts illustrated in figure 1d: *frontal pursuit* (1,080 frames), with the lure approximately ahead and moving away; *lateral pursuit* (510 frames), with lure motion oblique or transverse to the head-to-target bearing; and *close pass-by pursuit* (118 frames), with the lure passing close as its bearing changed markedly. Turning pursuit, in which the cheetah followed the lure around a bend, was retained as qualitative context only (figure S17). Geometry was quantified by horizontal head–target distance and the angle between local target travel and the head-to-target bearing (figure 4d,e).

To distinguish head orientations ahead of versus behind the moving lure across different pursuit geometries, we expressed yaw error relative to the lure’s local direction of travel. We calculated the directional projection 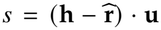, where **h**, 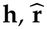 and **u** are unit horizontal vectors describing head direction, head-to-target bearing and target travel, respectively. The travel-directed yaw offset was then *γ*_∥_ = sgn (*s*) |*γ*|. Positive values indicate that a point along head direction at the current head–target distance lies ahead of the lure along its local travel direction; negative values indicate that it lies behind.

Within each of 500 joint reconstructions, aligned draws satisfied |*γ*| ≤ 10°; outside this band, positive and negative projections indicated ahead and lagged orientations, respectively. A frame was assigned a supported state when at least 95% of draws met its criterion; otherwise it was unresolved. State fractions and descriptive offset distributions were averaged equally across pursuits within each context, retaining unresolved frames in the state-prevalence denominator (figure 4e,f).

We compared head–target and torso–target absolute yaw-error ECDFs across the five lateral and two close pass-by pursuits (628 frames from four cheetahs; figure 4b), excluding frontal pursuits because target bearing and torso heading were closely aligned, providing little separation for this comparison. ECDFs were averaged equally across pursuits, with an animal-equal sensitivity analysis (figure S13); we also reported the pooled fraction of frames with |*γ*| < |*δ*|. Uncentred signed-error distributions were restricted to the five lateral pursuits (510 frames; figure 4c) to characterise sustained lateral tracking. Close pass-by pursuits were excluded from this distributional comparison because their rapidly changing target bearings mix transient reorientation with sustained tracking offsets. Pointwise standard errors were calculated across pursuit-level distributions for run-equal summaries and across animal-level mean distributions for animal-equal summaries.

To assess whether cheetahs continued to close on the lure while head and torso orientations diverged in yaw, we examined closing speed in relation to head–torso yaw separation (figure S16a). Closing speed was calculated as *c* (*t*) = −d*R*_*T*_ (*t*)/d*t*, where *R*_*T*_ is horizontal torso–target range; positive values indicate decreasing range. Estimation, filtering and plotting details are provided in supplementary note S6.

### (c) Candidate models of head-yaw orientation

To examine possible mechanisms underlying head-yaw lead, we used the same five lateral pursuits, which provided sustained tracking geometry. We compared current-target alignment with three single-parameter rules: constant angular lead, constant-velocity target extrapolation and constant spatial lead (figure 5). Target-velocity estimation required 100 ms of preceding observations; excluding these initial frames from each continuous sequence left 456 of the 510 lateral-pursuit frames for model comparison.

Let **r** = **L − H** denote horizontal head-to-target displacement, *ℬ*(**r)** = atan2 (*r*_*y*_, *r*_*x*_) its bearing, and 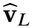 estimated target velocity. The predicted head directions were

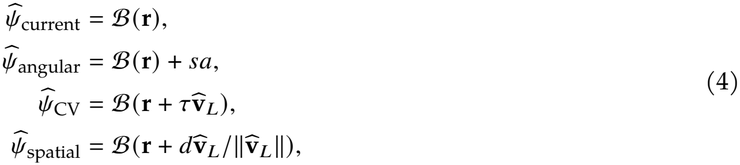

where 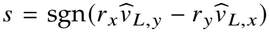 sets angular-lead direction from target motion. The fitted parameters were angular magnitude *a* (0–180°, 0.5° steps), horizon *τ* (0–400 ms, 5-ms steps) and distance *d* (0–5 m, 0.025-m steps). Target velocity was the endpoint derivative of a quadratic fit to the trailing 100 ms of target position. Fits restarted at gaps in accepted frames and required the full history; all models used identical frames valid across all 500 reconstruction draws.

Leave-one-run-out cross-validation selected each parameter by minimising mean wrapped angular MSE across equally weighted training pursuits, then applied it unchanged to the withheld pursuit. Overall RMSE was the square root of the mean held-out pursuit MSE. Velocity estimation, fitting and evaluation were repeated for each joint Monte Carlo draw; medians and 2.5th–97.5th percentiles summarise reconstruction uncertainty.

### Software and reproducibility

Reconstructions and pitch analyses were performed in MATLAB R2026a (version 26.1; The MathWorks, Inc., Natick, MA, USA) on Ubuntu 24.04.4 LTS. Required MATLAB add-ons were the Computer Vision Toolbox, Image Processing Toolbox, Optimization Toolbox, Signal Processing Toolbox, Curve Fitting Toolbox, and Statistics and Machine Learning Toolbox. The Parallel Computing Toolbox was used, where available, to parallelise Monte Carlo propagation. Code and supporting analysis files are available at https://github.com/rebl-ucl/CheetahHeadLab.

## 3 Results

### (a) Head-on-torso counter-rotation maintains precise target-relative head pitch

Across 1,590 reconstructed frames, head pitch was on average more closely aligned with the target than torso pitch. In a representative reconstruction, torso pitch showed pronounced, approximately sinusoidal oscillations during running, whereas head–target pitch error varied much less across the Monte Carlo reconstructions (figure 2b). The mean within-pursuit SD of world-frame head pitch was 2.79 ± 0.19°, compared with 8.36 ± 0.39° for torso pitch (run-equal mean ± SE across 16 pursuits). Across these pursuits, 71.5 ± 7.5% of frames had head–target error within 10°, compared with 55.0 ± 3.5% for torso–target error (run-equal mean SE; figure 2c). At 15°, the corresponding proportions were 96.8 ± 1.5% for the head and 74.0 ± 3.7% for the torso.

After within-pursuit centring, the head–target error distribution was narrower than the torso–target distribution (FWHM 7.6° versus 26.3°; figure 2d). Equal weighting of cheetahs preserved the greater accuracy and precision of head–target pitch alignment relative to torso–target alignment (figure S10).

This target-relative precision was accompanied by strong attenuation of torso-pitch motion at the head. In a representative pursuit, large changes in torso pitch were accompanied by opposing changes in head-on-torso pitch, whereas world-frame head pitch and target elevation varied much less (figure 3b).

**Figure 3:**
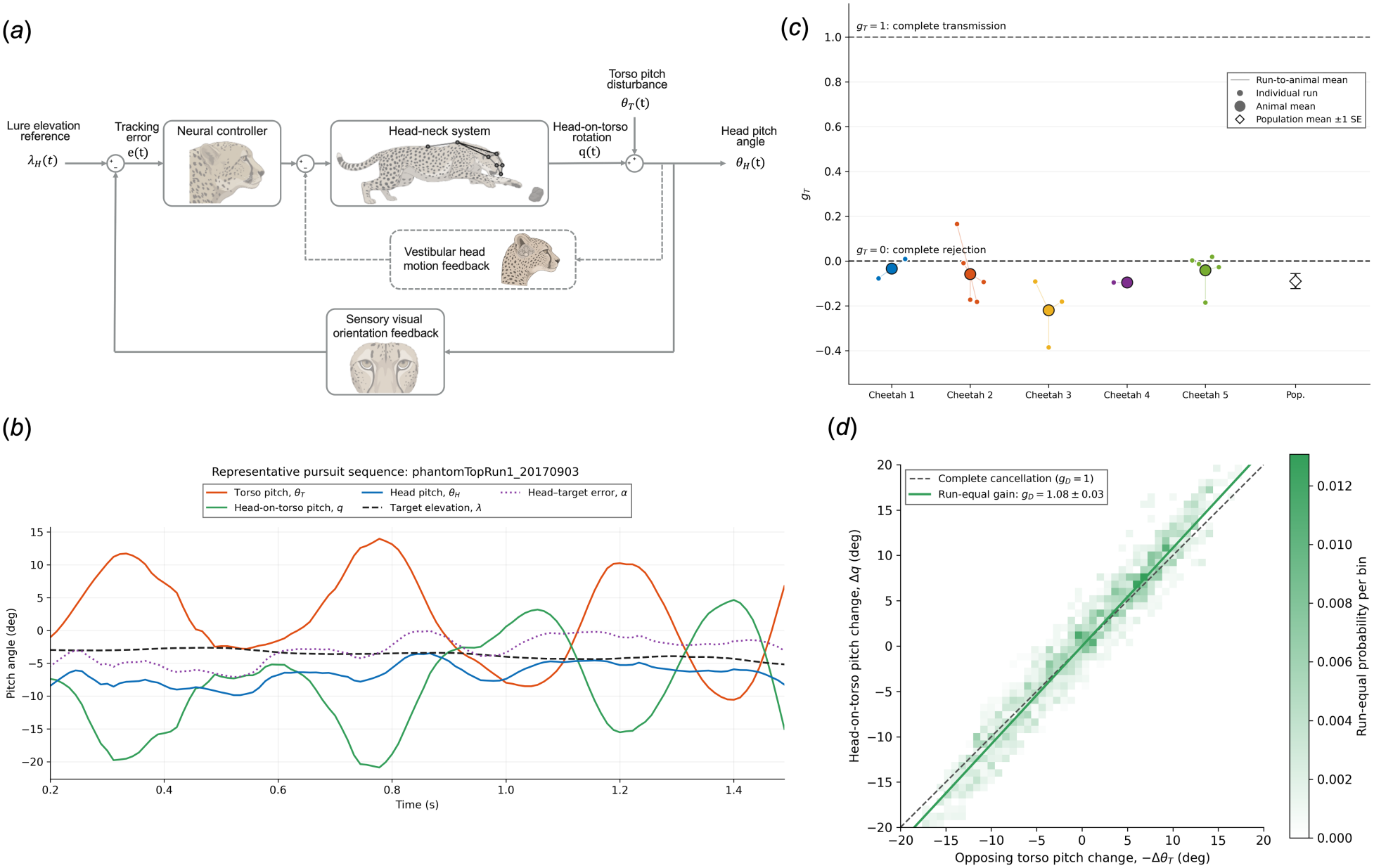
Head-on-torso counter-rotation functionally rejects torso-pitch disturbances during pursuit. **(a)** Task-level functional control representation of cheetah head-pitch stabilisation. Target elevation *λ*_*H*_ (*t*), torso pitch *θ*_*T*_ (*t*), head-on-torso pitch *q* (*t*), and world-frame head pitch *θ*_*H*_ (*t*) were obtained from reconstructed kinematics. The outer visual-feedback loop supports target tracking. The dashed inner loop represents proposed vestibulocollic feedback from head motion in space to neck control, acting to oppose head-motion disturbances. The sensory, neural and biomechanical components were not measured directly and represent a functional control hypothesis rather than a separately identified dynamical model. **(b)** Representative pursuit sequence (phantomTopRun1_20170903) showing opposing changes in torso pitch and head-on-torso pitch, together with world-frame head pitch, head–target-line elevation, and pitch error *α*. **(c)** Partial torso-to-head transmission gains estimated separately within each run while accounting for concurrent changes in target elevation. Small points denote individual runs, large coloured points denote animal means, and the open diamond denotes the animal-equal population mean with SE. A gain of zero indicates complete rejection of torso-pitch motion, whereas a gain of one indicates complete transmission to world-frame head pitch. **(d)** Run-equal raw distribution of head-on-torso pitch change, Δ*q*, against opposing torso-pitch change, Δ*θ*_*T*_, over 1,504 overlapping 50-ms intervals. Colour indicates the run-equal probability per bin. The dashed line shows the unity relationship predicted for complete cancellation. The green line has slope equal to the run-equal mean partial gain from the within-run models, which additionally included concurrent Δ*λ*_*H*_.

This attenuation was also evident from the magnitudes of the reconstructed 50-ms changes. The run-equal mean within-pursuit root-mean-square (RMS) was 2.08° for head pitch Δ*θ*_*H*,pitch_, compared with 7.81° for torso pitch Δ*θ*_*T*,pitch_ and 0.70° for target elevation Δ*λ*_*H*,pitch_. The mean within-pursuit head-to-torso RMS ratio Δ*θ*_*H*,pitch_/ Δ*θ*_*T*,pitch_ was 0.27, indicating that head-pitch changes were approximately 73% smaller than torso-pitch changes.

The animal-equal torso-to-head transmission gain was *g*_*T*_ = −0.09 0.03 SE (5 cheetahs, 1,590 frames; figure 3c). Thus, a 1° change in torso pitch was associated with only a 0.09° change in head pitch in the opposite direction after accounting for concurrent target-elevation change. Torso-pitch motion was therefore almost completely attenuated at the head. The corresponding animal-equal counter-rotation gain was *g*_*D*_ = 1.09 ± 0.03 SE, with pursuit-level estimates ranging from 0.83 to 1.39 (table S7). Head-on-torso rotation therefore changed by approximately the same amount as torso pitch in the opposing direction. The run-equal mean gain was *g*_*D*_ = 1.08 ± 0.03 SE (figure 3d). This near-unit counter-rotation provides the kinematic explanation for the low transmission of torso-pitch motion to world-frame head pitch.

Finally, we summarised these measured relationships in a conceptual task-level feedback representation (figure 3a), following the use of closed-loop block diagrams to organise sensorimotor systems [34]. In this representation, target elevation is treated as the task reference, torso pitch as an internally generated loco-motor disturbance, head-on-torso pitch as the compensatory rotation and world-frame head pitch as the task output. The diagram expresses a functional hypothesis consistent with the observed kinematic attenuation; it does not identify a specific sensory pathway, neural control law or neck–body dynamical model.

### (b) Head yaw is directed ahead of moving targets by an amount that depends on pursuit geometry

Head-yaw offsets varied with pursuit geometry, remaining small in frontal pursuit and shifting towards larger positive travel-directed values in lateral and close pass-by pursuit (figure 4d,e). In frontal pursuits, run-median target-travel angles relative to the head-to-target bearing ranged from 2.0° to 24.4°, and all 11 run-median yaw offsets lay within the ±10° alignment band. Lateral pursuits spanned travel angles of 32.5°– 105.2° and had positive median offsets of 10.7°–17.1°. The two close pass-by pursuits had median horizontal head–target distances of 0.44 and 1.07 m and travel angles of 112.0° and 162.5°; their median yaw offsets were 63.4° and 129.5°, respectively. Positive offsets denote head directions displaced towards target travel relative to the current target bearing.

**Figure 4:**
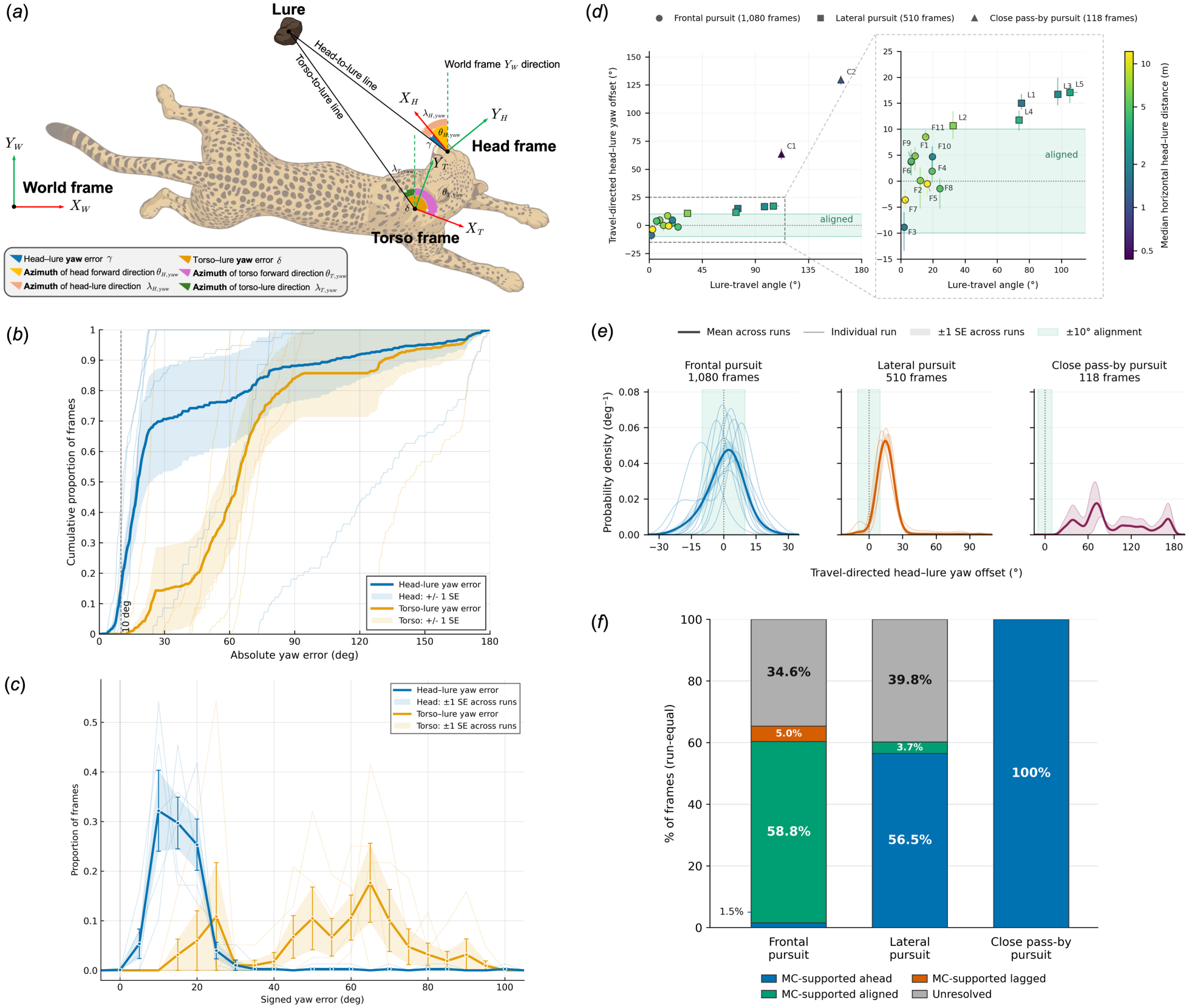
Cheetahs orient their heads ahead of moving targets with a geometry-dependent angular lead. **(a)** Yaw geometry and definitions of head–target (*γ*) and torso–target (*δ*) errors. **(b)** Absolute yaw-error ECDFs for lateral and close pass-by pursuits. Blue and orange denote head and torso; thick curves show run-equal means, thin curves individual pursuits, and envelopes ±1 SE. The dashed line marks 10^°^ alignment gate. **(c)** Uncentred signed-error distributions for lateral pursuits (5^°^ bins). Colours and line weights follow (b); shading and error bars show ±1 SE. Signs follow (a). **(d)** Travel-directed yaw offset versus target-travel angle. Positive offsets indicate ahead orientation; travel angles of 0^°^, 90^°^ and 180^°^ denote motion away from, transverse to and towards the head. Points are pursuit medians: circles, frontal; squares, lateral; triangles, close pass-by. Colour shows median horizontal head–target distance on a logarithmic scale; bars show 95% reconstruction intervals. **(e)** Smoothed travel-directed offset distributions by context. Thin curves show individual pursuits, thick curves run-equal means, and envelopes ±1 SE. Green bands in (d,e) mark 10^°^ alignment; horizontal ranges in (e) differ. **(f)** Run-equal proportions of Monte Carlo-supported ahead, aligned and lagged frames by context. Classification requires at least 95% support with a ±10^°^ alignment gate; remaining frames are unresolved.

Monte Carlo-supported classifications preserved this context dependence (figure 4f). At the ±10° alignment gate, frontal pursuits had run-equal prevalences of 58.8% aligned, 1.5% ahead and 5.0% lagged frames. In lateral pursuits, 56.5% were supported ahead, 3.7% aligned and 0% lagged. All 118 frames in the two close pass-by pursuits were supported ahead. Narrowing the alignment band to ±5° increased supported-ahead fractions to 8.5% in frontal and 81.3% in lateral pursuits. Widening it to ±15° reduced these fractions to 0.1% and 27.8%, respectively, while supported alignment increased to 86.2% and 31.5%. Both close pass-by pursuits remained 100% supported ahead at either gate (figure S14). At the primary ±10° gate, 34.6% of frontal and 39.8% of lateral frames were unresolved (run-equal means). Most unresolved lateral frames (80.4%) had at least 95% combined support for aligned or ahead orientation; unresolved frontal frames included both aligned–ahead and aligned–lagged ambiguity.

Representative image sequences, 3D reconstructions and yaw-error trajectories illustrate a lateral pursuit in which 87.0% of frames were supported ahead, with the remaining 13.0% unresolved between ahead and aligned (figure S11). A close pass-by example is shown in figure S12.

Despite these ahead-of-target offsets, head orientation remained more closely aligned with the target than torso orientation. Across 628 frames from lateral and close pass-by pursuits, head–target absolute yaw error was smaller than torso–target absolute yaw error in 92.8% of frames. The run-equal fractions within 10° of the target bearing were 14.9% for the head and 0% for the torso (figure 4b). Equal weighting of the cheetahs gave corresponding fractions of 17.1% and 0%; the animal-equal distributions are shown in figure S13.

Within the five lateral pursuits, the uncentred signed-error distributions showed head yaw concentrated at smaller positive offsets from the target, whereas torso yaw spanned a broader range of larger offsets (figure 4c).

These orientations co-occurred with continued approach to the target. Across 399 valid frames from the 4 lateral pursuits with net range closure, closing speed was positive in 87.2% of frames across a broad range of head–torso yaw separations (figure S16a). A representative top-view reconstruction shows distinct head and torso orientations along a closing trajectory (figure S16b).

## 4 Discussion

High-speed pursuit presents a fundamental sensory challenge. The long strides, flexible spine and rapid manoeuvres that enable cheetahs to pursue agile prey also generate cyclic rotations and accelerations of the body that could destabilise visual orientation. Our results reveal complementary patterns of head control in pitch and yaw. Head-on-torso counter-rotation nearly cancelled the approximately sinusoidal torso-pitch oscillations, preserving precise target-relative head pitch. Head yaw varied with pursuit geometry: frontal pursuits favoured current-target alignment, whereas lateral and close pass-by pursuits showed ahead-of-target orientation. Qualitatively similar ahead-of-prey head orientations in independent footage suggest that this behaviour also occurs during natural hunting (figure S3). Comparable predictive control has also been reported across diverse pursuit and interception systems [35–40].

These complementary, axis-specific patterns could help preserve useful visual information during pursuit. Suppressing stride-related pitch oscillations could reduce vertical image motion caused by locomotion, making target motion easier to distinguish from self-motion, while yaw orientation follows the target’s path. The potential visual benefit of head stabilisation is supported by studies in other animals. For example, experiments and modelling in hawkmoths show that head stabilisation helps the visual system detect both slow and fast self-motion [19]. To the best of our knowledge, this study provides the first precise 3D quantification of target-relative head orientation in freely pursuing cheetahs, revealing distinct patterns of stabilisation and tracking in the two axes. Our results also establish a quantitative benchmark for future studies of visual control, head–neck coordination and sensorimotor prediction in cursorial predators.

### (a) Head orientation as a proxy for visual tracking

A central assumption of this study is that head orientation provides a biologically meaningful estimate of visual orientation during pursuit. Several lines of evidence support this approximation. First, compared with other felids, cheetahs have enlarged vestibular structures associated with greater sensitivity to angular head motion, consistent with selection for stable head and gaze orientation during high-speed pursuit [32]. Second, their pronounced horizontal retinal visual streak [41] fits the terrain hypothesis: high acuity across the horizon supports panoramic monitoring in open habitats with less need for extensive eye movements [42, 43]. Third, experimental studies in felids demonstrate that gaze shifts are achieved through tightly coordinated eye-head movements, with the head contributing substantially to gaze direction during natural orienting behaviours [44–46]. During locomotion, compensatory eye movements primarily stabilise retinal images through the vestibulo-ocular reflex rather than generating large independent changes in gaze direction [47]. Together, these anatomical and behavioural observations support head orientation as a biologically meaningful proxy for visual orientation during pursuit.

However, eye-in-head movements were not recorded in these pursuits. Research shows that eye–head coordination varies with locomotor task and speed. Direct recordings in walking cats show frequent gaze shifts and fixations during inspection of the walking surface [48]. On complex surfaces, faster walking is associated with slightly less gaze shifting and more time looking a fixed distance ahead [49]. Tracking a moving target during high-speed pursuit may favour more sustained target-directed viewing. In the future, simultaneous eye recordings during cheetah pursuit would establish where the target falls on the retina and how gaze lead relates to the head-yaw lead quantified here.

### (b) Kinematic mechanisms of head pitch stabilisation

The observed near-unit counter-rotation could arise from a combination of sensory feedback, predictive motor commands and head–neck mechanics. While the present analysis identifies the kinematic strategy by which cheetahs stabilise head pitch during pursuit, it does not by itself determine the underlying sensorimotor mechanisms responsible for this coordination. Several complementary mechanisms could plausibly give rise to the observed coordination. Vestibular feedback through the vestibulocollic reflex can stabilise the head during body motion, while cervical proprioception supplies information about head–neck configuration [50, 51]. The dashed inner loop in figure 3a illustrates this proposed vestibulocollic contribution: sensed head motion in space feeds back to neck control to oppose disturbances, within the outer visual target-tracking loop. During self-generated locomotion, these signals can combine with predictive motor commands that compensate for expected stride-related disturbances [52, 53]. Head inertia and neck viscoelasticity may also attenuate body motion alongside active muscle control [54, 55]. Discriminating among these candidate mechanisms will require measurements beyond kinematics, such as neck muscle electromyography, neural recordings, or controlled mechanical and visual perturbation experiments. Nevertheless, by identifying the kinematic relationship between torso and head motion during high-speed pursuit, the present study establishes the behavioural constraint that any candidate control mechanism must satisfy. The observed near-unit counter-rotation therefore provides a kinematic benchmark for future investigations of the neural and biomechanical control of head stabilisation in cursorial predators.

### (c) Functional separation of target orientation and locomotor steering

Head–torso yaw separation during continued approach suggests that cheetahs can maintain target-oriented head movements while the body follows a closing trajectory (figure S16b). This coordination could accommodate the different demands of visual tracking and locomotion. Wild cheetah hunts involve rapid acceleration, braking and turning [5], so the body must respond to demands that extend beyond pointing towards the target. A comparable separation occurs in dragonflies, whose heads track prey while their bodies manoeuvre to align with the prey’s flight path [35]. The separation in cheetah further motivates a hypothesis in which the relationship between visual orientation and locomotor steering changes as interception approaches. During approach, orienting the head towards the target may support tracking while allowing the body to follow a different heading. Near interception, target-oriented head movements could provide a reference for subsequent torso reorientation, linking visual tracking more closely to the final steering adjustments. This hypothesis predicts a stronger association between head turns and subsequent torso turns at shorter target ranges, after accounting for preceding torso rotation and changes in target bearing. Our exploratory 100-ms analysis identified a near-range increase in one run across 99.8% of joint reconstruction draws. The short, strongly autocorrelated sequences limit inference about this proposed coordination. Longer recordings through interception, with comparable target manoeuvres at different ranges, would test whether head turns increasingly precede torso reorientation as the cheetah closes on its target.

### (d) Interpreting the mechanisms underlying prospective head yaw orientation

Orienting ahead of a moving target could help prepare subsequent tracking and capture despite delays in sensory processing and movement. In salamanders, prey-motion extrapolation accounts for accurate tongue strikes despite measured visuomotor delays [36].

To examine what could produce the observed head-yaw lead, we compared current-target alignment with constant angular lead, constant-velocity extrapolation and constant spatial lead (figure 5a). Lateral pursuits provided sustained tracking with more stable geometry than close pass-by pursuits. Across 456 evaluation frames from five lateral pursuits, leave-one-run-out prediction gave equal-run RMSEs of 19.1° for current-target alignment, 11.3° for constant angular lead, 13.2° for constant-velocity extrapolation and 12.6° for constant spatial lead (figure 5b). All three lead models reduced aggregate error, with rankings varying among pursuits.

**Figure 5:**
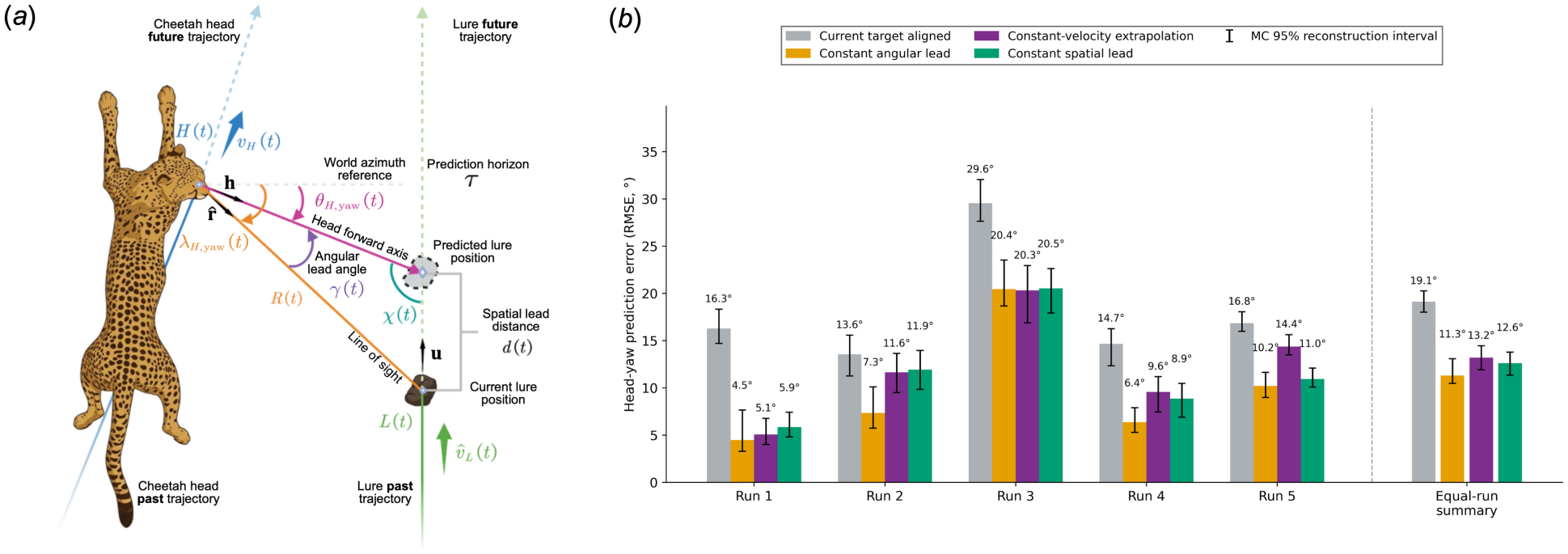
Candidate descriptions of head-yaw lead during lateral pursuit. **(a)** Geometry of angular lead *γ* (*t*), spatial lead *d* (*t*) and prediction horizon *τ*. Model predictions are bearings from the current head origin to the reference position. Dashed paths illustrate future motion schematically, and head translation is not extrapolated. **(b)** Head-yaw RMSE for current-target alignment and three lead models, evaluated by leave-one-run-out cross-validation with equal training-run weights. Bars and numerical labels show median RMSE across paired Monte Carlo reconstructions. Whiskers show 95% reconstruction intervals. The equal-run summary is the square root of the mean within-run squared error.

Constant spatial lead fixes the distance ahead along target travel, whereas constant-velocity extrapolation sets it to 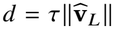, where *τ* is the prediction horizon and 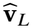 is estimated target velocity. Their predictions are similar over a narrow speed range and diverge as speed changes. The same spatial lead can produce small angular offsets in frontal pursuit and larger offsets in lateral pursuit. A fixed non-zero angular lead cannot capture the differences among frontal, lateral and close pass-by pursuits, but provides a useful approximation of head orientation during sustained lateral pursuit.

Within sustained lateral pursuit, an approximately maintained angular lead also has a useful spatial interpretation. For locally straight target motion, the horizontal geometry gives

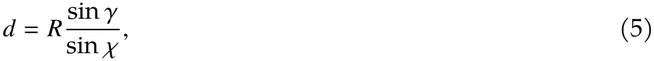

where *R* is horizontal head–target distance, *d* is distance ahead along the straight horizontal target path, *γ* is the angular lead relative to current target bearing, and *χ* is the angle from head-forward direction to local target travel (figure 5a). Measured along the reconstructed 3D target path, median spatial lead was positive in every lateral pursuit. Using 3D path length and head–target range, nominal pursuit-level medians of *d*/*R* ranged from 0.24 to 0.56 among frames with a valid forward path intersection (figure S15a). The median intersection therefore lay approximately one-quarter to just over one-half of the current head–target range ahead of the target. During intervals when both *γ* and *χ* remain approximately stable, the horizontal geometry predicts that spatial lead decreases as the cheetah approaches. The head can therefore remain directed ahead along the target’s path while its forward-path intersection moves spatially closer to the target, without requiring a change in the aiming rule.

A single pursuit-level prediction horizon cannot capture the complete frame-by-frame head trajectory. The non-zero residuals in figure 5b may therefore reflect variation in the effective prediction horizon as the cheetah combines current target velocity with acceleration, recent motion history and learned expectations from previously experienced trajectories. In the turning pursuit (figure S17), the head points behind the lure relative to its outbound direction of travel. As the lure approaches the turn, however, the head is already directed towards the path along which the lure subsequently returns (figure S17b). What appears to be lag when judged against instantaneous motion is therefore consistent with prospective orientation towards the return trajectory. Prior trajectory information could allow the cheetah to orient towards this forthcoming path before the lure reverses direction. In human interception tasks, knowing where contact will occur similarly favours directing gaze towards that location in advance [40].

However, as the recordings contain no controlled lure-speed perturbations and too few comparable changes in target motion, we cannot distinguish these mechanisms. Controlled accelerations and decelerations at comparable target ranges and travel directions would separate the speed-dependent predictions of constant-velocity extrapolation from those of constant spatial lead. Controlled changes in target direction would also test whether head orientation follows the changing path or maintains a fixed angular offset. Comparing familiar and unfamiliar routes, including unexpected reversals, would test how experience shapes prospective head orientation.

## Ethics

This study involved secondary analysis of previously collected video data from the AcinoSet dataset [26]. The original video collection was approved by the University of Cape Town Science Faculty Animal Ethics Committee. No new animal experiments, handling or video collection were performed for the present study.

## Data accessibility

The data analysed in this study were obtained from the previously published AcinoSet dataset [26] and are available at https://github.com/African-Robotics-Unit/AcinoSet. The analysis code and supporting reconstruction and analysis data are available at https://github.com/rebl-ucl/CheetahHeadLab.

## Declaration of AI use

We used AI for English proofreading, and the partial development and debugging of analysis and plotting code. We used BioRender AI to create illustration cheetah images in figure 1-5.

## Authors’ contributions

S.Z.: Conceptualization, Data curation, Methodology, Formal analysis, Investigation, Visualization, Software, Validation, Writing—original draft, Writing—review and editing. K.N.: Conceptualization, Methodology, Writing—review and editing. J.Y.: Data curation, Visualization. D.K.: Writing—review and editing. A.P.: Conceptualization, Methodology, Supervision, Project administration, Funding acquisition, Writing—review and editing.

## Acknowledgements

We thank Professor Noah Cowan for the insightful discussions of this project.

