## Supplementary Material for "Cheetahs Combine Head Pitch Stabilisation with Prospective Yaw Orientation during High-speed Pursuit"

**Shengyang Zhuang et al.**

#### Contents

This electronic supplementary material contains:

- Supplementary Figures S1–S17, providing qualitative pursuit context, camera-calibration diagnostics, reconstruction validation, pitch and yaw analyses, and sensitivity checks.
- Supplementary Notes S1–S7, describing camera synchronisation and self-calibration, camera trust, rigid-body pose fitting, bidirectional smoothing, uncertainty propagation, kinematic analyses and predictive-yaw modelling.
- Supplementary Algorithms S1–S4, detailing self-calibration, camera-trust assessment, robust framewise pose fitting and bidirectional smoothing.
- Supplementary Tables S1–S7, reporting camera self-calibration corrections and offsets, template coordinates, reconstruction quality, perturbation scales, analysis cohorts and run-level coefficients.
- Supplementary Movies S1–S3, showing overview or reconstruction method, example cheetah pose estimation and reconstruction during pursuit.

### Supplementary Figures

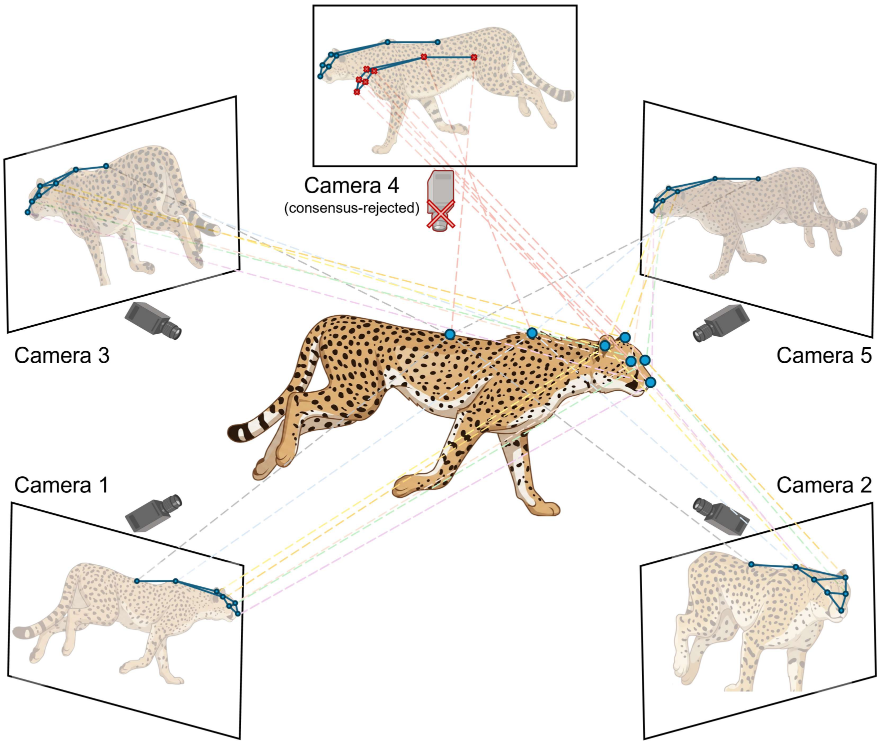


**Figure S1: Camera consensus analysis and multiview triangulation.** A leave-one-camera-out (LOO) reprojection assessment identified views inconsistent with the remaining camera consensus (illustrated by rejection of camera 4). Each landmark is triangulated individually through multiple camera views.

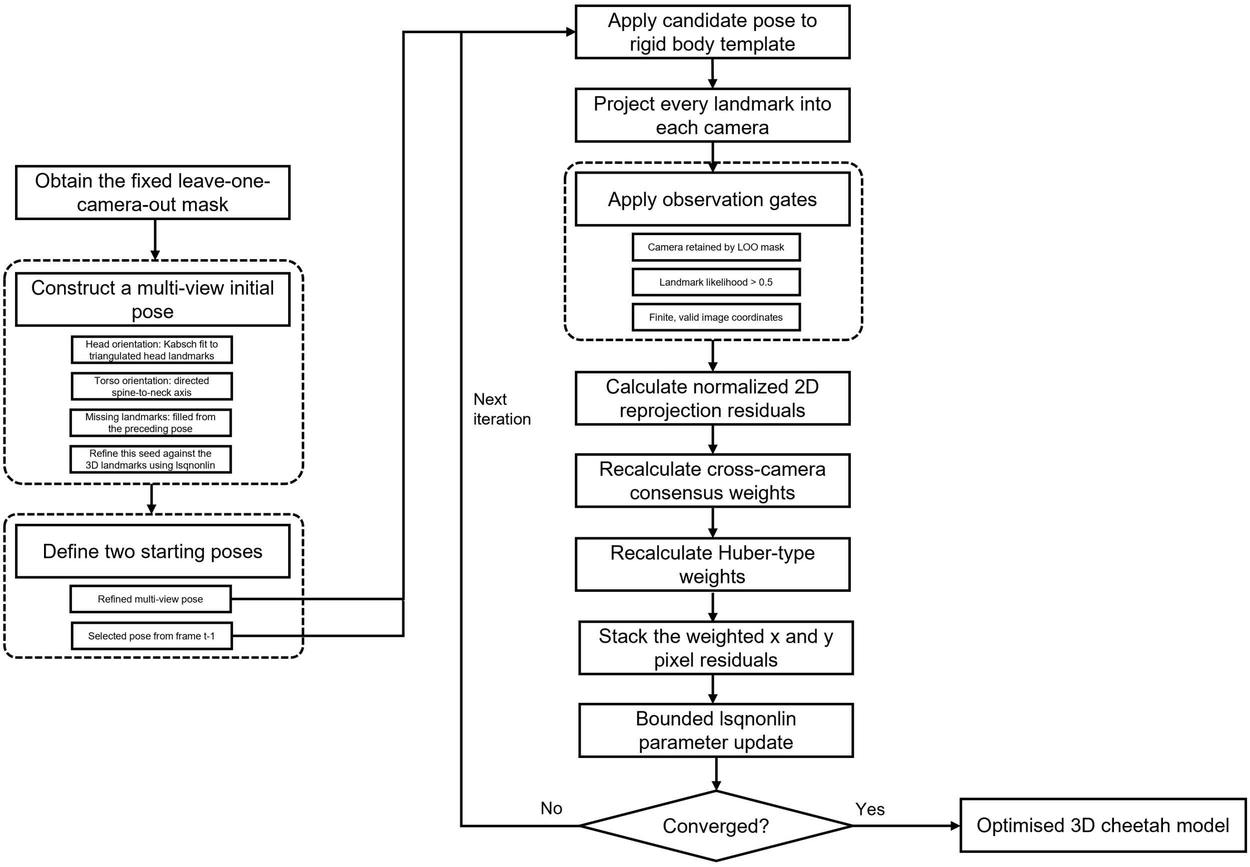


**Figure S2: Per-frame model-based reconstruction via nonlinear least square optimisation.** For each frame (t), observations from cameras excluded by the fixed LOO mask were removed, and a multiview pose was initialised from triangulated landmarks. Head orientation was estimated by least-squares rigid alignment of the head template to the triangulated head landmarks, torso orientation from the directed spine-to-neck-base axis, and missing landmarks from the preceding pose; this estimate was then refined against the reconstructed 3D landmarks. The main bounded `lsqnonlin` optimisation was run independently from this refined pose and from the solution retained at frame (t-1). At each iteration, the rigid-body template was transformed and projected into all retained cameras; valid observations were selected, normalised reprojection residuals were calculated, and cross-camera consensus and Huber-type weights were updated. The weighted image-coordinate residuals were supplied to `lsqnonlin` until convergence. The converged solution with the lower robust residual norm was retained as ($\hat{x}_t$) and propagated as one initialisation for frame (t+1).

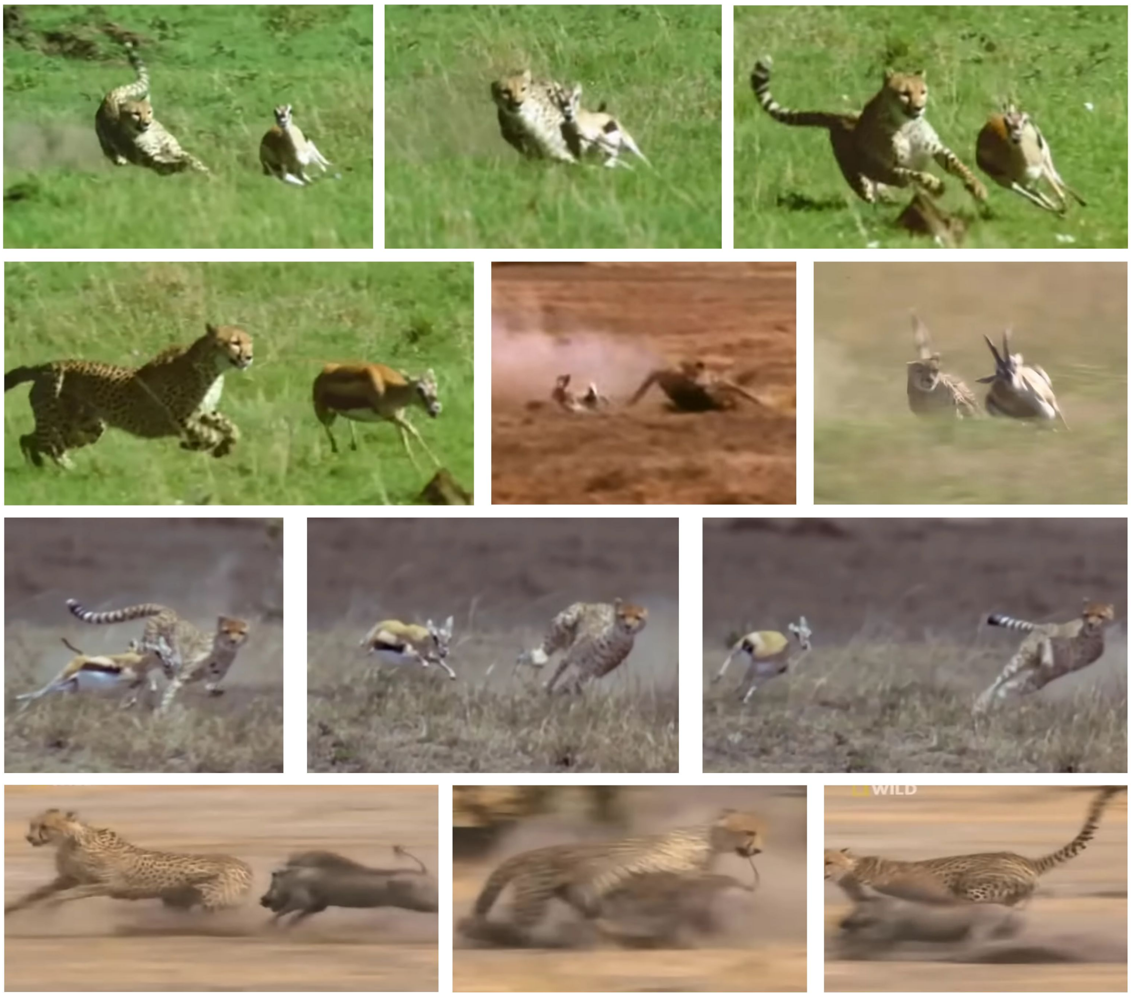

**Figure S3: Qualitative examples of ahead-of-prey head yaw orientation during wild cheetah pursuit.** Representative frames from footage of cheetahs pursuing wild prey. The cheetah's head appears oriented ahead of the prey's instantaneous position along its direction of motion, qualitatively resembling the predictive head-yaw orientation observed in our reconstructed lure pursuits. These frames are presented as qualitative behavioural context and were not included in the quantitative analysis. Frames extracted from publicly available footage on YouTube (`https://www.youtube.com/watch?v=xaV1_M2j200`; accessed 6 August 2026).

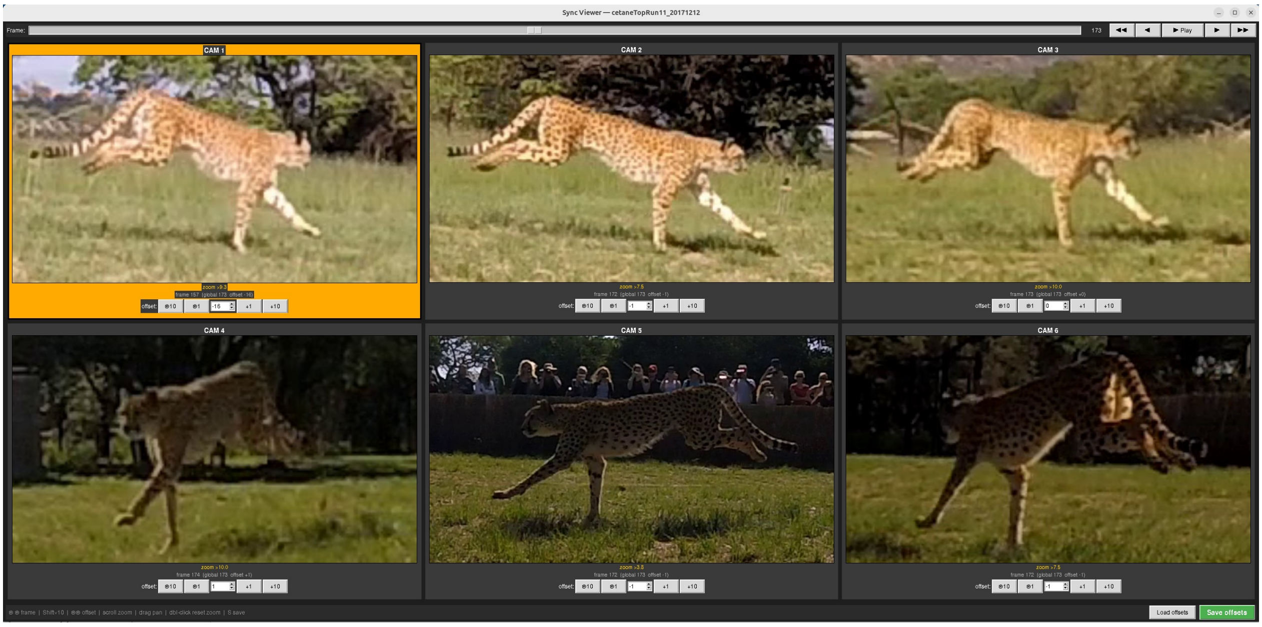


**Figure S4: Custom viewer for event-based camera synchronisation.** The viewer displays all camera views simultaneously and allows rapid frame navigation using the top scroll bar and playback controls, including single-frame and ten-frame forward/backward stepping. Each view can be zoomed independently, and camera-specific frame offsets can be adjusted below each video. Offsets were manually tuned until salient cheetah gait and posture events were temporally aligned across all views. These manual offsets were the starting values for self-calibration (Supplementary Note S1). For the sequence `cetaneTopRun11_20171212`, the manual offsets were cam1 = −16, cam2 = −1, cam3 = 0, cam4 = +1, cam5 = −1, and cam6 = −1; self-calibration changed only cam5, to 0 (Table S3).

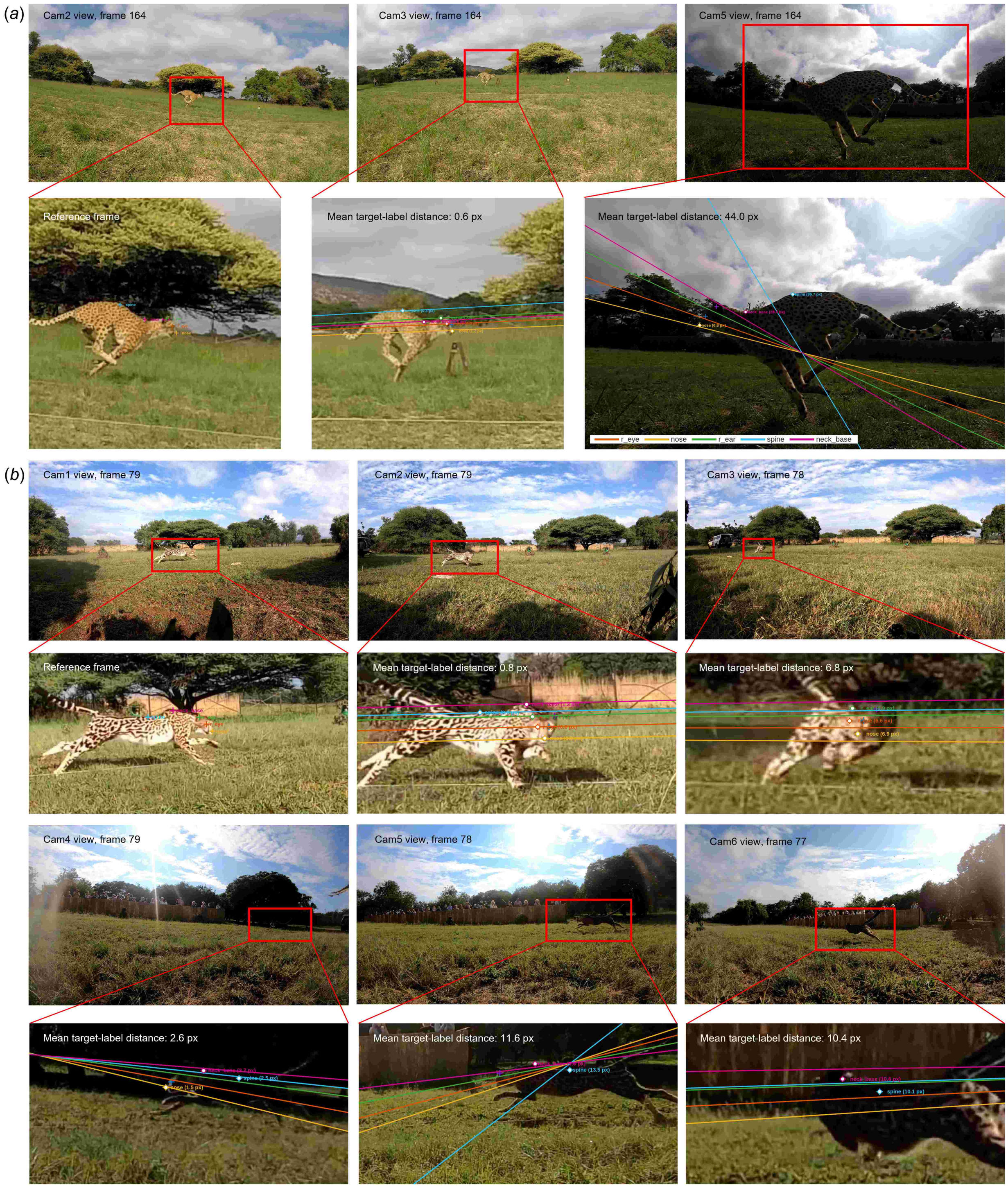


**Figure S5: Epipolar geometry and proximity-amplified calibration bias.** Images are undistorted camera views from representative AcinoSet sequences, shown with the original calibration and manual frame offsets (before self-calibration). Coloured lines show the epipolar lines predicted from manually labelled source points in the reference view; corresponding target-view landmarks are marked with circles, and the annotated values give their perpendicular distance from the predicted line in pixels. (**a**) Close-view configuration (`cetaneTopRun1_20171216`; cameras 2, 3 and 5), in which the cheetah lies near one side of the array and appears much larger in some views. Small residual calibration or sub-frame synchronisation errors therefore produce larger image-space discrepancies: using camera 2 as the reference, the mean target-label distance was 0.6 px in camera 3 but 44.0 px in camera 5. (**b**) More balanced configuration (`julesFlick1_20190309`; cameras 1-6), in which the cheetah has a more similar apparent size across views and the epipolar correspondences generally agree more closely. Using camera 1 as the reference, the mean target-label distance was 0.8 px in camera 2, 6.8 px in camera 3, 2.6 px in camera 4, 11.6 px in camera 5, and 10.4 px in camera6. These examples motivated the self-calibration of camera timing and extrinsics, and the camera–landmark residual weighting, robust loss and consensus checks applied to the remaining inconsistencies, rather than treating close-view residuals as independent labelling noise.

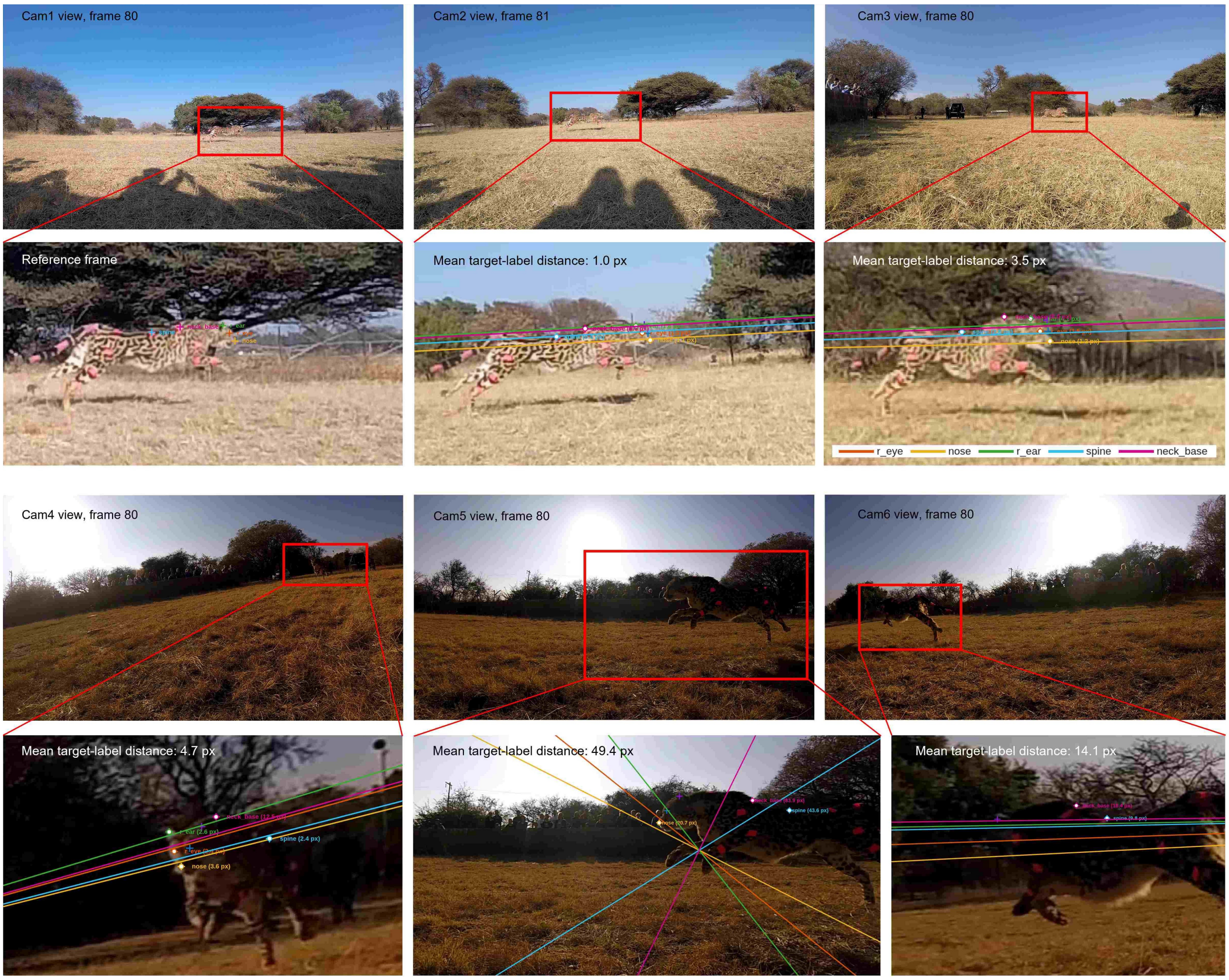


**Figure S6: Epipolar and reprojection diagnostics for `julesTopRun1_20170902`.** Epipolar-geometry check at absolute frame 80. The camera-1 view is used as the reference, coloured lines in cameras 2–6 are the epipolar lines induced by corresponding manually labelled landmarks in the reference view, and the marked target landmarks and annotations show their perpendicular point-to-line distances. These images are displayed in undistorted pixel coordinates and use the original calibration and manual frame offsets (before self-calibration).

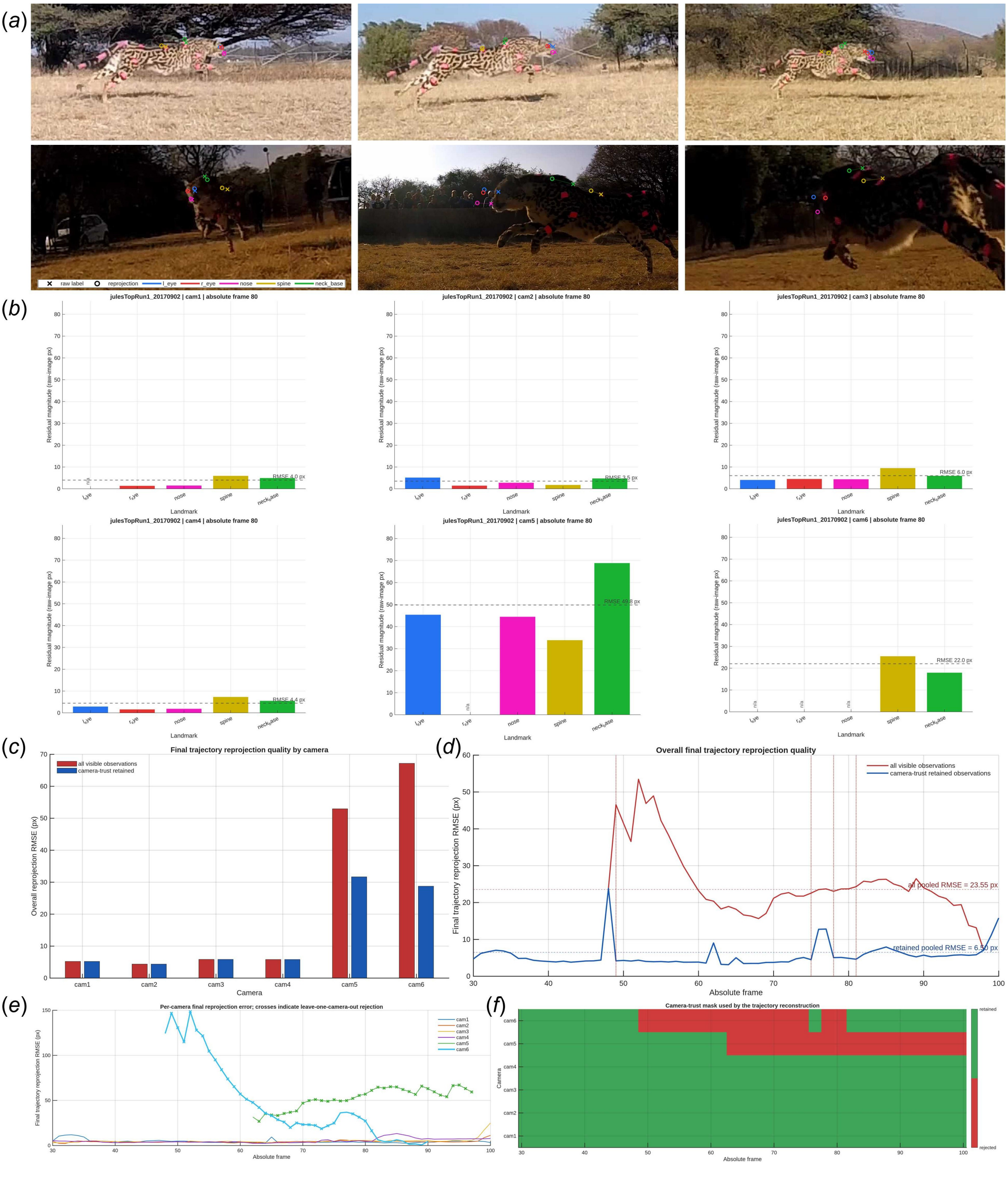


**Figure S7: Reprojection diagnosis of `julesTopRun1_20170902`.** All panels use the original calibration and manual frame offsets (before self-calibration). (**a**) Reprojection overlays at the same absolute frame 80, with the manual per-camera frame offsets applied. Crosses denote raw manual labels, open circles denote projections of the final reconstruction trajectory, connecting segments show the reprojection residuals. (**b**) Bar plots summarise the per-landmark residual magnitudes in raw-image pixels. (**c**) Overall per-camera final reprojection RMSE, comparing all visible observations with camera-trust-retained observations. (**d**) Overall final-trajectory reprojection RMSE over absolute frames; the vertical dotted lines mark the camera-6 rejection intervals. The larger residuals and coherent camera-6 pattern, together with the predominantly horizontal landmark residuals, are consistent with a view-specific, proximity-amplified geometric error. Self-calibration subsequently shifted cameras 4–6 of this sequence by one frame (Table S3). (**e**) Sequence-wide camera diagnostics showing per-camera final-trajectory reprojection RMSE as a function of absolute frame, with crosses indicating LOO camera-trust rejection; (**f**) shows the corresponding retained/rejected camera mask. Camera 6 is rejected over absolute frames 49–75 and 78–81, while camera 5 is rejected during a later portion of the sequence.

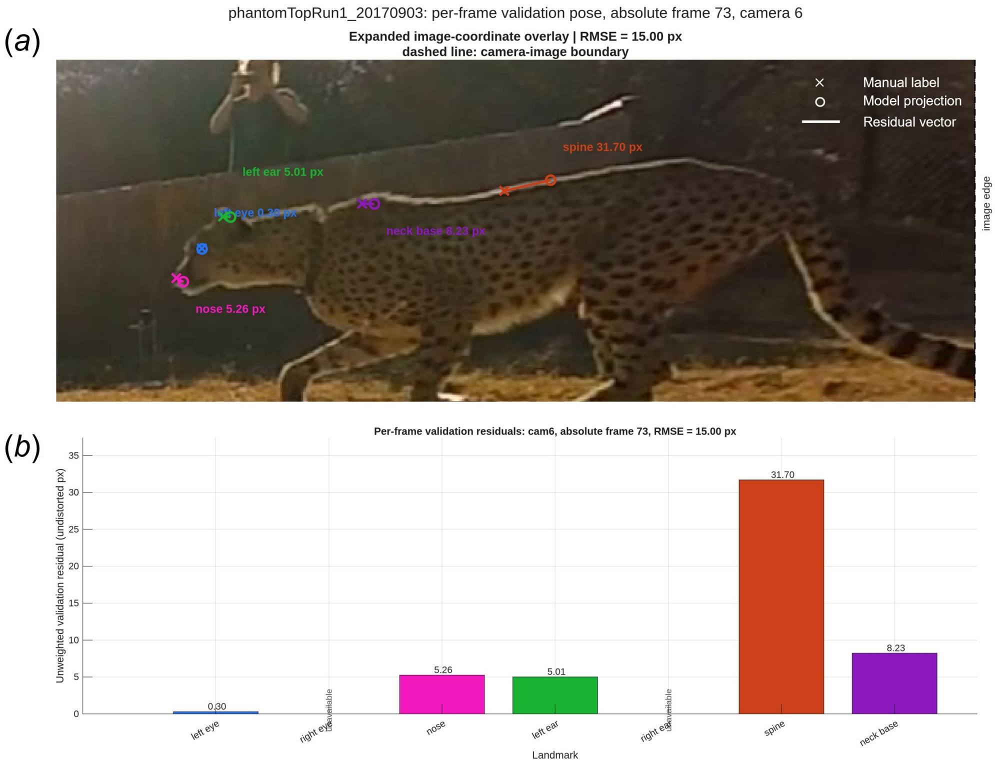


**Figure S8: In-view camera-level reprojection residuals.** Undistorted camera-6 image from `phantomTopRun1_20170903` at frame 73, evaluated using the stored per-frame validation pose. (**a**) shows the expanded image-coordinate overlay: crosses denote observed landmark labels, open circles denote model projections, coloured segments denote residual vectors, and the dashed line marks the camera-image boundary. (**b**) shows the corresponding unweighted residual magnitude for each landmark. The validation RMSE is 10.13 px, computed from five available landmarks; the right-eye and right-ear observations were unavailable.

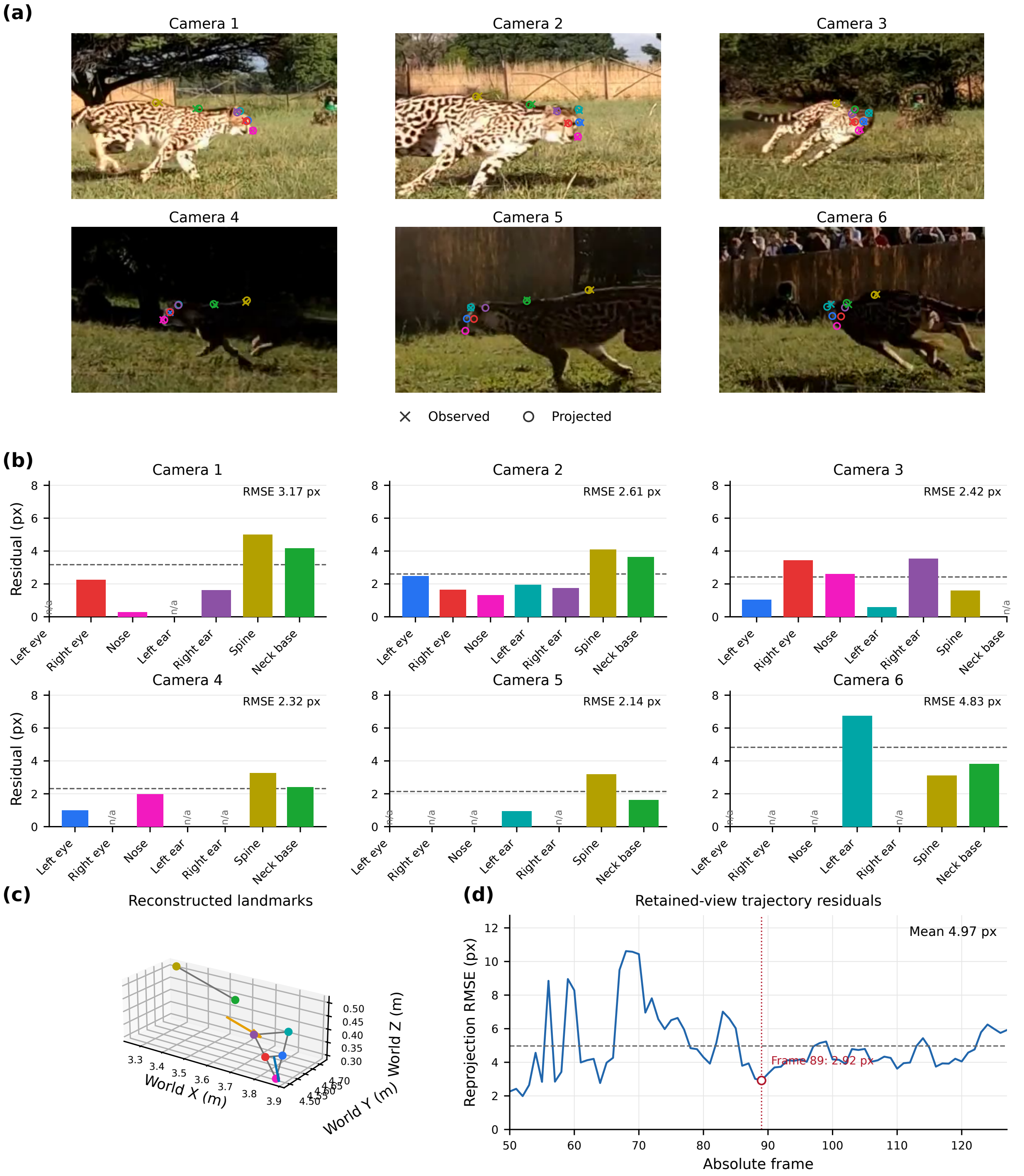


**Figure S9: Multiview reprojection validation of the current trajectory reconstruction at absolute frame 89 of `julesFlick1_20190309`.** (**a**) Undistorted camera views with landmark observations (×) and model projections (open circles), using the corrected camera calibration and timing. Colours identify the seven landmarks in panel (b); segments join corresponding observations and projections. (**b**) Landmark residuals in undistorted image pixels. Dashed lines show camera-specific RMSE; n/a denotes an unavailable observation. (**c**) Reconstructed landmarks in world coordinates, with head (blue) and torso (orange) forward directions. (**d**) Retained-view reprojection RMSE over the 78 analysed frames. The dashed line marks the mean (4.97 px); the red marker identifies frame 89 (2.92 px).

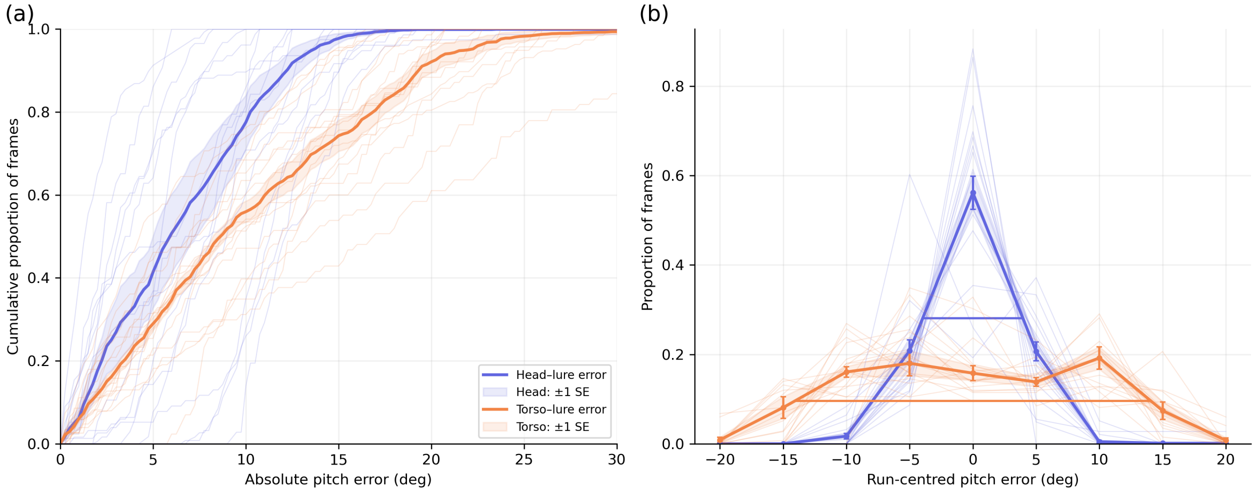


**Figure S10: Animal-equal sensitivity analysis of pitch alignment.** Results comprise 1,590 frames from 16 frontal and lateral pursuits involving 5 cheetahs. Both close pass-by pursuits were excluded. Pursuit-level distributions were averaged within each cheetah before weighting the five animals equally. (**a**) Cumulative distributions of absolute pitch error. Thick curves show animal-equal means, faint curves individual pursuits, and shading ±SE across cheetahs. At 5°, 10° and 15°, head–target proportions were 41.4 ± 8.6%, 77.5 ± 6.5% and 97.7 ± 1.2%, respectively, compared with 29.0 ± 1.5%, 55.9 ± 2.5% and 74.3 ± 3.7% for torso–target error. (**b**) Signed pitch-error distributions after centring each pursuit on its own mean error. Thick curves show animal-equal means and error bars ±SE across cheetahs. Horizontal bars mark FWHM: 7.9° for the head and 28.1° for the torso.

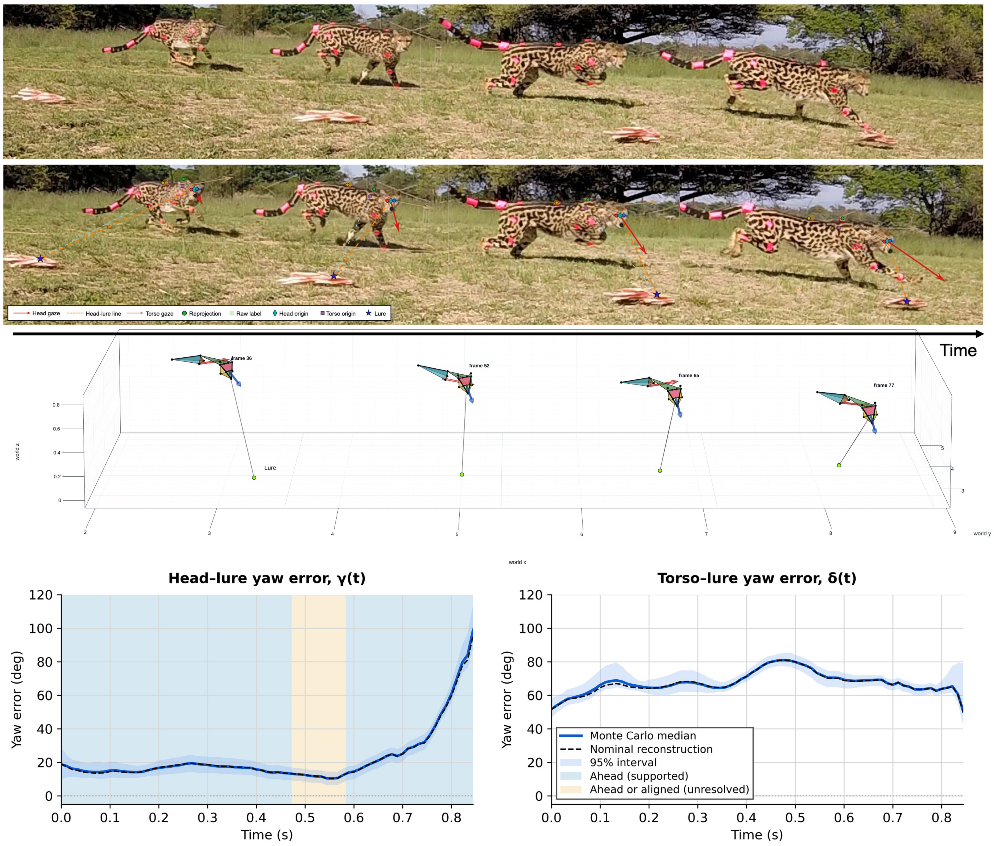


**Figure S11: Representative reconstruction of lateral pursuit.** The example shows `julesTopRun1_20171209`. From top to bottom: image sequences; projected landmarks, head and torso directions, and head–target lines; reconstructed poses and target positions at absolute frames 36, 52, 65 and 77; and head–target yaw error $\gamma(t)$ (left) and torso–target yaw error $\delta(t)$ (right). Time progresses from left to right through the snapshots. The trajectories contain 77 retained frames (absolute frames 10–86). Solid blue curves show medians across 500 joint Monte Carlo reconstructions, dashed black curves show the nominal reconstruction, and envelopes show 95% reconstruction intervals. Background shading in the head-yaw plot marks supported ahead orientation (blue) and frames unresolved between ahead and aligned (pale orange). Ahead requires an absolute yaw error exceeding 10° and a positive travel-directed offset, with at least 95% support across the joint draws.

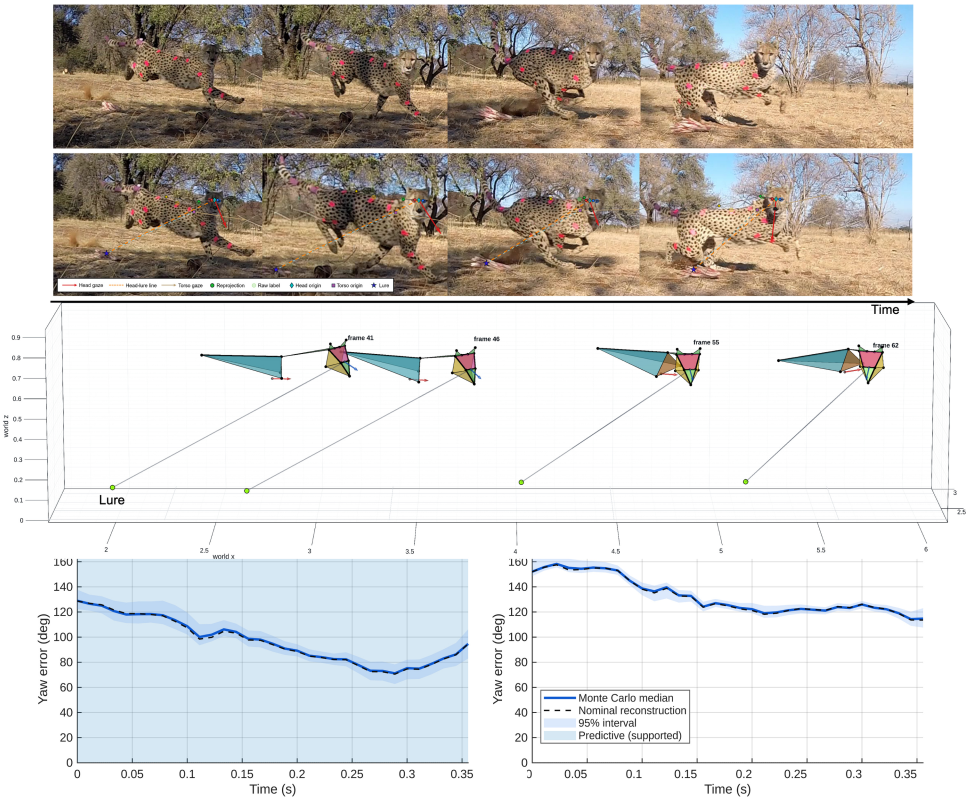


**Figure S12: Representative reconstruction of close pass-by pursuit.** The example shows `zorroBottomFlick2_20170903`. From top to bottom: image sequences; projected landmarks, head and torso directions, and head–target lines; reconstructed poses and target positions at absolute frames 41, 46, 55 and 62; and head–target yaw error $\gamma(t)$ (left) and torso–target yaw error $\delta(t)$ (right). Time progresses from left to right through the snapshots. Solid blue curves show Monte Carlo medians, dashed black curves show the nominal reconstruction, and envelopes show 95% reconstruction intervals. The blue background in the head-yaw plot denotes supported ahead-of-target orientation. All 83 retained frames meet the joint ahead criterion: an absolute yaw error exceeding 10° and a positive travel-directed offset, with at least 95% support across 500 joint reconstruction draws.

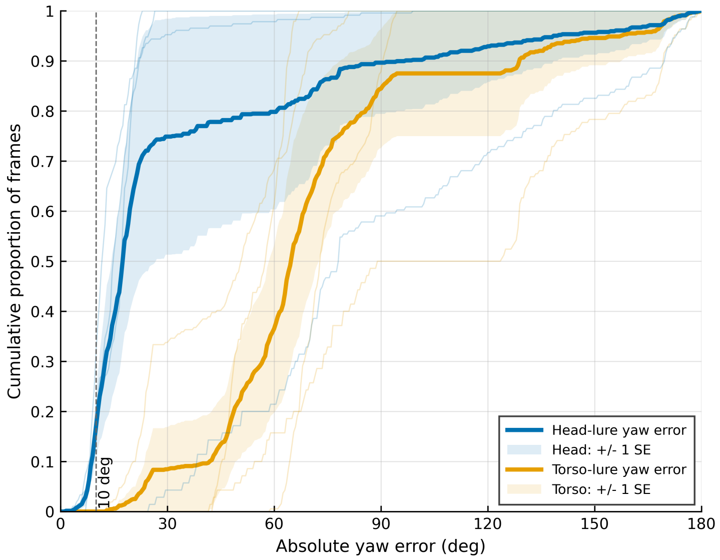


**Figure S13: Animal-equal sensitivity analysis of absolute head–target and torso–target yaw error.** The analysis comprises 628 matched frames from 4 cheetahs: 5 lateral and 2 close pass-by pursuits. Errors are absolute values of wrapped Monte Carlo-median yaw errors. Run ECDFs were averaged within each cheetah before weighting the four animals equally. Thick blue and orange solid curves show the head and torso means, respectively; coloured envelopes denote ±1 SE across animals, and thin curves show individual-animal ECDFs. The dashed vertical line marks 10°, within which the animal-equal fractions were 17.1% for the head and 0% for the torso. SE bands describe variation across animals.

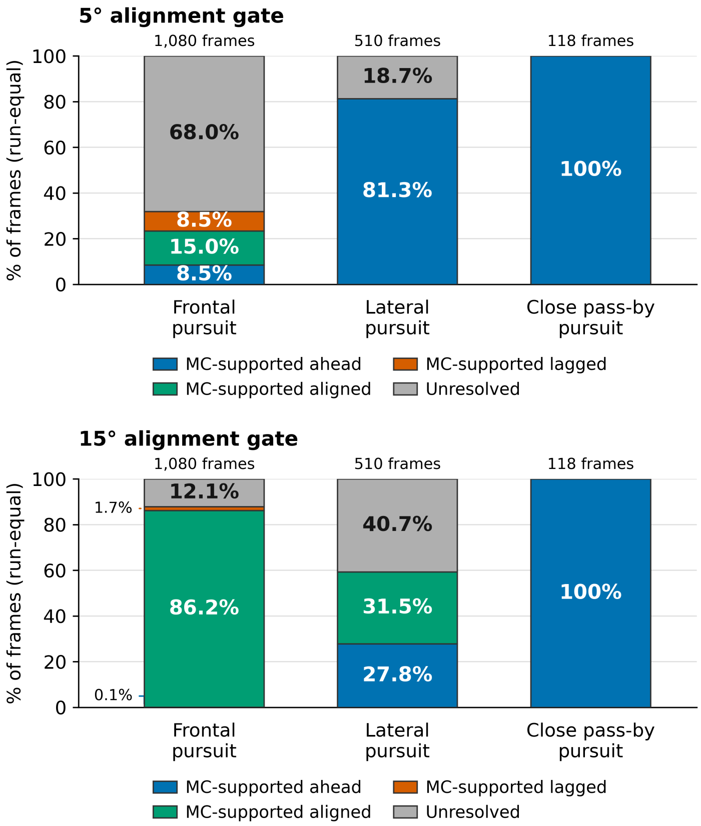


**Figure S14: Sensitivity of Monte Carlo-supported head-yaw states to the alignment gate across all 18 pursuits.** The upper and lower panels use alignment gates of 5° and 15°, respectively, for 11 frontal pursuits (1,080 frames), 5 lateral pursuits (510 frames) and 2 close pass-by pursuits (118 frames), involving 6 cheetahs. Bars average within-run frame fractions equally within each context; counts above bars give analysed frames. In each joint reconstruction draw, aligned means $|\gamma| \leq \theta_{\text{align}}$. Outside this band, ahead and lagged states have positive and negative travel-directed head–target yaw offsets, respectively, using the same local target-travel criterion as main-text figure 4f. A frame requires at least 95% support across the same 500 joint draws for assignment to a state. All remaining frames are unresolved and remain in the denominator. Blue, green, orange and grey denote ahead, aligned, lagged and unresolved states, respectively. The primary 10° gate is shown in main-text figure 4f.

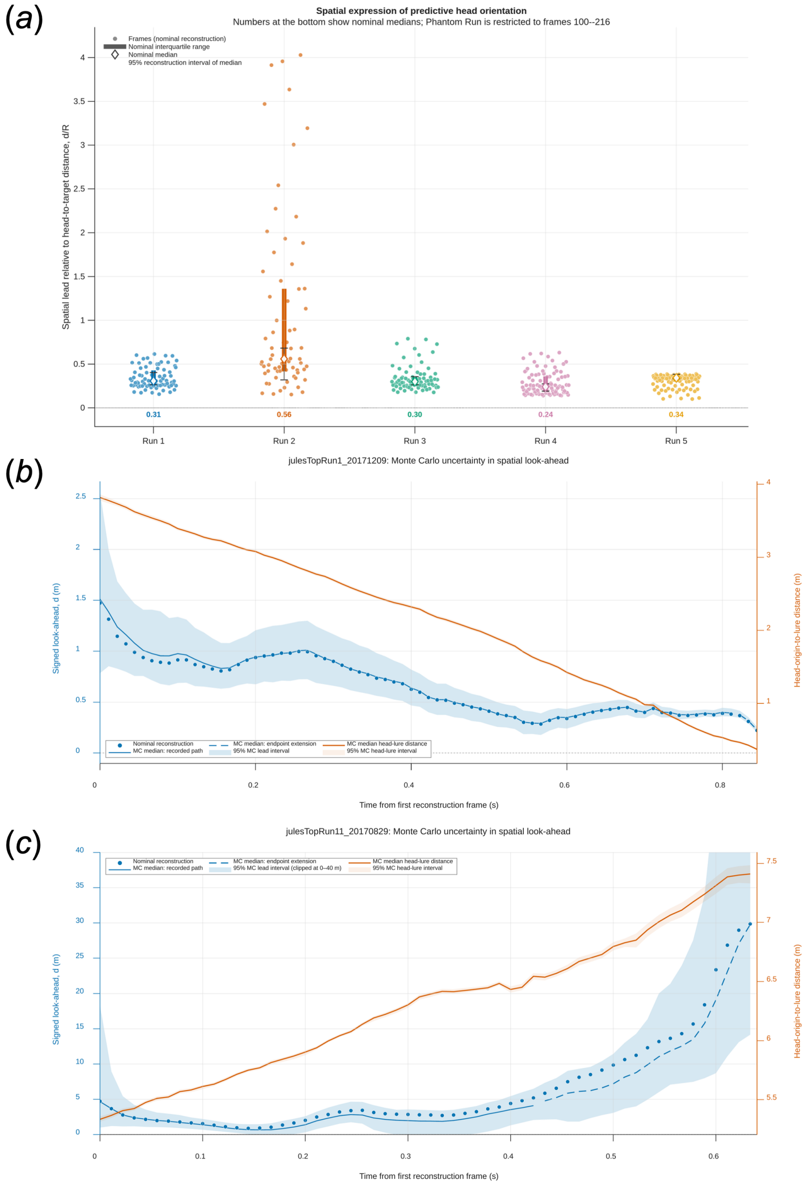


**Figure S15: Spatial expression of ahead-of-target head orientation.** (**a**) Signed distance ahead along the reconstructed target path divided by current 3D head–target distance, $d/R$, across five lateral pursuits. Points are nominal frames with a forward ray–path intersection; bars show nominal IQRs, diamonds nominal medians, and capped lines 95% reconstruction intervals for the median across 500 joint draws. Runs 1–5 are `julesFlick1_20190309`, `julesTopRun11_20170829`, `julesTopRun1_20171209`, `lilyRun_20190309` and `phantomRun_20190303`, contributing 78, 70, 77, 96 and 113 nominal intersections. Phantom uses available frames 100–212 within the predefined 100–216 window. Medians range from 0.24 to 0.56. All panels include forward endpoint continuation where the head ray misses the recorded path. (**b,c**) Spatial lead and head–target range for `julesTopRun1_20171209` and `julesTopRun11_20170829`. These trajectories include forward extension of the recorded path along its terminal horizontal direction at constant height. Lines show Monte Carlo medians and envelopes 95% reconstruction intervals. Dashed segments identify frames where the nominal estimate or at least half the draws use the endpoint extension; crosses mark frames with less than 95% intersection support. Direction uncertainty is propagated within each draw, conditional on continued straight travel. Panel (c) displays times before 0.64 s and a 0–40 m spatial-lead range; the uncertainty envelope extends beyond the displayed limits. These display limits do not affect the calculations.

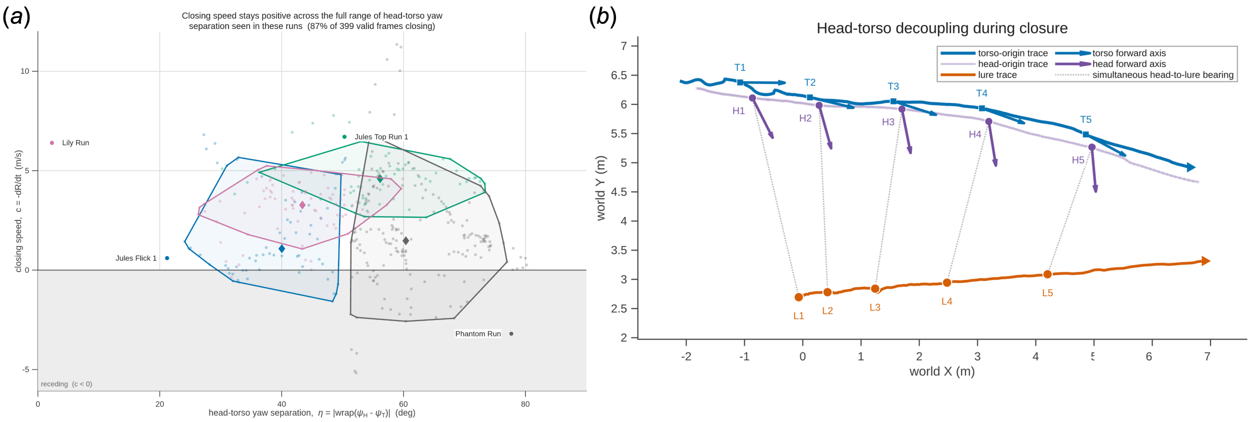


**Figure S16: Head–torso separation during continued approach to the target.** (**a**) Absolute head–torso yaw separation versus closing speed in four lateral pursuits with net range closure. Points represent 399 valid frames, diamonds run medians, and outlines the central 90% of each run; 87.2% of frames have positive closing speed. (**b**) Top view of `phantomRun_20190303`, showing head, torso and lure trajectories with simultaneous orientation axes and head-to-lure bearings at five snapshots.

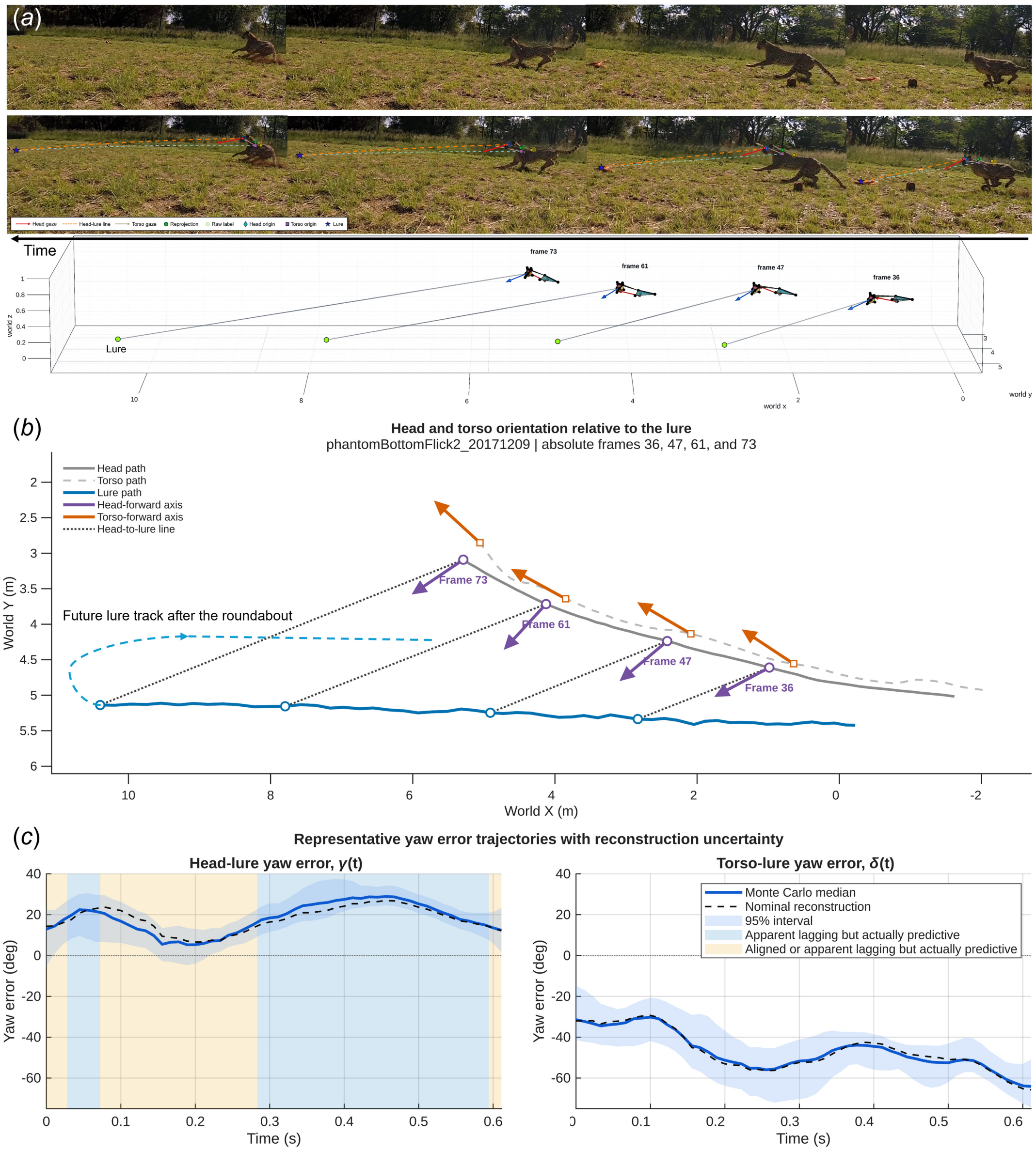


**Figure S17: Qualitative example of head orientation during turning pursuit.** (**a**) Image sequence, projected landmarks and directions, and reconstructed poses for `phantomBottomFlick2_20171209` at absolute frames 36, 47, 61 and 73. The arrow indicates temporal progression from right to left. (**b**) Top view of head, torso and lure trajectories and simultaneous orientations. The dashed extension illustrates the lure's subsequent path around the turn, allowing comparison of head orientation with both current target bearing and the post-turn path. (**c**) Head–lure and torso–lure yaw-error trajectories. Solid blue lines show Monte Carlo medians, dashed black lines the nominal reconstruction and envelopes 95% reconstruction intervals. Background shading distinguishes the illustrated alignment and apparent-lagging categories. The predictive interpretation stated in the panel legend is a hypothesis about orientation towards the subsequent path, not an independently established tracking state. This example provides qualitative context and is excluded from the 18-pursuit quantitative cohort and the five-lateral-pursuit model comparison.

### Supplementary Notes

#### Supplementary Note S1. Camera synchronisation and calibration

The AcinoSet GoPro cameras were not hardware-synchronised and shared a single generic intrinsic calibration. Their extrinsics were estimated from checkerboard recordings with few common frames per camera pair. We used landmark annotations to diagnose residual timing and geometric errors and to refine camera timing and extrinsics in the first stage of CRBR.

##### Initial event-based camera frame offsets

We first aligned camera views manually using a custom multiview viewer (figure S4). Foot touchdown, limb orientation and head–body posture provided corresponding events across views. Several events per sequence were checked to distinguish poses that remained similar across consecutive frames. These integer offsets provided the initial values and prior centres for self-calibration; final offsets are listed in table S3.

##### Camera models

For each camera $i$, the AcinoSet scene-calibration JSON file supplies an intrinsic matrix $K_i$, four distortion coefficients $D_i$, and a world-to-camera rotation $R_{cw,i}$ and translation $\mathbf{t}_{cw,i}$. The coordinate convention and corresponding camera-to-world pose are

$$\mathbf{X}_{c,i} = R_{cw,i}\mathbf{X}_w + \mathbf{t}_{cw,i}, \qquad R_{wc,i} = R_{cw,i}^{\top}, \qquad \mathbf{C}_{w,i} = -R_{cw,i}^{\top}\mathbf{t}_{cw,i}. \tag{S1}$$

Undistortion used MATLAB's four-coefficient `cameraIntrinsicsKB` model, constructed from $K_i$ and $D_i$ by the helper `cameraIntrinsicsFromOpenCV`. The JSON files did not explicitly specify their distortion convention.

For landmark $p$ reconstructed from pose $\boldsymbol{\theta}_f$ at frame $f$, the predicted undistorted image position and reprojection residual are

$$\widehat{\mathbf{y}}_{i,p,f} = \pi_i\left(R_{cw,i}\mathbf{X}_p(\boldsymbol{\theta}_f) + \mathbf{t}_{cw,i}\right), \qquad \mathbf{r}_{i,p,f} = \mathbf{y}_{i,p,f} - \widehat{\mathbf{y}}_{i,p,f}, \tag{S2}$$

where $\pi_i$ projects into undistorted image coordinates. Intrinsics were used as supplied. Refined extrinsics and frame offsets were held fixed during subsequent pose fitting and smoothing.

##### Epipolar-geometry check

We assessed agreement between views using epipolar distances. For cameras $i$ and $j$, the relative pose and fundamental matrix were

$$\begin{aligned} R_{ji} &= R_{cw,j}R_{cw,i}^{\top}, \qquad \mathbf{t}_{ji} = \mathbf{t}_{cw,j} - R_{ji}\mathbf{t}_{cw,i}, \\ F_{ji} &= K_j^{-\top}[\mathbf{t}_{ji}]_{\times}R_{ji}K_i^{-1}. \end{aligned} \tag{S3}$$

A homogeneous source point $\widetilde{\mathbf{x}}_i$ defines the target-view line $F_{ji}\widetilde{\mathbf{x}}_i = (a, b, c)^{\top}$. Its perpendicular distance from the corresponding labelled point $(u_j, v_j)$ is

$$d_{\text{epi}} = \frac{|au_j + bv_j + c|}{\sqrt{a^2 + b^2}}. \tag{S4}$$

Figure S5 shows these distances before self-calibration, using the original camera geometry and manual frame offsets. Larger discrepancies in close views are consistent with increased sensitivity to geometric errors, but epipolar distances alone cannot separate calibration, timing and landmark-localisation errors.

#### Image-space sensitivity to calibration and timing errors

To interpret the larger discrepancies in close views, consider a point at camera depth $Z$ in the undistorted perspective image. Small lateral camera-translation errors produce approximately

$$|\delta u| \approx \frac{f_x|\delta t_x|}{Z}, \qquad |\delta v| \approx \frac{f_y|\delta t_y|}{Z}, \tag{S5}$$

where $f_x$ and $f_y$ are focal lengths in pixels. The same metric error therefore produces a larger pixel displacement at shorter range. This dependence is specific to translation-like errors: rotation errors give approximately depth-independent angular displacements at a fixed viewing direction, while intrinsic-parameter errors follow different relationships.

A timing offset $\Delta t$ produces a displacement approximately equal to image velocity multiplied by $\Delta t$. We examined this component by triangulating landmarks without one camera and projecting them into the held-out view. In the most inconsistent views, residuals followed the animal's image velocity. For example, camera 5 of `cetaneTopRun1_20171216` had median residual components of 77 px along and 7 px across image motion. Shifting labels by one to three frames largely removed the discrepancies in these views. Refining extrinsics after timing correction further reduced held-out error, indicating an additional calibration component.

#### Spatiotemporal self-calibration from manual annotations

Self-calibration used the manual annotations to estimate camera timing and extrinsics jointly for each sequence, with free landmark trajectories and no rigid-body template. The corrected scene calibration, frame offsets and labels were passed to the subsequent stages of CRBR. Algorithm S1 gives the fitting, held-out selection and export procedure.

**Model and priors.** The annotation in video row $r$ of camera $c$ was modelled as captured at absolute frame time

$$t_{c,r} = r - o_c - d_c, \tag{S6}$$

where $o_c$ is the manual integer frame offset and $d_c$ a continuous timing correction. Each landmark $j$ was represented by a free 3D trajectory $\mathbf{X}_j(t)$ defined at integer frames and evaluated at fractional times by Catmull–Rom interpolation. Labels were predicted with the full fisheye camera model in raw image pixels, $\hat{\mathbf{u}}_{c,j,r} = \pi_c(\mathbf{X}_j(t_{c,r}))$, with camera rotation $R_c = \exp([\boldsymbol{\omega}_c]_\times)R_c^0$ and centre $\mathbf{C}_c = \mathbf{C}_c^0 + \Delta\mathbf{C}_c$ perturbed from the supplied values $R_c^0$ and $\mathbf{C}_c^0$. Trajectories, timing corrections and extrinsic corrections minimised

$$E = \sum_{c,j,r} \rho_{2.5}\left(\frac{\|\hat{\mathbf{u}}_{c,j,r} - \mathbf{u}_{c,j,r}\|}{\sigma_j}\right) + \sum_{j,t} \rho_3\left(\frac{\|\Delta^2\mathbf{X}_j(t)\|}{\sigma_a}\right) + \frac{1}{2}\sum_c \left(\frac{d_c^2}{\sigma_d^2} + \frac{\|\boldsymbol{\omega}_c\|^2}{\sigma_\omega^2} + \frac{\|\Delta\mathbf{C}_c\|^2}{\sigma_C^2}\right), \tag{S7}$$

where $\rho_k$ is the Huber loss with threshold $k$, $\sigma_j$ is the annotation scale of landmark $j$ used in pose fitting, and $\Delta^2$ is the second temporal difference, with $\sigma_a$ corresponding to an acceleration of 100 m s$^{-2}$. The priors $\sigma_d = 2$ frames, $\sigma_\omega = 0.5°$ and $\sigma_C = 5$ cm are centred on the manual offsets and the supplied extrinsics, so that a camera with little information retains its original values. A similarity transformation applied to all cameras together cannot be determined from the images, and this seven-degree-of-freedom component of the corrections was constrained to zero. Equation (S7) was minimised by sparse, iteratively reweighted least squares in two stages: timing only, started from the manual offsets and from a pairwise epipolar estimate (the lower-cost solution was retained), and then timing and extrinsics jointly. The lure trajectory was subsequently fitted with the camera geometry fixed.

**Held-out model selection.** Three camera geometries were compared for every sequence: the original geometry, timing correction only, and timing plus extrinsic correction. Each sequence was divided into four contiguous time blocks. For each block, the geometry was estimated from the other three blocks; the landmark trajectories in the held-out block were then reconstructed without one camera and projected into that camera, so that neither the held-out frames nor the predicted camera's labels contributed to the prediction. Performance was summarised as the median held-out label error in raw pixels. Extrinsic refinement was adopted only if it did not increase this error relative to timing correction alone. A camera whose refined extrinsics predicted its own held-out labels more than 25% worse than timing correction alone retained its original extrinsics, provided the overall held-out error did not increase.

**Export.** The refined extrinsics were written to the scene calibration file and the integer part of each corrected offset, $o_c + \mathrm{round}(d_c)$, to the frame-offset table. The sub-frame remainder $\phi_c = d_c - \mathrm{round}(d_c)$, with $|\phi_c| \leq 0.5$ frame, was removed from the labels themselves. Each label assigned to frame $f$ but captured at $f - \phi_c$ was moved by the image displacement predicted by the fitted motion,

$$\mathbf{u}'_{c,j,f} = \mathbf{u}_{c,j,f} + \pi_c\big(\mathbf{X}_j(f)\big) - \pi_c\big(\mathbf{X}_j(f - \phi_c)\big)\,. \tag{S8}$$

Only the predicted motion is removed; the residual of each manual annotation is preserved.

**Corrections.** Table S3 lists the corrections for all 20 reconstructed sequences. Timing plus extrinsic correction was selected for 18 of them and timing correction alone for the remaining two. Integer offsets changed for 37 of 113 cameras, the median absolute sub-frame correction was 0.15 frame (maximum 0.46), and extrinsic updates did not exceed 2.2° in rotation or 13.5 cm in camera position. The median held-out label error decreased in every sequence, from 4.6–28.6 px to 2.6–6.7 px.

**Validation.** Two tests assessed whether self-calibration recovers the correct geometry. First, synthetic sequences were generated from the fitted trajectories, cameras and visibility pattern of four self-calibrated sequences (six simulations), with known timing errors of up to ±2.5 frames, camera perturbations of approximately 0.7° and 6 cm, and annotation noise at the pipeline scales. Starting from the perturbed geometry and the original offsets, self-calibration reduced the median head-axis error from 21–72° to 6.2–7.8°, the torso-axis error from 7.1–20.4° to 3.5–7.7°, and the 3D landmark error from 90–175 mm to 23–37 mm. These values approach a reference reconstruction with the true geometry (head 6.7–7.7°, torso 3.4–5.6°, landmarks 18–25 mm), which reflects annotation noise alone. Second, two copies of `zorroBottomFlick2_20170829` with different manual offsets reconstructed with reprojection errors of 20–38 px and 4–7 px, respectively. Self-calibration of the poorly synchronised copy recovered offsets of 0, 1.7, −0.4, 0.3, 1.8 and 0.2 frames relative to camera 1, in agreement with the hand-corrected integer offsets except for camera 6. Its reconstruction agreed with the self-calibrated hand-corrected copy to within 1.2° in head pitch, 0.1° in torso pitch and 3.2° in torso yaw (medians), compared with differences of up to 35–56° (90th percentile) without self-calibration. A remaining constant head-yaw difference of about 7° was attributable to the absence of ear annotations in the poorly synchronised copy.

**Limitations.** No ground-truth 3D pose is available for the recorded sequences, so the evidence rests on held-out prediction, synthetic recovery and the natural experiment above. Timing and extrinsic errors are partly confounded during near-constant-velocity running, because a time offset and a camera translation along the direction of travel produce almost identical image errors. Individual offsets and camera positions are therefore not uniquely identified (in the synthetic tests, offsets were recovered up to a common shift of 0.2–1.4 frames, absorbed by the reference camera's position) and should not be interpreted as exact clock offsets, although the reconstruction is unaffected. Each camera was assumed to have a constant offset. Offsets re-estimated in sliding windows were stable for most cameras (window-to-window interquartile range 0.01–0.07 frame), but camera 1 of `julesBottomFlick21_20171221` was not: its windowed offset varied between −0.8 and +1.3 frames, with the largest deviations during the turn around the roundabout, which a single constant correction cannot fully represent. The shared intrinsic calibration was not refined: per-camera focal-length and rolling-shutter terms were tested by cross-validation and gave no consistent improvement. Finally, the closest views retained larger residuals than the other cameras (for example, cameras 5 and 6 of `cetaneTopRun1_20171216`); these were handled by the robust procedures below.

Remaining inconsistencies were handled through camera-trust filtering (Supplementary Note S2) and observation-level weighting (Supplementary Note S3).

**Algorithm S1** Spatiotemporal camera self-calibration and label correction.

**Require:** Raw landmark and lure annotations $\mathbf{u}_{c,j,r}$, camera intrinsics and initial extrinsics, manual offsets $o_c$, frame rate
**Ensure:** Selected camera geometry, integer offsets, motion-corrected labels
Define candidates $O$ (original), $T$ (timing only), and $J$ (joint timing and extrinsics)
In each fit, estimate free landmark trajectories using equation (S7), without rigid-body templates
For $T$ and $J$, initialise timing from manual offsets and an epipolar estimate; retain the lower-cost timing-only fit
For $J$, refine timing and extrinsics jointly, constraining the common seven-degree-of-freedom similarity correction to zero
*Held-out evaluation and selection*
**for** each candidate $m \in \{O, T, J\}$ **do**
  **for** each of four contiguous temporal blocks $B$ **do**
    Fit candidate $m$ to observations outside $B$
    **for** each camera $c$ **do**
      With geometry fixed, reconstruct trajectories in $B$ from the other cameras and predict camera $c$'s labels
      Store held-out raw-pixel reprojection errors
    **end for**
  **end for**
  Compute pooled median $E_m$, mean $A_m$, and per-camera medians $E_{m,c}$ and counts $n_{m,c}$
**end for**
Select $m^* \leftarrow T$
**if** $E_J \le E_T$ and $A_J \le 1.10 A_T$ **then**
  $m^* \leftarrow J$
  $F \leftarrow \{c : n_{T,c} \ge 30,\ E_{J,c} > 1.25 E_{T,c}\}$, using finite per-camera medians
  **if** $F \ne \varnothing$ **then**
    Repeat joint fitting and cross-validation with extrinsics of cameras in $F$ fixed at their original values; call this candidate $J_F$
    **if** $E_{J_F} \le E_J$ and $A_{J_F} \le A_J$ **then**
      $m^* \leftarrow J_F$
    **end if**
  **end if**
**end if**
**if** $E_O < E_{m^*}$ and $A_O < A_{m^*}$ **then**
  $m^* \leftarrow O$
**end if**
*Final fit and export*
Refit the selected model to all available observations; for $O$, fit trajectories with camera geometry and timing fixed
Fit the lure trajectory with selected camera geometry fixed; refine timing from lure labels only for cameras without body observations
**for** each camera $c$ **do**
  $o'_c \leftarrow o_c + \mathrm{round}(d_c)$; $\phi_c \leftarrow d_c - \mathrm{round}(d_c)$
  **for** each valid label at integer-offset-corrected frame $f$ **do**
    $\mathbf{u}'_{c,j,f} \leftarrow \mathbf{u}_{c,j,f} + \pi_c(\mathbf{X}_j(f)) - \pi_c(\mathbf{X}_j(f - \phi_c))$
  **end for**
**end for**
Export corrected camera geometry, $o'_c$, labels $\mathbf{u}'$ and the calibration report for downstream pose reconstruction

### Supplementary Note S2. Multiview triangulation and camera trust

We triangulated landmarks to initialise pose and estimate animal-specific scale. Camera-trust filtering identified persistently inconsistent views before the final triangulation and pose fit.

#### Aligned observations and undistortion

We collected simultaneous observations using the corrected integer offsets and sub-frame-aligned labels from Supplementary Note S1.

For a raw labelled pixel location $\mathbf{y}^{\text{raw}}_{c,j,t} = (u^{\text{raw}}, v^{\text{raw}}, 1)^{\mathsf{T}}$ of landmark $j$ at frame $t$, we calculated its undistorted pixel coordinate as

$$\mathbf{y}_{c,j,t} = \Pi\left[\mathbf{K}_c\, \mathcal{D}_c^{-1}\left(\mathbf{K}_c^{-1}\mathbf{y}^{\text{raw}}_{c,j,t}; \mathbf{D}_c\right)\right], \tag{S9}$$

where $\mathcal{D}_c^{-1}$ denotes the inverse lens-distortion mapping defined by the calibrated distortion coefficients $\mathbf{D}_c$, and $\Pi$ converts homogeneous image coordinates to 2D pixels. All triangulation and reprojection calculations used the resulting undistorted coordinates $\mathbf{y}_{c,j,t}$.

#### Multiview triangulation

For each landmark $j$ at frame $t$, all valid visible observations were assembled into a multiview point track. A 3D point was estimated only when at least two camera views were available. We used MATLAB's `pointTrack` and `triangulateMultiview` functions to estimate the world-coordinate landmark position.

The initial all-camera triangulation was used to calculate diagnostic reprojection errors and to support the LOO trust assessment described below. After camera trust had been assessed, the anatomical landmarks were triangulated again using retained cameras only. These final triangulated points were used for scale estimation and pose initialisation.

#### LOO trust assessment

A camera can appear to fit well in an ordinary joint triangulation even when it contains a coherent calibration, annotation or synchronisation bias, because the triangulated point is pulled towards that camera. We therefore assessed each camera using held-out reprojection error. As derived in S1, translation-like camera-level calibration errors can be amplified in close-view configurations. Hence, making an independent assessment of each camera is particularly important.

For every visible landmark in a frame, each camera was held out in turn. The landmark was triangulated from the remaining views and reprojected into the held-out camera. If camera $c$ was held out, its landmark-specific LOO residual was

$$e^{\text{LOO}}_{c,j,t} = \left\|\pi_c\left(\hat{\mathbf{X}}^{(-c)}_{j,t}\right) - \mathbf{u}^{\text{obs}}_{c,j,t}\right\|_2, \tag{S10}$$

where $\hat{\mathbf{X}}^{(-c)}_{j,t}$ is the 3D point triangulated without camera $c$. An LOO residual was calculated only when at least three views of the landmark were available, such that at least two cameras remained after withholding one view.

For each camera and frame, the camera-level score was the median LOO residual across all landmarks for which a held-out estimate could be obtained:

$$E^{\text{LOO}}_{c,t} = \text{median}_j\left(e^{\text{LOO}}_{c,j,t}\right). \tag{S11}$$

A camera was provisionally classified as an outlier when its score exceeded both three times the median score of the other scored cameras and 8 pixels:

$$E^{\text{LOO}}_{c,t} > \max\left[3\,\text{median}_{c'\neq c}\left(E^{\text{LOO}}_{c',t}\right),\ 8\text{ pixels}\right]. \tag{S12}$$

**Algorithm S2** LOO camera-trust assessment.

**Require:** Undistorted landmark observations, calibrated camera models
**Ensure:** Frame-specific retained-camera mask
Initialise every camera as retained
**for** each frame $t$ **do**
    **for** each camera $c$ **do**
        Initialise an empty set of held-out residuals
    **end for**
    **for** each anatomical landmark $j$ **do**
        Collect all visible camera observations of landmark $j$
        **if** at least three views are available **then**
            **for** each viewing camera $c$ **do**
                Hold out camera $c$
                Triangulate landmark $j$ from the remaining views
                Reproject the point into camera $c$
                Store the held-out reprojection residual $e_{c,j,t}^{\mathrm{LOO}}$
            **end for**
        **end if**
    **end for**
    **for** each camera $c$ **do**
        $E_{c,t}^{\mathrm{LOO}} \leftarrow$ median of its available held-out residuals
    **end for**
    **if** at least four cameras are present and at least three have scores **then**
        **for** each scored camera $c$ **do**
            Compare $E_{c,t}^{\mathrm{LOO}}$ with the median score of the other cameras
            Flag $c$ if it exceeds both the relative threefold threshold and the 8-pixel absolute threshold
        **end for**
    **end if**
    Update each camera's retained/rejected state after two consecutive outlier or non-outlier frames
    Re-admit cameras if necessary to retain at least three visible views
**end for**
**for** each frame $t$ and landmark $j$ **do**
    Re-triangulate landmark $j$ from retained cameras only
    Store the point if at least two retained observations are available
**end for**

Camera rejection was considered only when at least four cameras had visible observations in that frame and at least three cameras had finite LOO scores. To avoid frame-to-frame flicker, an outlier classification had to persist for two consecutive frames before a camera was excluded; similarly, two consecutive non-outlier frames were required before a rejected camera was re-admitted. At least three visible cameras were always retained. The resulting frame-specific camera mask was used both for final triangulation and for the subsequent nonlinear pose fitting. Figure S6 shows a large camera-5 epipolar discrepancy, consistent with a view-specific geometric error. Figure S7 shows how the LOO mask identifies such inconsistent views and yields lower reprojection error for the retained observations than for all observations.

#### Lure reconstruction

The lure target was reconstructed independently of the head–torso rigid-body model. Its labelled 2D positions were corrected for lens distortion and triangulated directly from available camera views, requiring at least two valid observations. No anatomical template, kinematic constraint or temporal smoothing was imposed on the lure trajectory.

### Supplementary Note S3. Rigid-body templates, scale estimation and framewise pose fitting

To estimate stable anatomical axes for pitch and yaw analysis, we fitted separate rigid head and torso models directly to the retained multiview observations. The templates constrained landmark geometry when visibility was incomplete or individual annotations were inconsistent. This was particularly relevant to neck-base and spine landmarks, whose apparent positions could vary with viewpoint, occlusion, fur and body deformation. Triangulated landmarks supplied animal-specific scales and initial poses; the lure was triangulated independently.

#### Rigid-body template

The head template used the eyes, nose and ears, with its origin at the inter-eye midpoint. The ears extended the landmark configuration backwards and laterally, helping constrain head pose when facial landmarks were occluded or viewed obliquely; they were not used as gaze indicators. The torso template used the neck base and mid-spine to estimate the longitudinal body axis. Shoulder landmarks were excluded because they were difficult to label consistently and moved relative to the trunk during running. Two midline landmarks constrain torso heading and inclination but do not identify rotation about the torso axis.

Template coordinates were defined in each segment's local frame and multiplied by the estimated head or torso scale to account for animal size.

For a head landmark $j$, its model-predicted world position was

$$\mathbf{X}_{j,t}^{\text{model}} = \mathbf{o}_{H,t} + R_{H,t}\left[s_H \mathbf{q}_{H,j}\right], \tag{S13}$$

where $\mathbf{q}_{H,j}$ is its local template coordinate and $s_H$ is the head scale. The corresponding torso landmark position was

$$\mathbf{X}_{j,t}^{\text{model}} = \mathbf{o}_{T,t} + R_{T,t}\left[s_T \mathbf{q}_{T,j}\right]. \tag{S14}$$

The rotations were parameterised as

$$R_{H,t} = R_z(\phi_{H,z,t})R_x(\phi_{H,x,t})R_y(\phi_{H,y,t}), \tag{S15}$$

and equivalently for the torso. The forward axis of each segment was the world-coordinate image of its local $x$-axis.

#### Individual scale estimation

Head and torso scales were estimated separately from the final trusted-camera triangulations. For each sequence, we calculated the following distances across valid frames:

$$d_{\text{eye},t} = \left\|\mathbf{X}_{L\text{eye},t} - \mathbf{X}_{R\text{eye},t}\right\|_2, \tag{S16}$$

$$d_{\text{nose},t} = \left\|\mathbf{X}_{\text{nose},t} - \frac{\mathbf{X}_{L\text{eye},t} + \mathbf{X}_{R\text{eye},t}}{2}\right\|_2, \tag{S17}$$

and

$$d_{\text{torso},t} = \left\|\mathbf{X}_{\text{neck},t} - \mathbf{X}_{\text{spine},t}\right\|_2. \tag{S18}$$

Only frames in which all landmarks required for the relevant distance had at least two retained views were used. Robust median distances were converted into scale estimates by comparison with the corresponding template distances:

$$s_{\text{eye}} = \frac{\text{median}_t(d_{\text{eye},t})}{0.0600}, \qquad s_{\text{nose}} = \frac{\text{median}_t(d_{\text{nose},t})}{\sqrt{0.0550^2 + 0.0550^2}}, \tag{S19}$$

and

$$s_T = \frac{\text{median}_t(d_{\text{torso},t})}{0.3700}. \tag{S20}$$

The head scale was set to $s_{\text{nose}}$. The eye width is the shortest landmark baseline and is therefore the most sensitive to annotation noise and limited camera coverage. It also exceeded the nose-based estimate in every sequence (by a median of 24%, range 6–89%), indicating that the annotated eye spacing is wider than the template eye width. $s_{\text{eye}}$ was therefore used only as a diagnostic. Using a single estimator in all sequences also avoids a threshold-dependent switch between estimators, which would change the head scale of a sequence by approximately 10%. The ear coordinates were subsequently scaled using this head-scale estimate.

#### Framewise pose initialisation

Triangulated landmarks were used to initialise, rather than directly define, the final pose estimate. When at least three non-collinear head landmarks were available, the scaled head template was aligned to the triangulated head landmarks using a rotation-only Kabsch alignment. This yielded an initial 3D head orientation and origin.

The torso forward axis was initialised from the directed vector from spine to neck base. A level, zero-roll torso rotation was constructed by mapping the local forward axis onto this vector. Because the torso is represented by only the spine and neck-base landmarks, roll about the spine–neck axis is not independently identifiable. This does not affect the forward axis used for the pitch and yaw analyses.

When fewer than two triangulated landmarks were available at a frame, the previous fitted pose was used only as an optimisation starting value. No temporal residual was included in this framewise fitting stage.

#### Consensus-weighted robust reprojection residual

Observation-level weighting retained useful landmarks within views that also contained outliers. For example, at frame 73 of `phantomTopRun1_20170903` (figure S8), camera 6 had an unweighted validation RMSE of 10.13 px across five observations after self-calibration. Residuals were 3.59 px for the left eye, 1.00 px for the nose, 4.24 px for the left ear, 21.71 px for the spine and 3.07 px for the neck base. The large spine residual therefore coexisted with useful head observations. These residuals alone do not identify the source of the discrepancy.

The LOO trust assessment preceded residual construction. Let $C_t^{\text{keep}}$ denote the set of cameras retained at frame $t$. If camera $c$ was rejected at that frame, it was excluded from both the final trusted-camera triangulation and the subsequent nonlinear pose optimisation. The camera–landmark consensus weighting described below was a separate procedure applied only to observations from cameras in $C_t^{\text{keep}}$.

Let $t$ index frames, $c$ cameras and $j$ anatomical landmarks, and let $L_{c,j,t}$ denote the corresponding landmark likelihood. An observation was eligible for fitting only when camera $c$ was retained by the frame-specific camera-trust mask, $L_{c,j,t} > 0.5$, and both the observed undistorted coordinate and the model projection were finite. Observations with $L_{c,j,t} \le 0.5$ were excluded.

The base image-plane residual-normalisation scales, in landmark order

{left eye, right eye, nose, left ear, right ear, spine, neck base},

were

$$\boldsymbol{\sigma}_{\text{fit}}^{\text{base}} = [3, 3, 2, 5, 5, 6, 6] \text{ pixels}. \tag{S21}$$

These scales were specified a priori as relative observation weights: smaller values gave greater weight to landmarks expected to be localised more precisely. They were not empirical standard deviations (SDs) estimated from repeated annotations.

The effective scale additionally depended on landmark likelihood:

$$\sigma_{c,j,t}^{\mathrm{eff}} = \begin{cases} \sigma_j^{\mathrm{base}}, & L_{c,j,t} \geq 0.85, \\ 3\sigma_j^{\mathrm{base}}, & 0.5 < L_{c,j,t} < 0.85. \end{cases} \tag{S22}$$

Thus, an eligible observation with intermediate likelihood remained in the fit but received one third of the residual amplitude, and hence one ninth of the initial quadratic weight, of an otherwise identical high-likelihood observation before robust and consensus weighting.

For a candidate pose $\mathbf{x}_t$, the normalised 2D reprojection residual was

$$\mathbf{e}_{c,j,t}(\mathbf{x}_t) = \frac{\pi_c\left[\mathbf{X}_j^{\mathrm{model}}(\mathbf{x}_t)\right] - \mathbf{u}_{c,j,t}^{\mathrm{obs}}}{\sigma_{c,j,t}^{\mathrm{eff}}} \in \mathbb{R}^2. \tag{S23}$$

Here, $\mathbf{X}_j^{\mathrm{model}}(\mathbf{x}_t)$ is the world-frame model-landmark position, $\pi_c$ is projection into camera $c$'s undistorted image plane, and $\mathbf{u}_{c,j,t}^{\mathrm{obs}}$ is the corresponding observed undistorted coordinate. Division by $\sigma_{c,j,t}^{\mathrm{eff}}$, measured in pixels, made the residual dimensionless.

For each valid camera–landmark observation, the magnitude of the normalised residual was

$$a_{c,j,t} = \left\|\mathbf{e}_{c,j,t}\right\|_2. \tag{S24}$$

Thus, $a_{c,j,t}$ is the radial image-plane error expressed in units of the landmark-specific uncertainty scale.

Let $n_{j,t} = |C_{j,t}|$ be the number of retained cameras observing landmark $j$. When $n_{j,t} \geq 3$, the landmark-specific consensus scale was

$$s_{j,t}^{\mathrm{cons}} = \max\left[\mathrm{median}_{c \in C_{j,t}}\left(a_{c,j,t}\right), 2\right]. \tag{S25}$$

The constant 2 is a dimensionless lower bound that prevents the consensus threshold from becoming excessively small when all cameras agree closely. The corresponding redescending consensus weight was

$$w_{c,j,t}^{\mathrm{cons}} = \begin{cases} \left[1 - \left(\dfrac{a_{c,j,t}}{4 s_{j,t}^{\mathrm{cons}}}\right)^2\right]^2, & a_{c,j,t} \leq 4 s_{j,t}^{\mathrm{cons}}, \\ 0, & a_{c,j,t} > 4 s_{j,t}^{\mathrm{cons}}. \end{cases} \tag{S26}$$

When fewer than three retained cameras observed the landmark, cross-camera consensus could not be estimated and we set $w_{c,j,t}^{\mathrm{cons}} = 1$. This weight acted at the individual camera–landmark level: a camera could therefore remain in $C_t^{\mathrm{keep}}$ while one of its landmark observations was downweighted or assigned zero weight.

A Huber-type weight provided additional protection against observations with large absolute residuals:

$$w_{c,j,t}^{\mathrm{Huber}} = \begin{cases} 1, & a_{c,j,t} \leq 3, \\ \dfrac{3}{a_{c,j,t}}, & a_{c,j,t} > 3. \end{cases} \tag{S27}$$

The threshold 3 is expressed in normalised-residual units. The residual supplied to `lsqnonlin` was

$$\widetilde{\mathbf{e}}_{c,j,t}(\mathbf{x}_t) = \sqrt{w_{c,j,t}^{\mathrm{cons}} w_{c,j,t}^{\mathrm{Huber}}}\, \mathbf{e}_{c,j,t}(\mathbf{x}_t). \tag{S28}$$

All residual magnitudes, consensus scales and weights were recomputed whenever the optimiser evaluated a new candidate pose.

Let $\mathcal{J}$ denote the complete set of head and torso model landmarks. The framewise reprojection objective was

$$\widehat{\mathbf{x}}_t = \arg\min_{\mathbf{x}_t} \sum_{j \in \mathcal{J}} \sum_{c \in C_{j,t}} \left\|\widetilde{\mathbf{e}}_{c,j,t}(\mathbf{x}_t)\right\|_2^2. \tag{S29}$$

The two components of each weighted image-plane residual were stacked into the residual vector supplied to `lsqnonlin`. For $a_{c,j,t} > 3$, the Huber-type weighting changed the contribution before consensus weighting from the quadratic value $a_{c,j,t}^2$ to the linear value $3a_{c,j,t}$, thereby limiting the influence of large residuals.

#### Bounded seed refinement and framewise nonlinear least-squares fitting

The pose vector was ordered as

$$\mathbf{x}_t = \left[\phi_{H,x}, \phi_{H,y}, \phi_{H,z}, x_H, y_H, z_H, \phi_{T,x}, \phi_{T,y}, \phi_{T,z}, x_T, y_T, z_T\right]^{\mathsf{T}}. \tag{S30}$$

Two bounded nonlinear least-squares stages were used: an initial 3D seed-refinement stage, followed by the main multiview reprojection fit (figure S2, algorithm S3).

**3D seed refinement.** The pose constructed from trusted-camera triangulation was first refined by minimising the unweighted 3D differences between the rigid-body model landmarks and the corresponding seed landmarks. At frames after the first, landmarks unavailable in the current triangulation were replaced by their positions under the preceding fitted pose for this initialisation step only.

For the head seed, the first and second rotation parameters were bounded within $\pm\pi/5$ of their initial values, the third rotation parameter within $\pm\pi/2$, and each origin coordinate within $\pm 0.5$ m. For the torso seed, the first rotation parameter was bounded absolutely by $[-\pi/5, \pi/5]$, the second within $\pm\pi/5$ of its initial value, the third within $\pm\pi/2$, and each origin coordinate within $\pm 0.5$ m.

For the first frame, this seed-refinement solve used `StepTolerance` and `OptimalityTolerance` of $10^{-10}$, with maximum limits of $5 \times 10^4$ function evaluations and $5 \times 10^3$ iterations. For subsequent frames, the corresponding settings were $10^{-8}$, $10^4$ function evaluations and $10^3$ iterations. If fewer than two trusted triangulated landmarks were available, seed refinement was skipped and the preceding fitted pose was used. If a subsequent-frame seed-refinement solve raised a solver error, the preceding fitted pose was likewise used as the fallback initialisation.

**Main reprojection fit.** The main fit used global bounds shared by the framewise and bidirectional reconstruction stages. For both head and torso,

$$\phi_x \in [-\pi/5, \pi/5], \qquad \phi_y \in [-\pi/3, \pi/3], \qquad \phi_z \in [-\pi, \pi]. \tag{S31}$$

Let $C_{c,x}$, $C_{c,y}$ and $C_{c,z}$ denote the world-coordinate components of the calibrated camera positions. The head and torso origins were bounded identically:

$$\begin{aligned} x &\in \left[\min_c C_{c,x} - 5,\ \max_c C_{c,x} + 5\right] \text{ m}, \\ y &\in \left[\min_c C_{c,y} - 2,\ \max_c C_{c,y} + 2\right] \text{ m}, \\ z &\in \left[\min_c C_{c,z} - 0.5,\ \max_c C_{c,z} + 1.5\right] \text{ m}. \end{aligned} \tag{S32}$$

Every starting pose was clamped componentwise to these bounds before optimisation.

The main framewise fit used MATLAB's `lsqnonlin` with `Display` set to `off`, `StepTolerance` and `OptimalityTolerance` set to $10^{-4}$, and maximum limits of $10^5$ function evaluations and $10^5$ iterations. All solver options not explicitly specified retained their MATLAB `R2026a` defaults.

Frame 1 was optimised once from its refined multiview seed. For each later frame, the main solver was normally run from two bounded starting poses: the current frame's refined multiview seed and the preceding frame's fitted pose. When the current multiview evidence was too sparse, or when seed refinement failed, the preceding pose supplied the current initialisation and the two starts could therefore coincide.

Each main solve minimised only the consensus-weighted robust multiview reprojection residual defined above; no temporal residual was included. Among the candidate solutions, the pose with the smallest value of `lsqnonlin`'s returned `resnorm`, equal to the sum of squared components of the weighted residual vector, was retained:

$$\widehat{\mathbf{x}}_t = \underset{\mathbf{x} \in \{\mathbf{x}_t^{(1)}, \mathbf{x}_t^{(2)}\}}{\arg\min} \ \|\widetilde{\mathbf{e}}_t(\mathbf{x})\|_2^2. \tag{S33}$$

The preceding pose was used only as an alternative initial value and did not introduce temporal coupling. Temporal regularisation was added only by the offline bidirectional stage in S4.

**Algorithm S3** Framewise consensus-weighted robust nonlinear least-squares pose fitting.

**Require:** Trusted triangulated landmarks, undistorted image observations, camera masks, calibrated camera models and scaled rigid-body template
**Ensure:** Framewise head–torso poses

**for** each frame $t$ **do**
  Obtain valid triangulated head and torso landmarks
  **if** at least three suitable head landmarks are available **then**
    Initialise head orientation and origin using Kabsch alignment
  **else**
    Use the previous fitted pose as the head initialisation
  **end if**
  **if** spine and neck base are available **then**
    Initialise torso forward axis from the spine-to-neck vector
  **else**
    Use the previous fitted pose as the torso initialisation
  **end if**
  Construct candidate starts from the triangulation-based pose and, for $t > 1$, the preceding fitted pose
  **for** each candidate starting pose **do**
    **for** each retained camera $c$ and visible landmark $j$ **do**
      Transform the template landmark into world coordinates
      Project the landmark into camera $c$
      Calculate the normalised residual $\mathbf{e}_{c,j,t}$
    **end for**
    **for** each landmark observed by at least three retained cameras **do**
      Calculate camera–landmark consensus weights $w^{\text{cons}}_{c,j,t}$
    **end for**
    Apply Huber weights to the normalised residuals
    Stack weighted residuals and optimise the pose using `lsqnonlin`
  **end for**
  Retain the candidate with the lowest robust residual norm
**end for**

### Supplementary Note S4. Offline bidirectional CV smoothing

To reduce isolated fluctuations caused by annotation noise or changes in camera visibility, we refitted the interior frames with temporal regularisation. Each update retained the original consensus-weighted robust reprojection residual and added a constant-velocity (CV) prior based on the neighbouring poses.

#### Pose representation and CV midpoint target

Let $\mathbf{p}_f$ denote the 12-parameter pose defined in Supplementary Note S3, with segment origin $\mathbf{x}_{k,f}$ and rotation $R_{k,f}$ for $k \in \{H, T\}$. Orientations were regularised using rotation matrices on $SO(3)$ to avoid Euler-angle wrapping. Below, $\log(R_a^{\mathsf{T}} R_b)$ denotes the three-component rotation vector taking $R_a$ to $R_b$, and exp maps this vector back to a rotation matrix.

For an interior frame $f$, the CV targets were the position and orientation midpoints between its neighbours:

$$\mathbf{x}^{\text{target}}_{k,f} = \frac{\mathbf{x}_{k,f-1} + \mathbf{x}_{k,f+1}}{2}, \qquad R^{\text{target}}_{k,f} = R_{k,f-1} \exp\left[\frac{1}{2} \log\left(R^{\mathsf{T}}_{k,f-1} R_{k,f+1}\right)\right]. \tag{S34}$$

These targets correspond to locally constant linear and angular velocity.

#### Temporal residuals

For a candidate pose at frame $f$, the deviation of head orientation from its midpoint target was measured by

$$\mathbf{r}_{H,f}^{\text{rot}} = \frac{\log\left[\left(R_{H,f}^{\text{target}}\right)^{\mathsf{T}} R_{H,f}\right]}{\sigma_{\text{rot}}}, \tag{S35}$$

and the corresponding positional deviation was

$$\mathbf{r}_{H,f}^{\text{pos}} = \frac{\mathbf{x}_{H,f} - \mathbf{x}_{H,f}^{\text{target}}}{\sigma_{\text{pos}}}. \tag{S36}$$

Equivalent residuals, $\mathbf{r}_{T,f}^{\text{rot}}$ and $\mathbf{r}_{T,f}^{\text{pos}}$, were calculated for the torso.

We used $\sigma_{\text{rot}} = 2°$ and $\sigma_{\text{pos}} = 0.015$ m to set the strength of temporal regularisation.

The temporal residual vector was

$$\mathbf{r}_f^{\text{temp}} = \left[\mathbf{r}_{H,f}^{\text{rot}}; \mathbf{r}_{T,f}^{\text{rot}}; \mathbf{r}_{H,f}^{\text{pos}}; \mathbf{r}_{T,f}^{\text{pos}}\right]. \tag{S37}$$

Each component was transformed using a Huber pseudo-residual with tuning constant $c = 3$. For a scaled residual $r$, this transformation was

$$\widetilde{r} = \begin{cases} r, & |r| \le c, \\ \operatorname{sign}(r)\sqrt{2c|r| - c^2}, & |r| > c. \end{cases} \tag{S38}$$

The resulting penalty was quadratic for small deviations and linear for large deviations, allowing image-supported departures from the CV target.

#### Framewise nonlinear update

At each interior frame, `lsqnonlin` minimised

$$\hat{\mathbf{p}}_f = \arg\min_{\mathbf{p}} \left\| \left[\mathbf{r}_f^{\text{data}}(\mathbf{p}); \widetilde{\mathbf{r}}_f^{\text{temp}}(\mathbf{p})\right] \right\|_2^2. \tag{S39}$$

The data residual was the consensus-weighted robust reprojection residual from Supplementary Note S3, with the same LOO camera mask. Thus, views rejected during camera-trust filtering also remained excluded from smoothing.

#### Alternating forward and backward passes

Starting from the robust framewise poses $\{\mathbf{p}_f^{(0)}\}_{f=1}^{N}$, we performed up to six passes, alternating forward through frames $2, \ldots, N-1$ and backward through $N-1, \ldots, 2$. Each midpoint used the most recently available neighbours, including poses already updated in the current pass.

Each update used two initial poses: the estimate from the preceding pass and the original framewise fit. We retained the solution with the smaller objective value. The first and final frames remained at their framewise estimates because they lacked two neighbours. Iteration stopped after six passes or when the median rotational change between successive passes fell below 0.03°.

**Algorithm S4** Offline bidirectional CV smoothing.

**Require:** Framewise poses $\{\mathbf{p}_f^{(0)}\}_{f=1}^{N}$; original multiview landmark observations; camera-consensus masks
**Ensure:** Smoothed poses $\{\mathbf{p}_f\}_{f=1}^{N}$

$\mathbf{p}_f \leftarrow \mathbf{p}_f^{(0)}$ for all frames $f$
**for** $k = 1$ to 6 **do**
  $\mathbf{p}^{\text{old}} \leftarrow \mathbf{p}$
  $\mathbf{p}^{\text{new}} \leftarrow \mathbf{p}^{\text{old}}$
  **if** $k$ is odd **then**
    $\mathcal{F} \leftarrow [2, 3, \ldots, N-1]$
  **else**
    $\mathcal{F} \leftarrow [N-1, N-2, \ldots, 2]$
  **end if**
  **for all** $f \in \mathcal{F}$ **do**
    Construct head and torso midpoint targets from $\mathbf{p}_{f-1}^{\text{new}}$ and $\mathbf{p}_{f+1}^{\text{new}}$
    Form the combined data and Huberised temporal residual vector $\mathbf{r}_f(\mathbf{p})$
    Optimise from $\mathbf{p}_f^{\text{old}}$ using `lsqnonlin`
    Optimise from $\mathbf{p}_f^{(0)}$ using `lsqnonlin`
    Retain the solution with the smaller final residual norm as $\mathbf{p}_f^{\text{new}}$
  **end for**
  $\mathbf{p} \leftarrow \mathbf{p}^{\text{new}}$
  **if** median rotational change $< 0.03°$ **then**
    **break**
  **end if**
**end for**
**return p**

The smoother uses both preceding and subsequent observations. Its output therefore describes reconstructed kinematics and cannot establish causal response latency.

### Supplementary Note S5. Monte Carlo propagation of reconstruction uncertainty

We propagated uncertainty in landmark annotations, lure position and anatomical templates through $M = 500$ joint reconstructions per pursuit. Each draw perturbed the inputs for a complete sequence, refitted head and torso poses, applied bidirectional smoothing and independently re-triangulated the lure. The resulting variation measures sensitivity to the input distributions specified below.

#### Image-plane landmark uncertainty

Let $\mathbf{u}_{jct}$ denote the undistorted image coordinate of anatomical landmark $j$ in camera $c$ at frame $t$. In Monte Carlo draw $m$, a perturbed coordinate was generated as

$$\mathbf{u}_{jct}^{(m)} = \mathbf{u}_{jct} + \boldsymbol{\epsilon}_{jct}^{(m)}, \qquad \boldsymbol{\epsilon}_{jct}^{(m)} \sim \mathcal{N}_2\left(\mathbf{0}, \frac{s_j^2}{2}\mathbf{I}_2\right), \tag{S40}$$

where $s_j$ is the RMS magnitude of the 2D perturbation. Consequently, the SD in each image coordinate was $s_j/\sqrt{2}$. Perturbations were independent between image coordinates, landmarks, cameras and frames. These perturbation scales were assumed because repeated independent annotations were unavailable.

Original landmark visibility and the nominal LOO camera-trust mask were held fixed. Invisible landmarks remained excluded. Within retained views, camera–landmark consensus and Huber-type weights were recalculated for each candidate pose using the perturbed observations.

The lure perturbation was applied to the original distorted image-plane lure annotations. For each visible lure annotation,

$$\mathbf{v}_{ct}^{(m)} = \mathbf{v}_{ct} + \boldsymbol{\eta}_{ct}^{(m)}, \qquad \boldsymbol{\eta}_{ct}^{(m)} \sim \mathcal{N}_2\left(\mathbf{0}, 2^2\mathbf{I}_2\right) \text{ pixels.} \tag{S41}$$

The perturbed annotations were undistorted and re-triangulated using the corresponding camera model. At least two visible camera observations were required. Lure perturbations were independent between coordinates, cameras and frames. This represents uncertainty in the annotated lure point; the lure's physical extent and centreline were not modelled.

#### Anatomical-template uncertainty

The rigid-body template was parameterised by ten signed dimensions,

$$\mathbf{L} = [L_{\text{eye},y}, L_{\text{nose},x}, L_{\text{nose},z}, L_{\text{ear},x}, L_{\text{ear},y}, L_{\text{ear},z}, L_{\text{neck},x}, L_{\text{neck},z}, L_{\text{spine},x}, L_{\text{spine},z}]. \tag{S42}$$

Before animal-specific scaling, their nominal values were

$$\mathbf{L}_{\text{template}} = [0.030,\ 0.055,\ -0.055,\ -0.0725,\ 0.0608,\ 0.0441,\ 0.040,\ 0.100,\ -0.330,\ 0.100]\ \text{m}. \tag{S43}$$

Head dimensions were multiplied by the estimated head scale for the individual, whereas neck and spine dimensions were multiplied by the estimated torso-length scale.

For dimension $k$, the value in Monte Carlo draw $m$ was

$$L_k^{(m)} = L_{k,0} + \tau_k z_k^{(m)}, \qquad z_k^{(m)} \sim \mathcal{N}(0, 1), \tag{S44}$$

where $L_{k,0}$ is the animal-scaled nominal dimension and $\tau_k$ is its SD. For all non-ear dimensions,

$$\tau_k = 0.08\,|L_{k,0}|. \tag{S45}$$

The ear dimensions used axis-specific SDs derived from the observed inter-dataset variation in head-local ear position:

$$\tau_{\text{ear},x} = 0.011\ \text{m}, \qquad \tau_{\text{ear},y} = 0.005\ \text{m}, \qquad \tau_{\text{ear},z} = 0.009\ \text{m}. \tag{S46}$$

Each draw used one template throughout the pursuit. Paired landmarks shared sampled dimensions with opposite signs on the bilateral axis, preserving bilateral symmetry. Gaussian dimension distributions were untruncated; no dimension changed sign in the successful production draws.

#### Reconstruction within each Monte Carlo draw

Each perturbed sequence was reconstructed using the bounded pose fit in Supplementary Note S3, retaining its residual scales, robust loss and weighting rules. Where available, the nominal poses at the current and preceding frames supplied two starting values; the solution with the smaller objective was retained. The sequence then underwent six alternating forward and backward smoothing passes with the midpoint scales and Huber tuning defined in Supplementary Note S4. Target-relative pitch, yaw and 3D errors were calculated from the resulting head, torso and lure trajectories.

Camera calibration, synchronisation offsets, binary landmark visibility, animal scale estimates, analysed frame ranges and nominal hard camera-trust masks were fixed across draws. Their uncertainty is therefore excluded from the reported reconstruction intervals.

#### Random-number generation and reconstruction diagnostics

We used master seed 1 and 500 indexed `mrg32k3a` streams, one per draw, giving identical random inputs in serial and parallel execution. Streams were reinitialised for each pursuit, so matching draw indices across pursuits identify pseudorandom streams, not biological replicates.

For each draw, we stored the sampled template dimensions, perturbed lure trajectory, framewise initial pose, final CRBR pose, derived angles, reprojection diagnostics, convergence diagnostics and success status. A draw was considered successful when its final pose trajectory was finite and it produced finite derived angular values. All 500 draws were successful for every pursuit included in the reported analyses.

##### Monte Carlo summaries

For a signed angular quantity $q(t)$, let $q_0(t)$ be its value in the nominal reconstruction and $q_m(t)$ its value in Monte Carlo draw $m$. To avoid errors at the angular branch cut, summaries were calculated from nominal-centred wrapped deviations,

$$d_m(t) = \mathrm{wrap}_{[-\pi,\pi)}\left[q_m(t) - q_0(t)\right] . \tag{S47}$$

The reported Monte Carlo median was

$$\widetilde{q}(t) = q_0(t) + \mathrm{median}_m\left[d_m(t)\right] , \tag{S48}$$

and the pointwise 95% reconstruction interval was

$$\mathcal{I}_{0.95}(t) = q_0(t) + \left[Q_{0.025}\{d_m(t)\}, Q_{0.975}\{d_m(t)\}\right] , \tag{S49}$$

where $Q_p$ is the empirical $p$th quantile across draws. Intervals apply to individual time points rather than simultaneously to the full trajectory.

For a categorical yaw state $s$, reconstruction support at frame $t$ was calculated as

$$\widehat{P}_t(s) = \frac{1}{500}\sum_{m=1}^{500}\mathbb{I}\left[S_m(t) = s\right] . \tag{S50}$$

A single state was considered supported when $\widehat{P}_t(s) \geq 0.95$, corresponding to at least 475 of the 500 reconstructions. Frames not reaching this threshold were retained as ambiguities between the smallest set of states whose cumulative support reached 0.95, or as unresolved when support was distributed more broadly.

These summaries quantify reconstruction sensitivity under the specified perturbations. Variation among pursuits and animals was assessed separately using the aggregation procedures in Supplementary Note S6.

#### Supplementary Note S6. Pitch- and yaw-analysis details

Calculations were performed separately within each pursuit on its retained analysis frames. Pitch analyses used 16 frontal and lateral pursuits (1,590 frames from 5 cheetahs), excluding both close pass-by pursuits. Yaw geometry and state classification used all 18 pursuits (1,708 frames from 6 cheetahs). The absolute yaw-error ECDFs used seven lateral and close pass-by pursuits (628 frames from 4 cheetahs), and signed yaw-error histograms used five lateral pursuits (510 frames from 3 cheetahs).

Pitch ECDFs, centred distributions and transmission estimates used nominal reconstruction trajectories. Uncertainty-aware yaw classification and run-level reconstruction intervals were calculated independently within each joint Monte Carlo draw before aggregation. Descriptive yaw plots and ECDFs used framewise median-based point estimates as specified below; these summaries were not substituted for joint trajectories in state-support calculations.

##### Stride-averaged running speed

To describe running speed, we used the original AcinoSet full-trajectory estimation (FTE) sequences for all 18 pursuits, taking each sequence's full available interval. We identified complete strides from repeated forward excursions of the right forelimb. Specifically, we projected the ankle's position relative to the shoulder onto the horizontal spine-to-neck axis and detected successive peaks, with a minimum prominence of 0.10 m and spacing of 0.2 s. Intervals longer than 0.7 s were excluded. In one short sequence without a complete peak-to-peak cycle, we used successive backward extrema of the same limb. These boundaries mark repeated limb positions within the gait cycle, without requiring foot-contact annotations.

We defined the trunk centre as the mean position of both shoulders and both hips. For each stride, we summed its frame-to-frame horizontal displacements and divided this path length by the stride duration:

$$\bar{v}_k = \frac{\sum_{n=a_k}^{b_k-1} \left\| \mathbf{p}_{n+1}^{xy} - \mathbf{p}_n^{xy} \right\|_2}{(b_k - a_k)\,\Delta t}, \tag{S51}$$

where $\mathbf{p}_n^{xy}$ is the horizontal trunk-centre position at frame $n$, $a_k$ and $b_k$ are the boundary frames of stride $k$, and $\Delta t$ is the FTE time step (1/90 or 1/120 s). The reported range and median pool the resulting 41 stride speeds with equal weight per stride. Limb-cycle traces and representative video sequences were checked visually.

#### Pitch-error ECDFs

Let $\alpha_{j,n}$ and $\beta_{j,n}$ denote the valid head–target and torso–target pitch errors, in degrees, for frame $n$ of pursuit $j$. The pursuit-specific ECDFs of absolute error were

$$F_{\alpha,j}(x) = \frac{1}{N_j} \sum_{n=1}^{N_j} \mathbf{1}\big(|\alpha_{j,n}| \le x\big), \qquad F_{\beta,j}(x) = \frac{1}{N_j} \sum_{n=1}^{N_j} \mathbf{1}\big(|\beta_{j,n}| \le x\big). \tag{S52}$$

The ECDFs were evaluated on a common grid at 0.25° intervals.

The primary run-equal ECDFs were

$$\overline{F}_\alpha(x) = \frac{1}{J} \sum_{j=1}^{J} F_{\alpha,j}(x), \qquad \overline{F}_\beta(x) = \frac{1}{J} \sum_{j=1}^{J} F_{\beta,j}(x), \tag{S53}$$

so every pursuit contributed equally irrespective of its number of frames. SEs were calculated as the sample SD of pursuit-level ECDF values divided by $\sqrt{J}$. For the animal-equal sensitivity analysis, ECDFs were first averaged equally across pursuits belonging to the same animal and the resulting animal-level ECDFs were then averaged equally across animals. Proportions within 5°, 10° and 15° were obtained by evaluating the corresponding pursuit-level ECDF at those thresholds before applying the same aggregation.

#### Within-pursuit pitch variability

We calculated the sample SD of world-frame head and torso pitch within each pursuit, using the retained pitch-analysis frames without additional smoothing. The 16 pursuit-level SDs were summarised by their mean and SE. These values describe world-frame pitch variability; target-relative precision was assessed from errors $\alpha$ and $\beta$ below.

#### Run-centred distributions and FWHM

Within-pursuit precision was evaluated after removing each pursuit's mean pointing offset:

$$\widetilde{\alpha}_{j,n} = \alpha_{j,n} - \overline{\alpha}_j, \qquad \widetilde{\beta}_{j,n} = \beta_{j,n} - \overline{\beta}_j. \tag{S54}$$

Each centred distribution was represented by unsmoothed bin proportions using 5°-wide bins centred at multiples of 5°. If $x_k$ is a bin centre and $B_k = [x_k - 2.5°, x_k + 2.5°)$, the pursuit-level bin proportions were

$$p_{\alpha,j}(x_k) = \frac{\sum_{n=1}^{N_j} \mathbf{1}(\widetilde{\alpha}_{j,n} \in B_k)}{\sum_{n=1}^{N_j} \mathbf{1}(\widetilde{\alpha}_{j,n} \in \mathcal{B})}, \tag{S55}$$

where $\mathcal{B}$ is the union of the histogram bins. Thus, each run is normalised by its number of samples within the histogram edges; the final bin includes its upper edge. The torso distribution is defined analogously. The plotted distributions were the run-equal means

$$\overline{p}_\alpha(x_k) = \frac{1}{J}\sum_{j=1}^{J} p_{\alpha,j}(x_k), \qquad \overline{p}_\beta(x_k) = \frac{1}{J}\sum_{j=1}^{J} p_{\beta,j}(x_k). \tag{S56}$$

The outermost bin centres were symmetric about zero. Their magnitude was the greater of 15° and the pooled 99th percentile of absolute centred errors, rounded up to the next multiple of 5°.

For either mean distribution $\overline{p}(x_k)$, let $k^\star$ be the location of its maximum and let

$$p_{1/2} = \frac{1}{2}\overline{p}(x_{k^\star}). \tag{S57}$$

Starting from $k^\star$, the algorithm searched separately towards smaller and larger errors for the adjacent bins bracketing $p_{1/2}$. The left and right crossing positions, $x_L$ and $x_R$, were obtained by linear interpolation between the corresponding bin centres. Full width at half maximum was then

$$\text{FWHM} = x_R - x_L. \tag{S58}$$

FWHM therefore describes the centred, run-equal discrete distribution. For animal-equal sensitivity, we first averaged pursuit distributions within each animal, then applied the same procedure to their equally weighted mean.

#### Construction of nominal 50-ms pitch changes

World-frame head pitch $\theta_H$, world-frame torso pitch $\theta_T$ and head-to-target elevation $\lambda_H$ were first divided into contiguous valid segments. A pair was never formed across an excluded frame or a break in absolute frame number. Within each segment, each angular trajectory was unwrapped and filtered with a centred moving median. For sampling frequency $f_s$, the nominal smoothing width was

$$w = \max[3, \text{round}(0.050 f_s)] \tag{S59}$$

samples and was increased by one when necessary to obtain an odd width. At a segment boundary, the available portion of this window was used.

The change lag was

$$L = \max[1, \text{round}(0.050 f_s)] \tag{S60}$$

frames. Consequently, "50 ms" denotes the nominal window; its realised duration in a sequence was $L/f_s$. For every admissible starting sample $n$, overlapping finite changes were constructed as

$$\begin{aligned}
\Delta\theta_H(n) &= \widetilde{\theta}_H(n+L) - \widetilde{\theta}_H(n),\\
\Delta\theta_T(n) &= \widetilde{\theta}_T(n+L) - \widetilde{\theta}_T(n),\\
\Delta\lambda_H(n) &= \widetilde{\lambda}_H(n+L) - \widetilde{\lambda}_H(n),\\
\Delta q(n) &= \Delta\theta_H(n) - \Delta\theta_T(n),
\end{aligned} \tag{S61}$$

where tildes denote the unwrapped, moving-median-filtered trajectories. Changes were converted to degrees before model fitting.

Within each pursuit, ordinary least squares with an intercept was used to fit

$$\Delta\theta_H = \beta_0 + g_\lambda \Delta\lambda_H + g_T \Delta\theta_T + \varepsilon. \tag{S62}$$

At least 20 valid change pairs were required. The equivalent counter-rotation model follows from $\Delta q = \Delta\theta_H - \Delta\theta_T$:

$$\Delta q = \beta_0 - g_D \Delta\theta_T + g_\lambda \Delta\lambda_H + \varepsilon, \qquad g_D = 1 - g_T. \tag{S63}$$

Coefficients were estimated separately for each pursuit. For the animal-equal summary, pursuit coefficients were first averaged within each animal; means and SEs were then calculated across animals. The run-equal visualisation instead gave every pursuit equal weight.

#### Local target-path tangent for yaw classification

The descriptive yaw-state classification required the local direction of target travel, not a future-position prediction. The tangent estimator used a centred path window and was therefore not treated as a causal sensorimotor model.

For target horizontal position $\mathbf{L}_{xy}(t) = [L_x(t), L_y(t)]^\top$, the initial half-window at sampling frequency $f_s$ was

$$h_0 = \max[2, \mathrm{round}(0.100 f_s/2)]\,, \tag{S64}$$

and the maximum half-window was

$$h_{\max} = \max[h_0, \mathrm{round}(0.300 f_s/2)]\,. \tag{S65}$$

For each frame $i$, valid target samples from $i - h$ to $i + h$, clipped at the sequence boundaries, were fitted by ordinary least squares:

$$L_x(t_k) = a_x + v_x(t_k - t_i) + \epsilon_{x,k}, \qquad L_y(t_k) = a_y + v_y(t_k - t_i) + \epsilon_{y,k}. \tag{S66}$$

The candidate oriented tangent was

$$\widehat{\mathbf{u}}_i = \frac{[v_x, v_y]^\top}{\sqrt{v_x^2 + v_y^2}}. \tag{S67}$$

Starting at $h_0$, the window was expanded one frame at a time until it satisfied all of the following:

$$\begin{gathered}\Delta_i = \sqrt{\left(\max_k L_x - \min_k L_x\right)^2 + \left(\max_k L_y - \min_k L_y\right)^2} \geq 0.05\ \mathrm{m},\\ \mathcal{L}_i = \frac{\sigma_1^2}{\sigma_1^2 + \sigma_2^2} \geq 0.65, \qquad \sqrt{v_x^2 + v_y^2} > 10^{-9},\end{gathered} \tag{S68}$$

where $\sigma_1 \geq \sigma_2$ are the singular values of the centred horizontal target coordinates. If no window at a frame met these criteria, the oriented tangent from the nearest-in-time frame with a reliable target direction was used. Behavioural state labels were not borrowed from neighbouring frames.

#### Ahead, aligned and lagged yaw states

For each nominal or Monte Carlo reconstruction, raw head–target yaw error was

$$\gamma_i = \mathrm{wrap}_{[-\pi,\pi)}\left[\psi_H(t_i) - \lambda_{H,\mathrm{yaw}}(t_i)\right]. \tag{S69}$$

Before classification, $\gamma_i$ was unwrapped and passed through a Hampel filter using three samples on either side of the current sample and a threshold of three scaled median absolute deviations. A flagged value was replaced by the local median, and the result was wrapped back to $[-\pi, \pi)$. The filtered head direction used for path projection was reconstructed by adding the filtered error to the head-to-target bearing.

Let $\mathbf{H}_{xy,i}$ be the head origin, let $\mathbf{d}_{H,i}$ be the unit horizontal head direction after this filtering step, and let

$$R_{H,i} = \left\|\mathbf{L}_{xy,i} - \mathbf{H}_{xy,i}\right\|_2 \tag{S70}$$

be the current horizontal head–target range. The head-forward ray was evaluated at this range:

$$\mathbf{G}_i = \mathbf{H}_{xy,i} + R_{H,i}\mathbf{d}_{H,i}. \tag{S71}$$

Its signed displacement along the oriented target path was

$$\ell_i = \left(\mathbf{G}_i - \mathbf{L}_{xy,i}\right)^{\mathsf{T}} \widehat{\mathbf{u}}_i. \tag{S72}$$

For classification, we used the dimensionless projection $q_i = \ell_i / R_{H,i}$, equivalently $(\mathbf{d}_{H,i} - \widehat{\mathbf{r}}_i)^{\mathsf{T}}\widehat{\mathbf{u}}_i$, where $\widehat{\mathbf{r}}_i$ is the unit head-to-target bearing. The state rule was

$$s_i = \begin{cases} \text{aligned}, & |\gamma_i| \le \theta_{\text{align}}, \\ \text{ahead}, & |\gamma_i| > \theta_{\text{align}} \text{ and } q_i > 10^{-10}, \\ \text{lagged}, & |\gamma_i| > \theta_{\text{align}} \text{ and } q_i < -10^{-10}, \\ \text{direction ambiguous}, & \text{otherwise}. \end{cases} \tag{S73}$$

The primary alignment gate was $\theta_{\text{align}} = 10°$. The ahead and lagged labels describe the displacement of an equal-range point along head direction relative to the current target, projected onto local target travel. The numerical tolerance retains indeterminate directions as ambiguous. Here $q_i$ is the directional projection denoted $s$ in the main Methods, not head-on-torso pitch. Within each draw, the travel-directed yaw offset was $\operatorname{sgn}(q_i)|\gamma_i|$; the alignment gate took priority when assigning states. The point estimates used in main-text figure 4d,e are described below.

The complete tangent estimation and classification were repeated independently for all $M = 500$ joint pose–target reconstruction draws. Framewise support for state $s$ was

$$p_{i,s} = \frac{1}{M}\sum_{m=1}^{M} \mathbf{1}\left(s_i^{(m)} = s\right). \tag{S74}$$

A frame was assigned an ahead, aligned or lagged state when its support was at least 0.95; otherwise it was reported as unresolved. State prevalence was first calculated within each pursuit, including unresolved frames in the denominator, and then averaged equally across pursuits within each context. Sensitivity analyses repeated classification at 5° and 15° while retaining the 0.95 support criterion. Figure S14 shows the 5° and 15° comparisons by context; the primary 10° analysis is shown in main-text figure 4f.

#### Descriptive yaw distributions and geometry summaries

For main-text figure 4d,e, the framewise travel-directed offset was

$$\gamma_{\parallel,i}^{\text{point}} = \operatorname{sgn}\left[\operatorname{median}_m \ell_i^{(m)}\right] \left|\operatorname{median}_m \gamma_i^{(m)}\right|, \tag{S75}$$

using filtered yaw errors and equal-range projections from the same reconstruction draws. This descriptive point estimate is distinct from the per-draw state classification. Horizontal head–target distance and the angle between target travel and head-to-target bearing were likewise summarised by framewise medians. Geometry scatter plots show the median of these framewise estimates within each pursuit. For 95% reconstruction intervals, the pursuit median was calculated separately within each joint draw and the 2.5th and 97.5th percentiles were taken across draws.

For the descriptive curves in main-text figure 4e, we estimated a Gaussian kernel density within each pursuit using a 5° bandwidth, then averaged densities equally within each context. Pointwise SEs were calculated across pursuit densities. Kernel smoothing affected only these descriptive curves; state assignments used the per-draw classification above.

##### Yaw-error ECDFs and uncentred signed distributions

For main-text figure 4b, each retained frame contributed the absolute value of its wrapped Monte Carlo-median head–target or torso–target yaw error. The seven pursuit-specific ECDFs were averaged equally. For figure S13, pursuit ECDFs were first averaged within each cheetah, followed by equal weighting of the four animals. We calculated pointwise SEs across pursuits or animals, respectively. The fraction with $|\gamma| < |\delta|$ was calculated by pooling the 628 matched frames; this pooled fraction is distinct from the run-equal ECDFs.

For main-text figure 4c, the five lateral pursuits contributed wrapped Monte Carlo-median signed errors, without run-centring, taking absolute values, or changing signs according to target travel. Errors were binned into 5°-wide bins, with edges covering all 510 frames. Bin counts were divided by the number of retained frames in each pursuit, then averaged equally across pursuits, with SEs calculated across the five runs. These histograms retain within-pursuit variation and between-pursuit pointing offsets, whereas the densities in figure 4e use travel-directed errors.

##### Closing-speed estimation

To examine continued approach during head–torso yaw separation, we estimated closing speed from horizontal torso–target range. With $\mathbf{T}_{xy}(t)$ denoting the horizontal torso origin, this range was

$$R_T(t) = \left\|\mathbf{L}_{xy}(t) - \mathbf{T}_{xy}(t)\right\|_2 . \tag{S76}$$

Closing speed was defined as $c(t) = -\dot{R}_T(t)$. Its implementation used a causal trailing quadratic fit. For each current sample $i$, the fitting window contained

$$n_c = \max[4, \mathrm{round}(0.100 f_s)] \tag{S77}$$

samples, including the current sample. With $u_k = t_k - t_i \le 0$, the coefficients were obtained from

$$\left(\widehat{b}_{0,i}, \widehat{b}_{1,i}, \widehat{b}_{2,i}\right) = \underset{b_0,b_1,b_2}{\arg\min} \sum_{k=i-n_c+1}^{i} \left[R_T(t_k) - b_0 - b_1 u_k - b_2 u_k^2\right]^2 . \tag{S78}$$

The endpoint estimate was

$$c(t_i) = -\widehat{b}_{1,i}, \tag{S79}$$

so $c > 0$ denotes decreasing range. No estimate was returned for the first $n_c - 1$ samples or when any range value in the fitting window was non-finite.

The descriptive closure-versus-orientation plot used nominal framewise $c(t_i)$ and absolute head–torso yaw separation

$$|\eta_{\mathrm{HT}}(t_i)| = \left|\mathrm{wrap}_{[-\pi,\pi)}\left[\theta_{H,\mathrm{yaw}}(t_i) - \theta_{T,\mathrm{yaw}}(t_i)\right]\right| . \tag{S80}$$

The plot was restricted to the 4 steady-lateral pursuits with net range closure; the pursuit in which range increased overall was excluded from this particular summary.

#### Supplementary Note S7. Causal kinematic estimation, cross-validated yaw prediction, and spatial-lead calculations

We compared candidate aiming rules using held-out pursuits and described head orientation in target-path coordinates. The model comparison and spatial-lead analysis used the procedures below.

##### Candidate-model cohort

The comparison used the five lateral pursuits over their full retained analysis windows (510 frames). Requiring 100 ms of preceding observations for target-velocity estimation left 456 matched evaluation frames. Each model used the same frames in the nominal reconstruction and all 500 joint reconstruction draws. All aiming directions were calculated in the horizontal world $x$–$y$ plane.

#### Causal endpoint-velocity estimation

To estimate local target motion without using future observations, we fitted a quadratic to its horizontal position over a trailing window. For sampling frequency $f_s$, the window contained

$$n_W = \max\{3, \mathrm{round}(0.100 f_s) + 1\} \tag{S81}$$

samples. At endpoint $i$, the sample times were $u_{i\ell} = (\ell - i)/f_s$, for $\ell = i - n_W + 1, \ldots, i$. The least-squares fit was

$$(\widehat{\boldsymbol{\beta}}_{0,i}, \widehat{\boldsymbol{\beta}}_{1,i}, \widehat{\boldsymbol{\beta}}_{2,i}) = \underset{\boldsymbol{\beta}_0,\boldsymbol{\beta}_1,\boldsymbol{\beta}_2}{\arg\min} \sum_{\ell=i-n_W+1}^{i} \left\| \mathbf{L}_{xy,\ell} - (\boldsymbol{\beta}_0 + \boldsymbol{\beta}_1 u_{i\ell} + \boldsymbol{\beta}_2 u_{i\ell}^2) \right\|_2^2, \tag{S82}$$

and the endpoint velocity was $\widehat{\mathbf{v}}_{L,i} = \widehat{\boldsymbol{\beta}}_{1,i}$. Fits restarted after gaps in accepted frames and required a complete, finite history. The subsequent forward extrapolation assumed constant velocity; the history fit estimated that velocity locally.

#### Candidate directions and shared evaluation frames

We compared current-target alignment with constant angular lead, constant-velocity extrapolation and constant spatial lead. With horizontal head-to-target displacement $\mathbf{r} = \mathbf{L}_{xy} - \mathbf{H}_{xy}$, bearing $\mathcal{B}(\mathbf{r}) = \mathrm{atan2}(r_y, r_x)$, and $s = \mathrm{sgn}(r_x \widehat{v}_{L,y} - r_y \widehat{v}_{L,x})$, the directions were

$$\begin{aligned} \widehat{\psi}_{\text{current}} &= \mathcal{B}(\mathbf{r}), \\ \widehat{\psi}_{\text{angular}} &= \mathcal{B}(\mathbf{r}) + sa, \\ \widehat{\psi}_{\text{CV}} &= \mathcal{B}(\mathbf{r} + \tau \widehat{\mathbf{v}}_L), \\ \widehat{\psi}_{\text{spatial}} &= \mathcal{B}\left(\mathbf{r} + d \frac{\widehat{\mathbf{v}}_L}{\|\widehat{\mathbf{v}}_L\|}\right). \end{aligned} \tag{S83}$$

The angular-lead sign was set to zero when the cross-product magnitude was below $10^{-12}\|\mathbf{r}\|\|\widehat{\mathbf{v}}_L\|$, a numerical tolerance for collinear vectors. Shared evaluation frames required finite head and torso yaw and target velocity, horizontal range above $10^{-4}$ m and target speed above $10^{-6}$ m s$^{-1}$ in the nominal reconstruction and every joint draw. These safeguards ensure a defined bearing and travel direction.

#### Leave-one-run-out parameter selection and scoring

Each lead model had one parameter $p$, selected from its own grid:

$$\begin{aligned} \mathcal{G}_{\text{angular}} &= \{0, 0.5, 1.0, \ldots, 180\}^\circ, \\ \mathcal{G}_{\text{CV}} &= \{0, 0.005, 0.010, \ldots, 0.400\}\ \text{s}, \\ \mathcal{G}_{\text{spatial}} &= \{0, 0.025, 0.050, \ldots, 5.000\}\ \text{m}. \end{aligned} \tag{S84}$$

Current-target alignment had no fitted parameter. For pursuit $j$, model $m$, and valid-frame set $\mathcal{V}_j$, the loss was

$$E_{jm}(p) = \frac{1}{|\mathcal{V}_j|} \sum_{t \in \mathcal{V}_j} \left\{ \mathrm{wrap}_{[-\pi,\pi)} [\psi_{H,j}(t) - \widehat{\psi}_{jm}(t;p)] \right\}^2. \tag{S85}$$

The parameter for held-out pursuit $h$ minimised the equally weighted training-pursuit loss:

$$p^*_{h,m} = \underset{p \in \mathcal{G}_m}{\arg\min} \frac{1}{J-1} \sum_{\substack{j=1 \\ j \neq h}}^{J} E_{jm}(p), \qquad J = 5. \tag{S86}$$

Ties selected the smallest grid value. The parameter was then applied unchanged to the withheld pursuit. Its RMSE was $(180/\pi)\sqrt{E_{hm}(p^*_{h,m})}$; the overall equal-run summary was

$$\mathrm{RMSE}_{m,\text{equal-run}} = \frac{180}{\pi}\sqrt{\frac{1}{J}\sum_{h=1}^{J} E_{hm}(p^*_{h,m})}. \tag{S87}$$

Thus, held-out mean squared errors were averaged before taking the square root.

#### Monte Carlo propagation

Velocity estimation, parameter selection and held-out scoring were repeated within each of the 500 complete joint pose–target reconstructions. Figure 5b reports medians and 2.5th–97.5th percentiles across these paired draws, with aggregate RMSE calculated within each draw before summarisation. These intervals quantify reconstruction uncertainty under the perturbations specified in Supplementary Note S5.

#### Path intersection and signed spatial lead

Spatial lead was calculated independently of CV horizon fitting. Let $\{\mathbf{P}_k\}_{k=1}^{K} \subset \mathbb{R}^3$ denote the full, time-ordered, piecewise-linear target path. Its cumulative 3D arc-length coordinate was

$$s_1 = 0, \qquad s_{k+1} = s_k + \|\mathbf{P}_{k+1} - \mathbf{P}_k\|_2 \,. \tag{S88}$$

At frame $i$, a horizontal ray was cast from the current head origin:

$$\mathbf{R}_i(\rho) = \mathbf{H}_{xy}(t_i) + \rho\mathbf{d}_i, \qquad \mathbf{d}_i = \begin{bmatrix}\cos\psi_H(t_i)\\ \sin\psi_H(t_i)\end{bmatrix}, \qquad \rho \geq 0. \tag{S89}$$

For target-path segment $k$, define

$$\mathbf{A}_k = \mathbf{P}_{k,xy}, \qquad \mathbf{e}_k = \mathbf{P}_{k+1,xy} - \mathbf{P}_{k,xy}, \qquad \mathbf{q}_{ik} = \mathbf{A}_k - \mathbf{H}_{xy}(t_i). \tag{S90}$$

For horizontal vectors, let $\mathbf{a} \times_2 \mathbf{b} = a_x b_y - a_y b_x$. When $D_{ik} = \mathbf{d}_i \times_2 \mathbf{e}_k \neq 0$, the ray and segment intersection parameters were

$$\rho_{ik} = \frac{\mathbf{q}_{ik} \times_2 \mathbf{e}_k}{D_{ik}}, \qquad \alpha_{ik} = \frac{\mathbf{q}_{ik} \times_2 \mathbf{d}_i}{D_{ik}}. \tag{S91}$$

A candidate intersection was retained when $\rho_{ik} \geq 0$ and $0 \leq \alpha_{ik} \leq 1$, subject to numerical tolerance. Parallel non-collinear segments were rejected. For a collinear segment, the nearest point on the forward ray was retained.

The 3D intersection and its path coordinate were interpolated using the same segment parameter:

$$\mathbf{P}^{\text{hit}}_{ik} = \mathbf{P}_k + \alpha_{ik}\left(\mathbf{P}_{k+1} - \mathbf{P}_k\right), \tag{S92}$$

$$s^{\text{hit}}_{ik} = s_k + \alpha_{ik}(s_{k+1} - s_k). \tag{S93}$$

When the head-forward ray intersected the path more than once, the intersection with the smallest forward ray distance $\rho_{ik}$ was selected. Ties were resolved by choosing the candidate whose path coordinate was closest to the target's current path coordinate. An intersection was first sought on the recorded path. If none existed, the five lateral-pursuit paths were continued forwards from their final recorded position, assuming that the lure maintained its terminal horizontal travel direction at constant height. This direction was estimated by a linear least-squares fit to horizontal target positions over the final 100 ms (using at least three observations). The extension was unbounded; only intersections forward of both the head and the target-path endpoint were accepted. Their additional horizontal distance was added to the recorded 3D path arc length. Frames without either an observed-path or extended-path intersection remained undefined. These extensions are descriptive assumptions about continuation beyond the recording, rather than observed future motion.

If $s_L(t_i)$ denotes the current target path coordinate, signed spatial lead was

$$d(t_i) = s^{\text{hit}}(t_i) - s_L(t_i). \tag{S94}$$

Positive values indicated intersections ahead of the current target along the time-oriented path, and negative values indicated intersections behind it. Normalised spatial lead was

$$\eta(t_i) = \frac{d(t_i)}{R(t_i)}, \qquad R(t_i) = \|\mathbf{L}_{3D}(t_i) - \mathbf{H}_{3D}(t_i)\|_2 \,. \tag{S95}$$

Thus, ray–path intersection was solved in the horizontal plane, whereas both the cumulative path coordinate and normalising head–target distance used 3D reconstructed geometry. All panels of figure S15 include endpoint continuation where required; panel (a) summarises the complete analysis windows, independently of the display crop in panel (c).

For a straight target path in the horizontal plane, the sine rule gives

$$\frac{d_h}{R_h} = \frac{\sin\gamma_h}{\sin\chi}, \tag{S96}$$

where $R_h$ is horizontal head–target range, $d_h$ is signed distance ahead along the straight horizontal target path, $\gamma_h$ is the signed angle from target bearing to head direction, and $\chi$ is the signed angle from head direction to target travel, using the same rotational convention. This relation requires a valid forward intersection and $\sin\chi \neq 0$. It makes proportional scaling with range conditional on fixed angular geometry. These planar quantities are distinct from the 3D path length and range used for the reported $d/R$ summaries.

#### Reconstruction uncertainty in spatial lead

Target reconstruction draws were available over analysed frames, whereas intersection required the complete target route. For each draw, the draw-specific displacement from the nominal target reconstruction was linearly interpolated in frame index onto the full nominal path. The nearest endpoint displacement was held constant outside the reconstructed analysis interval, and the draw-specific target positions were imposed exactly at analysed frames. Path arc length and all intersections were then recalculated from the resulting draw-specific full path. The terminal travel direction and any required endpoint extension were also recalculated within each draw, propagating reconstruction uncertainty under the continuation assumption.

Framewise spatial-lead percentiles were calculated across draws having a valid intersection. Intersection support was recorded separately as the fraction of all draws yielding an intersection at that frame. For the normalised summary, $d/R$ was calculated within each complete draw, followed by the median across eligible frames in each pursuit. Medians and 2.5th–97.5th percentiles of these draw-specific pursuit medians were then reported. For this spatial summary, Phantom Run used the available frames 100–212 within the predefined 100–216 window to omit the earlier near-stationary target segment. The candidate-model comparison used its full retained analysis window.

### Supplementary Tables

Table S1: **Final behavioural cohort and primary analysis subsets.** The final cohort comprised 1,708 matched, quality-retained frames from 18 pursuits involving 6 cheetahs. Frontal, lateral and close pass-by pursuits partition this cohort; the analysis subsets below overlap and must not be summed. Monte Carlo reconstructions are uncertainty samples, not additional biological observations.

| Cohort or analysis | Frames | Pursuits | Cheetahs | Use |
|---|---|---|---|---|
| Full behavioural cohort | 1,708 | 18 | 6 | Yaw geometry and tracking-state classification |
| Frontal pursuits | 1,080 | 11 | 5 | Frontal-context yaw summaries; included in pitch analyses |
| Lateral pursuits | 510 | 5 | 3 | Lateral-context yaw summaries, signed yaw-error distributions and candidate-model comparison; included in pitch analyses |
| Close pass-by pursuits | 118 | 2 | 1 | Both Zorro pursuits; retained for yaw analyses and excluded from pitch analyses |
| Pitch alignment and precision | 1,590 | 16 | 5 | Frontal and lateral pursuits |
| Pitch counter-rotation | 1,590 | 16 | 5 | 1,504 overlapping 50-ms change intervals derived from the retained frames |
| Absolute yaw-error ECDFs | 628 | 7 | 4 | Lateral and close pass-by pursuits |

Table S2: **Local coordinates of the head and torso template landmarks.** Model dimensions were sourced from the AcinoSet study [S1], except for the ear dimensions, which were estimated by multiview triangulation in the present study. Unit is in meters.

| Landmark | $x$ | $y$ | $z$ |
|---|---|---|---|
| Left eye | 0 | 0.0300 | 0 |
| Right eye | 0 | −0.0300 | 0 |
| Nose | 0.0550 | 0 | −0.0550 |
| Left ear | −0.0725 | 0.0608 | 0.0441 |
| Right ear | −0.0725 | −0.0608 | 0.0441 |
| Neck base | 0.0400 | 0 | 0.1000 |
| Spine | −0.3300 | 0 | 0.1000 |
| *Reference landmarks* | | | |
| Left ear top | −0.0725 | 0.0608 | 0.0780 |
| Right ear top | −0.0725 | −0.0608 | 0.0780 |
| Left ear inner | −0.0725 | 0.0200 | 0.0441 |
| Right ear inner | −0.0725 | −0.0200 | 0.0441 |
| Left lower head | −0.0650 | 0.0608 | −0.0500 |
| Right lower head | −0.0650 | −0.0608 | −0.0500 |
| Left front shoulder | 0.0400 | 0.1200 | 0.0150 |
| Right front shoulder | 0.0400 | −0.1200 | 0.0150 |

Table S3: **Camera self-calibration corrections and final frame offsets.** The table includes the 18 pursuits in the quantitative behavioural cohort and two turning sequences (`julesBottomFlick21_20171221` and `phantomBottomFlick2_20171209`) retained as supplementary context.

| Sequence | Final frame offset (manual) Cam 1 | Cam 2 | Cam 3 | Cam 4 | Cam 5 | Cam 6 | Max. $\lvert\phi_c\rvert$ (frame) | Max. rotation (deg) | Max. $\lVert\Delta \mathbf{C}_c\rVert$ (cm) | Held-out error (px) Original | Final |
|---|---|---|---|---|---|---|---|---|---|---|---|
| `cetaneTopRun11_20171212` | −16 | −1 | 0 | +1 | 0 (−1) | −1 | 0.46 | 0.9 | 7.9 | 4.8 | 3.5 |
| `cetaneTopRun1_20171216` | N/A | −1 (0) | 0 | −1 (0) | +1 (0) | +1 (0) | 0.43 | 0.3 | 2.2 | 15.0 | 5.9 |
| `julesBottomFlick2_20171209` | 0 | 0 | 0 (+1) | 0 | 0 | N/A | 0.18 | 0.4 | 6.1 | 9.9 | 4.4 |
| `julesTopRun1_20170902`† | 0 | +1 | 0 | +1 (0) | +1 (0) | +1 (0) | 0.37 | – | – | 6.5 | 4.6 |
| `lilyRun_20190305` | −2 | −2 (−1) | −2 (0) | −1 (0) | −3 (0) | −3 (0) | 0.44 | 1.7 | 9.3 | 19.6 | 6.7 |
| `lilyTopRun1_20171221` | 0 | −1 | 0 | −2 (0) | +1 (0) | 0 | 0.41 | 2.2 | 7.2 | 28.6 | 3.5 |
| `menyaRun_20190307` | 0 | +1 | +1 | +1 (0) | +1 | −1 | 0.39 | 1.4 | 13.5 | 10.4 | 5.1 |
| `phantomRun_20190307` | −1 (0) | 0 | −1 (0) | −1 (0) | 0 | −2 (0) | 0.19 | 1.2 | 11.0 | 9.4 | 3.7 |
| `phantomTopRun11_20170902` | 0 | +2 (+1) | 0 | 0 | 0 | 0 (+1) | 0.44 | 1.0 | 6.2 | 6.5 | 3.7 |
| `phantomTopRun13_20170902` | 0 | +2 | 0 | 0 | 0 | 0 | 0.20 | 1.2 | 3.8 | 5.6 | 2.6 |
| `phantomTopRun1_20170903`‡ | 0 | +3 | 0 | 0 | −1 | 0 (+1) | 0.37 | 0.7 | 7.1 | 4.6 | 3.1 |
| `julesFlick1_20190309` | 0 | 0 | +1 (−1) | +1 (0) | −1 | 0 (−2) | 0.37 | 1.3 | 9.4 | 9.1 | 5.0 |
| `julesTopRun11_20170829` | 0 | 0 | 0 | 0 | 0 | 0 | 0.31 | 1.1 | 5.7 | 8.2 | 4.8 |
| `julesTopRun1_20171209` | +1 (0) | +2 (+1) | +6 | +3 (+2) | +3 | N/A | 0.38 | 0.5 | 4.3 | 6.1 | 2.8 |
| `lilyRun_20190309` | 0 | 0 | +1 (0) | 0 | 0 | 0 | 0.28 | 1.0 | 10.7 | 14.5 | 6.1 |
| `phantomRun_20190303` | 0 | 0 | −1 (0) | N/A | N/A | +1 (0) | 0.42 | 0.7 | 5.3 | 11.3 | 3.2 |
| `zorroBottomFlick2_20170829` | 0 | +2 | 0 | +1 (0) | +2 | +1 | 0.45 | 0.9 | 6.1 | 6.4 | 3.6 |
| `zorroBottomFlick2_20170903`† | +2 (0) | +1 | +1 | +1 | +1 | +2 | 0.38 | – | – | 12.0 | 5.4 |
| `julesBottomFlick21_20171221`‡ | +2 (0) | +1 | +1 | +1 | +2 | +1 | 0.43 | 0.4 | 2.8 | 6.1 | 4.1 |
| `phantomBottomFlick2_20171209` | 0 | 0 | 0 | N/A | +1 (0) | N/A | 0.46 | 0.5 | 1.8 | 7.2 | 5.1 |

Offsets are the final integer offsets; the manual offset is given in parentheses where it changed. $\phi_c$ is the sub-frame remainder removed from the labels. Rotation and $\lVert\Delta \mathbf{C}_c\rVert$ are the largest extrinsic corrections over the cameras of each sequence. Held-out error is the median held-out label error in raw pixels under four-fold temporal-block cross-validation for the original geometry and for the selected geometry. N/A, camera not available. †Timing correction only (extrinsics unchanged). ‡One camera retained its original extrinsics.

Table S4: **Trajectory-reconstruction reprojection quality for the 18 analysed sequences.** The cohort comprises 11 pitch-tracking and 7 yaw-tracking sequences, reconstructed after self-calibration. Statistics use the final 1,708 matched, quality-retained frames. RMSE values are in undistorted image pixels; bracketed intervals give the IQR (Q1–Q3). Across all valid frames, the frame-weighted mean per-frame RMSE was 5.13 pixels. The retained-observation column gives the percentage of visible landmark observations surviving camera-trust filtering.

| Dataset | Valid frames | Median RMSE [IQR] (px) | Mean RMSE (px) | 95th pct. RMSE (px) | Retained obs. |
|---|---|---|---|---|---|
| `cetaneTopRun11_20171212` | 117 | 3.78 [3.03, 5.26] | 4.48 | 9.72 | 95.6% |
| `cetaneTopRun1_20171216` | 38 | 3.75 [3.20, 8.98] | 5.59 | 10.78 | 92.1% |
| `julesBottomFlick2_20171209` | 43 | 3.61 [3.24, 4.22] | 4.22 | 9.69 | 90.3% |
| `julesTopRun1_20170902` | 72 | 4.63 [3.73, 5.97] | 5.62 | 12.68 | 93.6% |
| `lilyRun_20190305` | 137 | 5.48 [4.78, 6.52] | 6.20 | 11.37 | 94.5% |
| `lilyTopRun1_20171221` | 63 | 3.98 [3.57, 4.67] | 4.24 | 6.32 | 98.2% |
| `menyaRun_20190307` | 86 | 7.83 [6.81, 8.79] | 7.77 | 11.07 | 95.5% |
| `phantomRun_20190307` | 185 | 4.53 [3.89, 7.46] | 5.62 | 10.92 | 98.3% |
| `phantomTopRun11_20170902` | 126 | 3.87 [2.66, 4.54] | 4.03 | 8.24 | 97.9% |
| `phantomTopRun13_20170902` | 78 | 3.13 [2.84, 3.69] | 4.06 | 10.10 | 98.2% |
| `phantomTopRun1_20170903` | 135 | 3.57 [2.82, 4.64] | 3.84 | 7.75 | 99.4% |
| `julesFlick1_20190309` | 78 | 4.29 [3.93, 5.89] | 4.97 | 9.04 | 95.6% |
| `julesTopRun11_20170829` | 70 | 4.85 [3.98, 6.02] | 5.75 | 12.41 | 92.3% |
| `julesTopRun1_20171209` | 77 | 2.56 [2.26, 3.31] | 3.03 | 5.67 | 97.7% |
| `lilyRun_20190309` | 96 | 4.17 [3.45, 5.70] | 5.20 | 10.53 | 88.4% |
| `phantomRun_20190303` | 189 | 3.31 [2.70, 3.87] | 3.51 | 6.39 | 94.4% |
| `zorroBottomFlick2_20170829` | 35 | 8.57 [5.37, 9.83] | 7.86 | 11.42 | 100.0% |
| `zorroBottomFlick2_20170903` | 83 | 8.23 [5.58, 12.74] | 9.84 | 19.56 | 96.5% |

Table S5: **Image-plane perturbation scales used in the Monte Carlo reconstruction.** The radial scale is the RMS magnitude of the 2D displacement.

| Landmark | Radial RMS $s_j$ (pixels) | Per-coordinate SD $s_j/\sqrt{2}$ (pixels) |
|---|---|---|
| Left eye | 2.5 | 1.768 |
| Right eye | 2.5 | 1.768 |
| Nose | 2.5 | 1.768 |
| Left ear | 3.0 | 2.121 |
| Right ear | 3.0 | 2.121 |
| Spine | 3.5 | 2.475 |
| Neck base | 3.5 | 2.475 |
| Lure | 2.828 | 2.000 |

Table S6: **Template-geometry perturbations used in the Monte Carlo reconstruction.** Each dimension was drawn independently from $\mathcal{N}(L_0, \sigma_L^2)$ once per Monte Carlo sample and held fixed across all frames. Here, $L_0$ is the template dimension multiplied by the dataset-specific head or torso scale. Left–right symmetry was preserved.

| **Template dimension** | **SD $\sigma_L$** |
|---|---|
| Ear position, $x$ | 11 mm |
| Ear position, $y$ | 5 mm |
| Ear position, $z$ | 9 mm |
| All remaining dimensions (eye $y$; nose, neck, and spine $x$, $z$) | $0.08\lvert L_0 \rvert$ |

Table S7: **Run-level counter-rotation and target-elevation coefficients.** Coefficients were estimated for 16 frontal and lateral pursuits (1,590 frames from 5 cheetahs), using 1,504 nominal 50-ms change intervals. The model included concurrent torso-pitch change, $\Delta\theta_T$, and head–target-line elevation change, $\Delta\lambda_H$. The number of change intervals contributing to each fit is denoted by $n_{\text{pairs}}$. A torso-disturbance gain of $g_D = 1$ corresponds to complete kinematic cancellation of the torso-pitch contribution to world-frame head motion.

| **Animal** | **Pursuit sequence** | $n_{\text{pairs}}$ | $g_D$ | $g_\lambda$ |
|---|---|---|---|---|
| Cetane | `cetaneTopRun11_20171212` | 112 | 1.077 | −2.253 |
| Cetane | `cetaneTopRun1_20171216` | 33 | 0.990 | 1.406 |
| Jules | `julesBottomFlick2_20171209` | 38 | 0.834 | 1.218 |
| Jules | `julesTopRun1_20170902` | 67 | 1.009 | −4.765 |
| Lily | `lilyRun_20190305` | 131 | 1.091 | −1.203 |
| Lily | `lilyTopRun1_20171221` | 58 | 1.385 | −3.755 |
| Menya | `menyaRun_20190307` | 80 | 1.095 | −1.527 |
| Phantom | `phantomRun_20190307` | 179 | 0.997 | −0.259 |
| Phantom | `phantomTopRun11_20170902` | 121 | 1.013 | −0.808 |
| Phantom | `phantomTopRun13_20170902` | 73 | 1.185 | −1.491 |
| Phantom | `phantomTopRun1_20170903` | 130 | 0.981 | −1.512 |
| Jules | `julesFlick1_20190309` | 72 | 1.172 | −1.901 |
| Jules | `julesTopRun11_20170829` | 65 | 1.182 | −1.120 |
| Jules | `julesTopRun1_20171209` | 72 | 1.093 | −0.216 |
| Lily | `lilyRun_20190309` | 90 | 1.181 | −3.002 |
| Phantom | `phantomRun_20190303` | 183 | 1.026 | 0.595 |

#### Supplementary Videos

**Supplementary movie S1** Overview of CRBR reconstruction method.

**Supplementary movie S2** Example 2D pose estimation for multi-view motion capture data.

**Supplementary movie S3** Example reconstruction of cheetah head and torso while chasing the lure. (Top left: reconstructed 3D scene. Top right: original camera view. Bottom: Top and side view of the reconstructed 3D scene.)
